# A red-emitting carborhodamine for monitoring and measuring membrane potential

**DOI:** 10.1101/2023.10.06.561080

**Authors:** Anneliese M. M. Gest, Julia R. Lazzari-Dean, Gloria Ortiz, Susanna K. Yaeger-Weiss, Steven C. Boggess, Evan W. Miller

## Abstract

Biological membrane potentials, or voltages, are a central facet of cellular life. Optical methods to visualize cellular membrane voltages with fluorescent indicators are an attractive complement to traditional electrode-based approaches, since imaging methods can be high throughput, less invasive, and provide more spatial resolution than electrodes.

Recently developed fluorescent indicators for voltage largely report changes in membrane voltage by monitoring voltage-dependent fluctuations in fluorescence intensity. However, it would be useful to be able to not only monitor changes, but also measure values of membrane potentials. This study discloses a new fluorescent indicator which can address both.

We describe the synthesis of a new sulfonated tetramethyl carborhodamine fluorophore. When this <u>c</u>arbo<u>rh</u>odamine is conjugated with an electron-rich, methoxy (-<u>OMe</u>) containing phenylenevinylene molecular wire, the resulting molecule, CRhOMe, is a voltage-sensitive fluorophore with red/far-red fluorescence.

Using CRhOMe, changes in cellular membrane potential can be read out using fluorescence intensity or lifetime. In fluorescence intensity mode, CRhOMe tracks fast-spiking neuronal action potentials with greater signal-to-noise than state-of-the-art BeRST (another voltage-sensitive fluorophore). CRhOMe can also measure values of membrane potential. The fluorescence lifetime of CRhOMe follows a single exponential decay, substantially improving the quantification of membrane potential values using fluorescence lifetime imaging microscopy (FLIM). The combination of red-shifted excitation and emission, mono-exponential decay, and high voltage sensitivity enable fast FLIM recording of action potentials in cardiomyocytes. The ability to both monitor and measure membrane potentials with red light using CRhOMe makes it an important approach for studying biological voltages.

**Significance Statement:** Biological membrane potentials are maintained by all forms of life. In electrically excitable cells, fast changes in membrane potential drive downstream events: neurotransmitter release, contraction, or insulin secretion. The ability to monitor changes in and measure values of cellular membrane potentials is central to a mechanistic understanding of cellular physiology and disease. Traditional modes for measuring membrane potential use electrodes, which are invasive, destructive, low throughput, and ill-suited to interrogate spatial dynamics of membrane potentials. Optical methods to visualize potentials with fluorescent dyes offer a powerful complement to traditional electrode approaches. In this study, we show that a new, red to farred fluorophore can both monitor changes in and measure values of membrane potential in a variety of living systems.

## Introduction

Membrane potential plays central roles in the physiology of all living systems. Rapid changes in membrane potential initiate neurotransmitter release in neurons,a prompt muscle contraction in cardiomyocytes,^2-3^ and evoke insulin secretion in pancreatic β cells.^4^ In a related fashion, the value of the membrane potential – set largely by the difference in internal and external potassium ion concentration and the selective permeability of the plasma membrane to K^+^ ions – plays an outsized role in the physiology of these cell types, controlling their excitability by setting the distance to action threshold. Further, values of membrane potentials in non-electrically excitable cells may play important roles in signaling, differentiation, and cell cycle progression.^5-7^ Reflecting the importance of membrane potential, some 10 to 50% of oxygen consumption is directed towards setting the “resting” membrane potential through the action of the ATP-driven Na-K exchanger.^8^

The primary mode of monitoring changes in and measuring values of membrane potential is patch-clamp electrophysiology. Despite the centrality of membrane potential to cellular physiology across all organ systems, electrophysiology is nonetheless invasive, low throughput, and difficult to implement across spatial scales. Electrophysiological analysis of multiple cells requires highly specialized instrumentation,^9-10^ and interrogation of subcellular or intracellular membranes remain largely inaccessible except in limited cases.^11-13^

To complement traditional electrode-based methods, we have been pursuing voltage-sensitive fluorophores (VoltageFluors, or VF dyes) that sense voltage via a photoinduced electron transfer (PeT)-based mechanism.^14-15^ This approach can be adapted to a wide range of chemically-synthesized fluorophores.^16^ Dyes with excitation and emission in the red region of the spectrum are especially useful, since they display lower levels of phototoxicity and minimize autofluorescence by avoiding excitation of endogenous chromophores;^17^ have higher photostability;^17-18^are compatible with optical indicators and actuators that use blue or green light; are well matched to commercially available powerful LEDs; and pair well with the wavelength-dependent quantum efficiency of typical sCMOS detectors, which falls off dramatically beyond 650 nm.^19-20^ These characteristics make VF dyes well suited for tracking fast membrane potential changes associated with action potentials in neurons (AP duration ∼2 ms) or cardiomyocytes (AP duration ∼100 ms).

We previously showed that VF dyes can be used to estimate values of membrane potential.^21^ Using fluorescence lifetime imaging microscopy (FLIM) to measure the fluorescence lifetime of VF dyes in cell membranes, we can estimate millivolt values of membrane potential, since fluorescence lifetime is more resistant than fluorescence intensity to variations in dye loading, fluctuations in excitation power, or bleaching.^22^ Our initial efforts with fluorescein-based VF2.1.Cl showed that FLIM enables membrane potential value estimation with up to 19× improved resolution compared to existing methods.^21^ However, the long acquisition times required for FLIM, coupled with the blue-light excitation of VF2.1.Cl means that even low levels of phototoxicity becomes limiting in sensitive systems (like cardiomyocytes or neurons).^23^ Further, VF2.1.Cl has a two-component fluorescence lifetime decay, complicating analysis and requiring longer photon collection times.^24-25^

Here, we describe a new VF dye based on a tetramethyl carborhodamine^26^ scaffold. The new carborhodamine indicators have red excitation and emission profiles and show high voltage sensitivity. One indicator in particular, a <u>C</u>arbo <u>Rh</u>odamine with a methoxy (<u>OMe</u>) wire, or CRhOMe (“chrome”), can detect rapid changes of membrane potential in hippocampal neurons with improved signal-to-noise ratio (SNR) compared to BeRST, our previous best in class. CRhOMe also possesses a long, monoexponential fluorescence lifetime, dramatically improving its performance for tracking membrane potential values and changes using FLIM. The unique photophysical properties of CRhOMe – high voltage sensitivity, long, monoexponential fluorescence lifetime (∼3 ns), emission above 640 nm, and excellent photostability – substantially improve its performance compared to VF2.1.Cl in FLIM, making CRhOMe an exceptionally promising candidate for measuring membrane potential values and changes with FLIM.

### Synthesis

We synthesized three sulfonated carborhodamine fluorophores (**7, 8**, and **9, Scheme 1**) and five carborhodamine voltage indicators (**13-17, Scheme 1**). The synthesis of sulfonated carborhodamine fluorophores begins with the triflation of anthracenone **1**^26^ with trifluoromethanesulfonic anhydride and pyridine in dichloromethane to give **2** in 50% yield (**Scheme 1**). Triflate **2** is then subjected to a Buchwald-Hartwig reaction with dimethylamine to yield **3** in 96% yield. This cross-coupling route was previously established to install amino groups on fluorescein derivatives to access rhodamines.^26-28^ To install the *meso* aromatic ring bearing a sulfonate group, we followed conditions used in the synthesis of BeRST 1.^18^ Compound **3** is triflated to increase reactivity, then reacted with the organolithiums derived from **4, 5**, or **6** with *n*-butyllithium in dichloromethane at -16 °C to yield sulfonated carborhodamines **7**-**9** in 13-17% yields (**Scheme 1**). Sulfonated carborhodamines **8** and **9** are then subjected to a Heck reaction with styrenes **10, 11**, or **12** to yield tetra<u>m</u>ethyl <u>c</u>arbo<u>rh</u>odamine (TMCRh) voltage indicators **14**-**17** and **13** (R^3^ = R^4^ = H), which lacks the alkyl-substituted aniline required for voltage sensitivity.^29^

### Spectroscopic Characterization

We examined the photophysical properties of the tetramethyl carborhodamine fluorophore **7** and dyes **13**-**17**. Sulfonated TMCRh **7** displays a λ_max_ of 610 nm in both MeOH (ε = 129,000 M^-1^ cm^-1^) and EtOH with 0.1% TFA (ε = 125,000 M^-1^ cm^-1^) and an 11-nm shift in aqueous buffer with a λ_max_ of 621 nm (ε = 129,000 M^-1^ cm^-1^) (**Figure S1**). In alcoholic solvents the λ_em_ is 633 nm, and in aqueous buffer the λ_em_ is 641 nm. Sulfonation of TMCRh shifts absorbance and emission to the blue by ∼16 nm compared to carboxylated tetramethylcarborhodamine.^28^ We observed a similar bathochromic shift for the comparison between sulfonated carbofluoresceins^30^ and carboxy-substituted carbofluoresceins.^26^

The TMCRh voltage indicator derivatives **13**-**17** display similar spectral profiles in aqueous buffer with a λ_max_ of 622 nm and a λ_em_ of 641 nm (**Table 1, Figure 1** and **Figure S1**). Fluorescence quantum yields (Φ) for putative voltage reporters **14-17** are low (0.038 to 0.096), while the voltage insensitive TMCRhZero (**13**) is much higher (Φ = 0.62), comparable to the TMCRh fluorophore **7** alone (Φ = 0.71). The 6- to 18-fold decrease in Φ for **14**-**17** compared to **13** indicates a high degree of PeT from the aniline donor to the fluorophore.

**Table 1.** Photophysical properties of carborhodamine dyes. ^*a*^ Determined in PBS, pH 7.4, 0.1% SDS. ^*b*^ Determined in MeOH. ^*c*^ Voltage-clamped HEK cells. Error is ± S.D. for n = 4-7 cells. ^*d*^ Determined in HEK cells. Error is ± S.E.M for n = 4 coverslips (>100 cells per coverslip) for relative brightness.

| Compound | R<sup>3</sup> | R<sup>4</sup> | λ<sub>max</sub> / nm<sup>a</sup> | λ<sub>em</sub> / nm<sup>a</sup> | Φ<sup>a/b</sup> | ΔF/F / 100 mV<sup>c</sup> | Relative brightness<sup>d</sup> |
| --- | --- | --- | --- | --- | --- | --- | --- |
| TMCRh-sulfonate **7** | n/a | n/a | 621 | 641 | 0.71<sup>a</sup> | n/a | n/a |
| TMCRhZero **13** | H | H | 622 | 641 | 0.62/0.61 | 0.3% ± 0.1% | 3.9 ± 0.12 |
| TMCRh.H **14** | H | NMe<sub>2</sub> | 622 | 642 | 0.096/0.087 | 1.2% ± 0.4% | 1.0 ± 0.04 |
| CRhOMe **15** | OMe | NMe<sub>2</sub> | 622 | 642 | 0.038/0.027 | 18% ± 2% | 3.8 ± 0.13 |
| isoTMCRh.H **16** | H | NMe<sub>2</sub> | 621 | 641 | 0.050/0.057 | 1.5% ± 0.2% | 5.4 ± 0.22 |
| isoCRhOMe **17** | OMe | NMe<sub>2</sub> | 621 | 641 | 0.058/0.054 | 9% ± 2% | 17 ± 0.84 |

**Figure 1.**
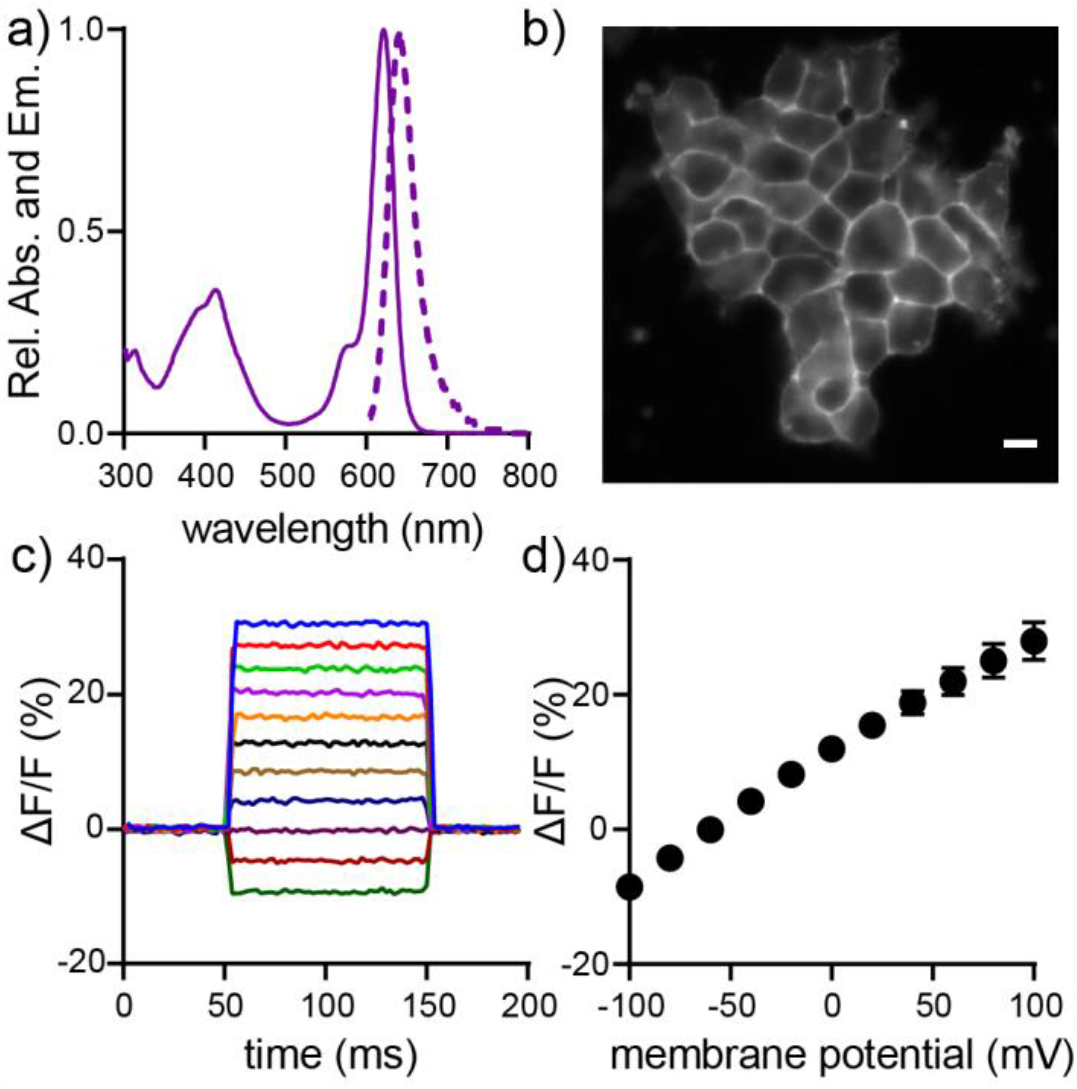
Characterization of CRhOMe (15) voltage sensitivity in HEK293T. **a)** Absorption (solid line) and emission (dashed line) spectra of CRhOMe (**15**) in PBS, pH 7.4, 0.1% SDS. **b)** Epifluorescence image of a group of HEK293T cells stained with 500 nM CRhOMe. Scale bar is 10 μm. **c)** Change in fluorescence for a single HEK293T cell under whole cell voltage clamp conditions in which the membrane potential was stepped from +100 to -100 mV in 20 mV increments. **d)** Average fluorescence intensity change (%ΔF/F) observed across multiple voltage-clamped HEK293T cells. Error bars are ± standard deviation for n = 8 cells.

### Voltage Sensitivity in HEK cells

All aniline-containing voltage reporter derivatives (**14**-**17**) stain cellular membranes and show V_mem_ sensitivity in HEK293T cells (**Figure 1c,d**, **Figure S1-2**). TMCRh voltage indicators with the unsubstituted wire (**14, 16**, R^3^ = H) display lower voltage sensitivity than their corresponding methoxy-substituted wire counterparts (**15, 17**, R^3^ = OMe). Compound **15** has a voltage sensitivity of 18% ± 2% ΔF/F per 100 mV and its isomer **17** has a voltage sensitivity of 9% ± 2% ΔF/F per 100 mV. We observe the highest voltage sensitivity in **15** and no voltage sensitivity in TMCRhZero **13** (0.3 ± 0.1%). These trends are similar to V_mem_ sensitivity patterns seen with other VoltageFluors, where we hypothesize that the added electron density of the methoxy-substituted molecular wire increases the dynamic range of PeT accessible to the molecule.^31^ Because of its superior ΔF/F, we selected compound **15**, a <u>C</u>arbo <u>Rh</u>odamine with a methoxy (<u>OMe</u>) wire, or CRhOMe (“chrome”), for further investigation in subsequent experiments, including fluorescence lifetime imaging studies, along with TMCRhZero as a voltage-insensitive control compound.

### Performance in Neurons

CRhOMe **15** stains cell membranes of dissociated rat hippocampal neurons and reports spontaneous action potentials from multiple cells (**Figure 2**). Field stimulation of neurons labeled with CRhOMe reveals that CRhOMe responds to evoked action potentials with a voltage sensitivity of 8.2% ± 1.7 (1.85 W/cm^-2^; SNR = 22 ± 2, *n* = 22 cells, **Figure S3**). When comparing **15** to BeRST 1 using minimal light power (0.93 W/cm^-2^), CRhOMe reported evoked action potentials with 8.4% ± 0.3% and SNR = 13 ± 0.4 (*n* = 50 cells), comparable or better than BeRST 1, which responded to evoked action potentials with 11% ± 0.7% and SNR = 9 ± 0.5 (*n* = 21 cells, **Figure 2, S4**). At all light powers examined, CRhOMe has significantly higher SNR than BeRST (p = 0.024, paired t-test) and a trend towards a lower nominal ΔF/F (9% vs 11%, p = 0.068, paired t-test, **Figure 2, S4**). CRhOMe shows excellent resistance to photobleaching, comparable to BeRST, the existing best-in-class (**Figure S4**).^18, 32^ At higher light powers, around 13 W/cm^2^, or approximately 10× higher than that used for routine neuronal imaging in our lab,^33^ both BeRST and CRhOMe display the previously reported phototoxicity associated with extended illumination (**Figure S4g-h**).

**Figure 2.**
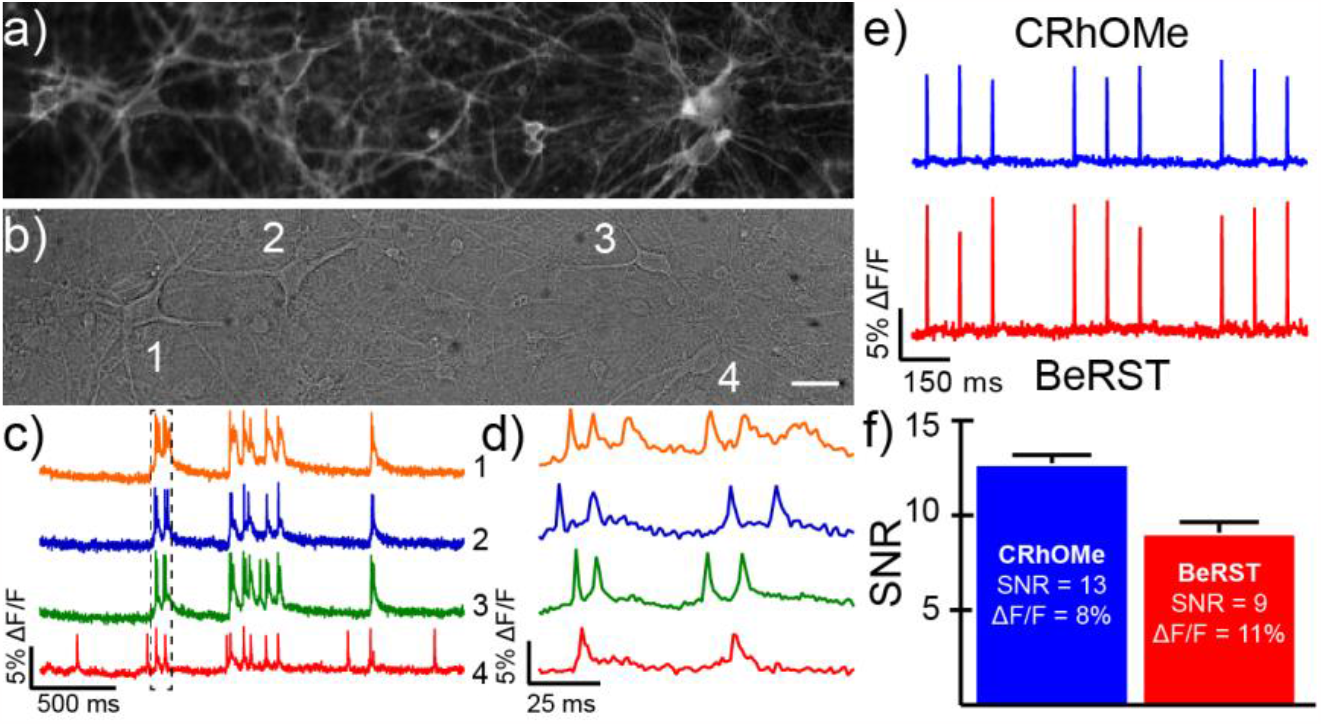
CRhOMe tracks spontaneous and evoked action potentials in cultured neurons. **a**) Wide-field microscopy fluorescence and **b)** DIC images of cultured rat hippocampal neurons stained with CRhOMe (500 nm, 30 min). Scale bar is 20 μm. **c)** Optical traces of spontaneous activity of the neurons in panels a-b) recorded at 500 Hz and shown as ΔF/F vs time. **d)** Highlighted action potentials from panel c). **e)** Plots of ΔF/F vs time for neurons stained with either CRhOMe (blue) or BeRST 1 (red) and then subjected to field stimulation to evoke action potential responses. **f)** Comparison of signal to noise ratio (SNR) for CRhOMe (blue) and BeRST (red) in neurons stimulated as in panel e. Data represent mean ± S.E.M. for n = 50 or 21 neurons for CRhOMe and BeRST, respectively.

### Lifetime Studies in Cells

We recorded the time resolved fluorescence decay of the TMCRh fluorophore **7** (1 μM in water) with time correlated single photon counting (TCSPC) FLIM on a point scanning confocal microscope. The time-resolved fluorescence decay was well described by a single exponential model and exhibited a lifetime of 3.06 ± 0.02 ns (mean ± SEM of 16 measurements, **Figure S5**). We then measured the fluorescence lifetime of CRhOMe in living cells (**Figure 3a**), which was also well described by a single exponential decay. We observe a mean lifetime of 2.57 ± 0.01 ns in HEK293T at rest (mean ± SEM of 132 cell groups) and a mean lifetime of 2.60 ± 0.02 ns in serum-starved A431 cells (mean ± SEM of 18 cell groups). For the voltage-insensitive control compound TMCRhZero in HEK293T (**Figure S7**), we again observe a single exponential decay with a lifetime of 3.70 ± 0.01 (mean ± SEM of 103 cell groups). Lifetime data collected with different instruments or at different excitation wavelengths give the same resting lifetime potentials in HEK293T cells (**Figure S6**).

**Figure 3.**
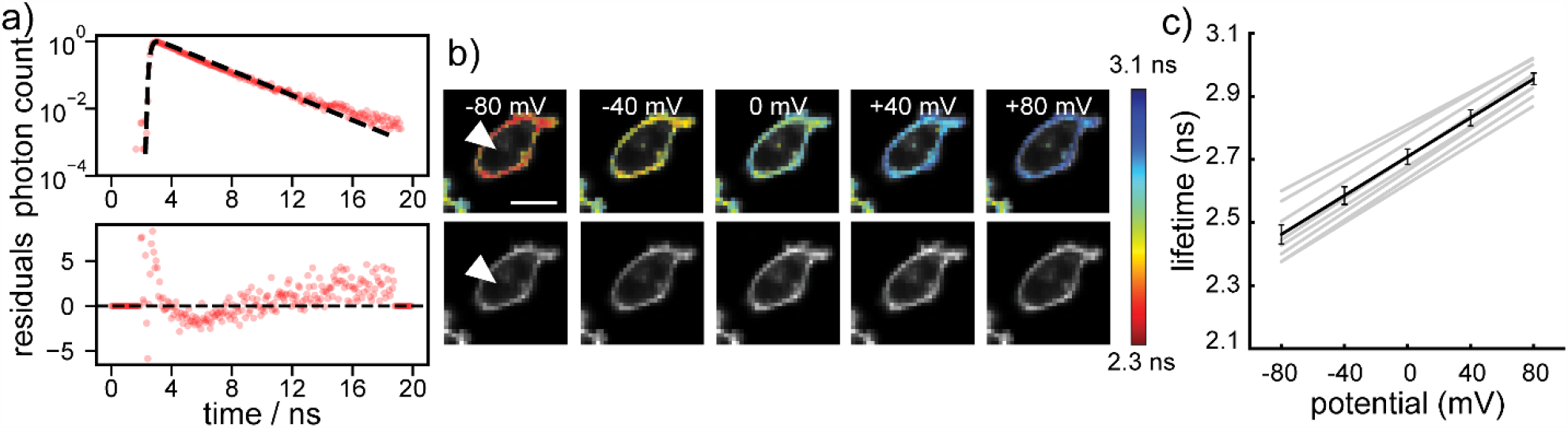
CRhOMe displays single-exponential fluorescence lifetime decays and a linear fluorescence lifetime-voltage relationship. **a)** Plots of fluorescence lifetime decay expressed as normalized photon counts (log scale) vs time for CRhOMe). Circles indicate photon count data; dashed line is a single exponential decay. Weighted residuals are plotted in the lower graph. **b)** Lifetime (colored) and intensity (grayscale) images of HEK 293T cells voltage-clamped at the indicated potentials. The lifetime heatmap is scaled from 2.3 to 3.1 ns. Scalebar is 20 μm. Arrowhead indicates the voltage-clamped cell. **c)** Plot of CRhOMe fluorescence lifetime vs membrane potential in HEK 293T cells. Gray lines are individual cell calibrations (n = 17). The average calibration is in black. Error bars are mean ± S.E.M.

As a comparison, we also investigated the fluorescence lifetime in HEK293T cells of our existing best in class far-red VF, BeRST1. Unlike CRhOMe, BeRST1 is not well described by a single exponential decay (**Figure 3a**, second trace), and was instead fit with a two-component exponential decay. We report its lifetime as the amplitude-weighted average of the two lifetime components, as we have done for previous green VoltageFluors in FLIM.^34^ In HEK293T cells, BeRST1 exhibits a weighted average lifetime of 1.80 ± 0.01 ns (mean ± SEM of 56 cell groups). This one component decay for CRhOMe is a critical advantage in FLIM studies. Simpler, one component decay models can be confidently fit with an order of magnitude fewer photons than more complicated multi-component models.^35-36^ CRhOMe FLIM data can therefore be acquired faster and with lower phototoxicity than either BeRST1 or our previously studied green VoltageFluors.

We additionally measured lifetimes across a range of dye loading concentrations for both CRhOMe and BeRST 1 to look for a concentration-dependent decrease in τ_fl_ (**Figure S7**), which we previously observed with green VoltageFluors.^34, 37^ All subsequent experiments were conducted at concentrations below the point where concentration quenching was observed (approximately 500 nM in HEK293T culture, with most experiments done at 300 nM dye).

### Calibration of CRhOMe in cells with electrophysiology and gramicidin treatment

We generated a voltage-τ_fl_ calibration using whole-cell, patch-clamp electrophysiology and simultaneous FLIM imaging to determine the voltage dependence of CRhOMe fluorescence lifetime. HEK 293T cells were treated with 300 nM CRhOMe, and then voltage-clamped at -80, -40, 0, +40, and +80 mV, and the fluorescence lifetime recorded. Individual measurements demonstrate a linear response of lifetime to changes in voltage (**Figure 3b-c**). Consistent responses among individual measurements allows for the development of an average calibration to represent the overall expected change in lifetime (in ps) per change in voltage (in mV). In HEK 293T cells, CRhOMe exhibits a sensitivity (slope of the calibration) of 3.09 ± 0.09 ps/mV, with a lifetime at 0 mV (the y-intercept of the calibration) of 2.71 ± 0.03 ns. Applying this calibration to our measured lifetime of HEK293T cells at rest, the mean resting membrane potential across all measurements is -41.0 ± 3.5 mV (mean ± SEM), consistent with previous optical and electrophysiological reports of HEK293T resting potential values.^21, 38-40^ The voltage-insensitive control compound, TMCRhZero, does not exhibit a change in lifetime to an applied voltage (**Figure S8**).

To provide a secondary, non-electrophysiological calibration for systems where electrophysiology may be impractical, we turned to the Na^+^/K^+^ ionophore gramicidin^41^ (**Figure S9-S10**). Because it increases the permeability to both Na^+^ and K^+^, we expect that gramicidin will depolarize cellular V_mem_ to approximately 0 mV. In HEK293T cells treated with 500 ng/mL gramicidin, the fluorescence lifetime of CRhOMe rose to 2.74 ± 0.09 ns (mean ± SD of 111 cell groups), which matches the 0 mV τ_fl_ of 2.71 ns determined by patch clamp calibration. We observed a similar increase in fluorescence lifetime in serum-starved A431 cells treated with gramicidin (**Figure S9**), whereas we observe minimal change in the fluorescence lifetime of TMCRhZero in gramicidin-treated HEK293T (**Figure S10**). These data suggest that gramicidin may be appropriate for creating a reference point for calibrations in cell lines where a ground-truth patch clamping experiment is not feasible.

The magnitude of CRhOMe lifetime to changes in V_mem_ is comparable to that of VF2.1.Cl (previously reported sensitivity 3.5 ps/mV, 0 mV lifetime 1.77 ns in HEK293T).^21^ The sensitivity of VF2.1.Cl in lifetime is the highest reported for a VF to date, despite the presence of probes with higher fractional voltage sensitivity (ΔF/F).^42^ CRhOMe’s high sensitivity, combined with its attractive spectral properties and single component decay, indicated that CRhOMe would be a superior choice for FLIM quantification than the previously reported VF2.1.Cl or BeRST 1. We therefore investigated applications of CRhOMe lifetime as a reporter of absolute V_m_.

### Monitoring EGF-induced hyperpolarization in A431 cells

A key advantage of red-shifted voltage sensors is the ability to record for long periods of time with low phototoxicity. To evaluate CRhOMe FLIM in this context, we recorded the response of serum-starved A431 cells to re-introduction of EGF (500 ng/mL) for 15 minutes. We previously performed this experiment with the green VoltageFluor VF2.1.Cl.^34^ Using VF2.1.Cl, we were able to make six, intermittent V_mem_ measurements within a 15 minute time window, with each measurement lasting 30 seconds. These previous experiments revealed a hyperpolarizing response to EGF stimulation, facilitated by the kinase activity of EGFR, dependent on internal Ca^2+^ release, and mediated by the action of the Ca^2+^-activated K^+^ channel, K_Ca3.1_.^21^

Using CRhOMe as a voltage indicator, we were able to record continuously for 15 minutes, with each measurement lasting 5 seconds across a field of view 1.5 fold larger than the VF2.1.Cl study (**Figure 4** and **S11**) for a total of ∼180 V_mem_ measurements, compared to 6 measurements with VF2.1.Cl, an order of magnitude increase in total imaging time. In EGF-treated samples, we observe an immediate lifetime decrease of 75 ps on average with a steady return to baseline that did not complete within 15 minutes (**Figure 4**), consistent with the expected hyperpolarization.^21^ Samples treated with imaging buffer vehicle show no change in τ_fl_ of CRhOMe (**Figure 4a**). The control TMCRhZero displayed a steady, slight shortening of the lifetime (**Figure 4d**), which was identical when treated with either vehicle or EGF. The spatial distribution of lifetime values in both vehicle and EGF-treated cells are intriguing and hint at the possibility of heterogenous V_mem_ distributions in otherwise “uniform” populations of cells that have been reported elsewhere.^43^

**Figure 4.**
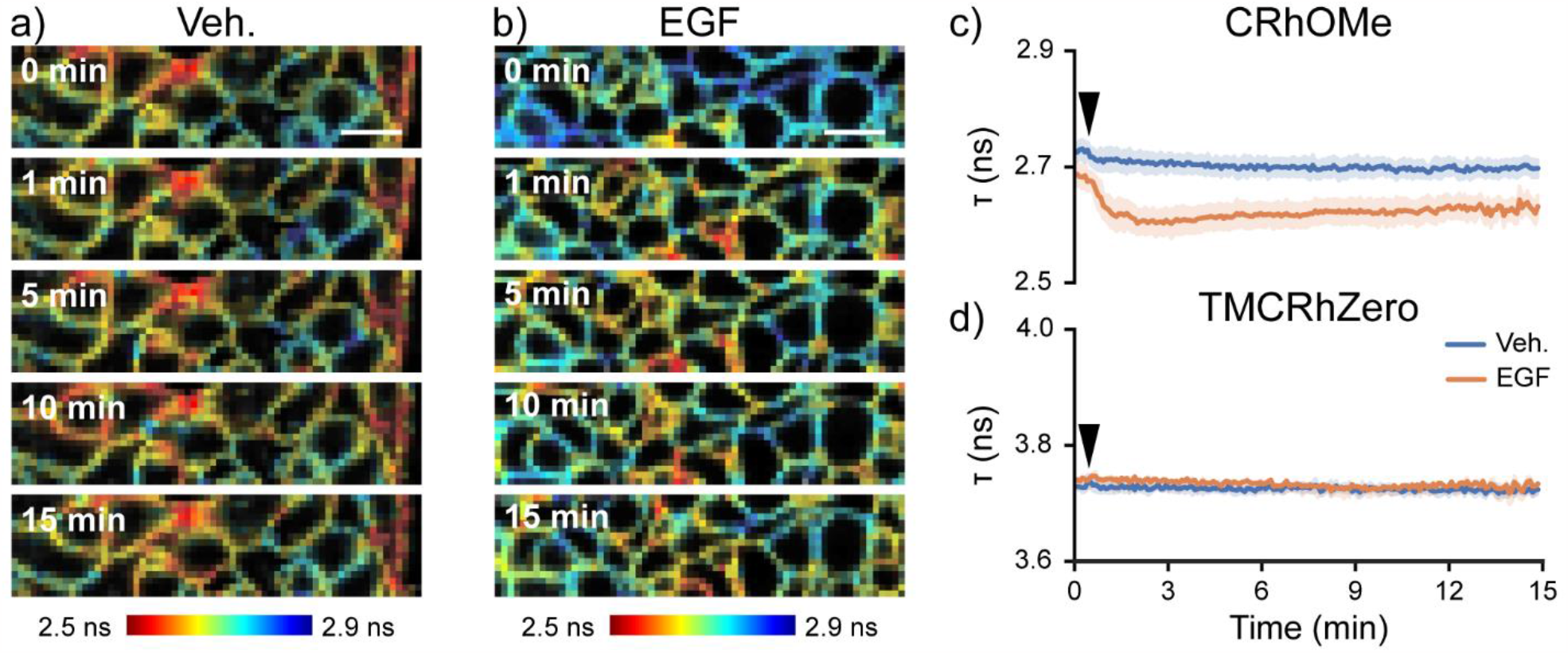
CRhOMe reports on EGF-induced hyperpolarization events in A431 cells with enhanced temporal and spatial resolution. Snapshots from a τ time series of serum-starved A431 cells treated with either **a)** vehicle (imaging buffer) or **b)** EGF. Scale bar is 20 μm. Each snapshot is acquired at the indicated time after starting the experiment. Vehicle or EGF is added 20 s into the experiment. **c)** Mean lifetime of CRhOMe (top) or **d)** TMCRhZero (bottom) across the full recording. Shading represents standard error of the mean. Vehicle or 500 ng/mL EGF was added at the black arrow. Sample sizes: CRhOMe Veh 5, EGF 5; TMCRhZero Veh 3, EGF 3.

### FLIM to monitor cardiac action potentials

To demonstrate the improved temporal resolution and stability to motion of our absolute V_mem_ sensing platform, we monitored the τ_fl_ of spontaneously beating human induced pluripotent stem cell derived cardiomyocytes (iCMs). Upon loading CRhOMe or TMCRhZero into iCM monolayers, we observed membrane localized fluorescence (**Figure 5, S12**). Because we are using a confocal pinhole larger than 1 airy unit to maximize photon count, some regions of the membrane appear as flat sheets where they traverse the optical section at a near-horizontal angle.

**Figure 5.**
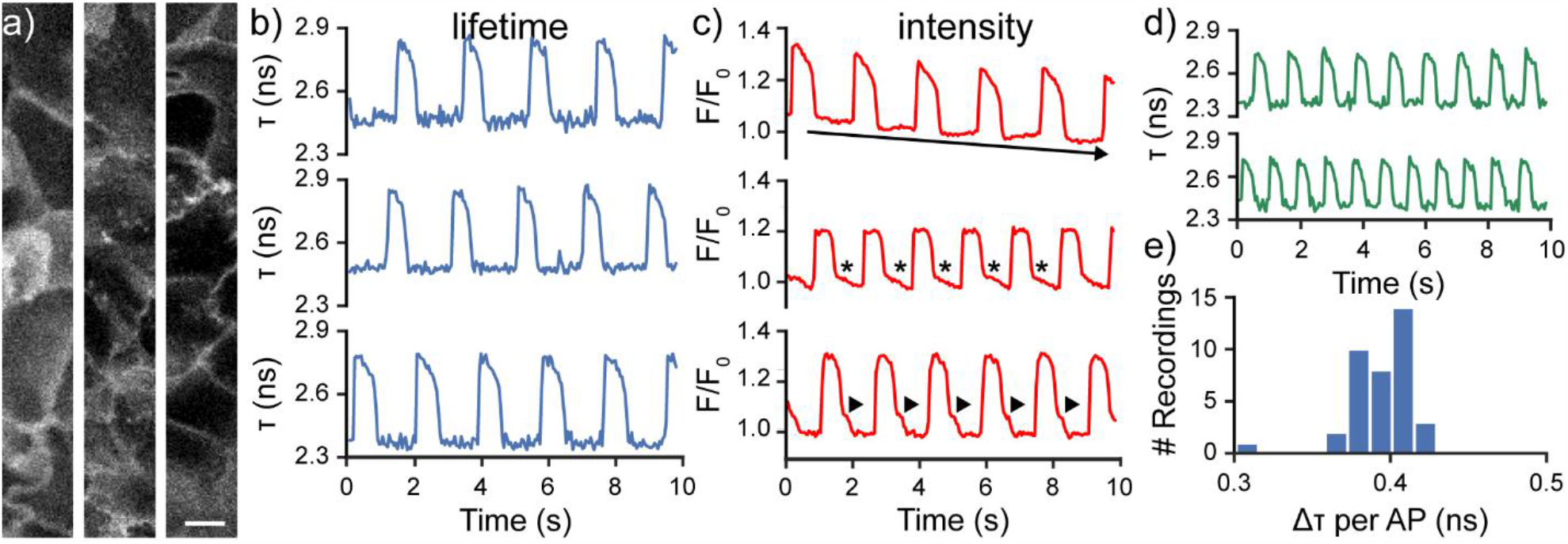
Fluorescence lifetime imaging of spontaneously beating cardiomyocytes using CRhOMe avoids artifacts associated with intensity-based imaging. **a)** Representative confocal images of regions of iCMs used for 20 Hz lifetime imaging. Images here are fluorescence intensity only and were acquired for 400 ms (line averaged) to improve contrast. iCMs were stained with 500 nM CRhOMe. Scale bar is 10 μm. **b)** Plots of fluorescence **lifetime** of CRhOMe vs time in active hiPSC-CMs, quantified from the fields of view in (a). **c)** Plots of fluorescence **intensity** of CRhOMe vs time in active hiPSC-CMs from the same regions as in a and b. Top panel: arrow depicts photobleaching; middle panel: asterisks depict apparent AP morphological artifact; bottom panel: arrowheads show AP artifact. None of these artifacts appears in the lifetime recording. **d)** Representative recordings of CRhOMe absolute V_mem_ imaging of iCMs treated with 1 μM isoproterenol, which accelerates beat rate. **e)** Amplitude of iCM action potentials, as measured by the change in τ of CRhOMe. Data represent 38 recordings from 8 iCM wells. Bin size for histogram was determined by the Freedman-Diaconis rule.

The action potential (AP) in iCMs is typically on the order of 500 ms in duration. Intensity-based V_mem_ imaging techniques in cardiomyocytes generally use sampling rates between 50 and 500 Hz,^44-46^ but fluorescence lifetime imaging studies in other systems rarely exceed frame rates of 0.1 Hz. Using TCSPC FLIM, we acquired a raw lifetime data stream at 40 Hz (25 ms acquisition per frame). We binned successive pairs of frames to produce a final recording with 20 Hz frame rate (**Figure 5d, Figure S13**). Because of CRhOMe’s high brightness and photostability, the limiting factor in the achievable frame rate was the speed at which the TCSPC electronics could process photons (the pile-up limit). We did not observe phototoxicity to the sample or drift in the lifetime; this is consistent with our ability to acquire continuously for 15 minutes in A431 cells (**Figure 4**).

We observed a stable lifetime baseline and consistent action potential morphology throughout all optical recordings (**Figure S13**). We used TMCRhZero to further verify that lifetime is insensitive to motion artifacts from the iCM contraction. Recordings with the voltage-insensitive TMCRhZero derivative are stable throughout the acquisition, despite normal contraction of the iPSCs (**Figure S14**).

In contrast to the stability of the lifetime recording, the photon count (fluorescence intensity) from cardio myocytes shows both a variable baseline and a variable AP waveform (**Figure 5c** and **S13b**), presumably from motion artifacts as the cardiomyocytes contract. Even though all wells and fields of view were loaded with the same concentration of dye (500 nM), the baseline intensity varied widely between recordings (**Figure S13b**). Differences in the amount of membrane in the field of view (**Figure 5a**), as well as differences in focal plane, are likely responsible for this. Normalizing the photon count into relative fluorescence units (**Figure 5c** and **S13c**), as is common with fluorescence intensity imaging, did not fully resolve these artifacts in either baseline drift or action potential morphology.

Using CRhOMe fluorescence lifetime as a proxy for absolute V_mem_, we catalogued the action potential amplitude, resting τ, and peak τ across 40 iCM recordings (**Figure 5e, Figure S13d, Figure S15**). We observe a consistent lifetime change of 0.394 ± 0.004 ns (mean ± SEM of 40 recordings across two differentiations). Resting τ (2.37 ± 0.01 ns) and peak τ (2.76 ± 0.01 ns) were somewhat more variable than the AP amplitude for these cells. Because of difficulties associated with electrophysiology in beating sheets of cardiomyocytes, we first attempted to apply the calibration from HEK293T cells to convert these lifetime changes to V_m_. Unfortunately, the sensitivity measured in HEK 293T cells (3.1 ps/mV) translates to a non-physical AP amplitude of around 127 mV in cardiomyocytes, based on an average lifetime change of 0.394 ns for each action potential. iCM action potentials are approximately 100 mV in amplitude, although they may range from 80-113 mV depending on differentiation conditions.^48-51^ The discrepancy between this value and the reported iCM AP amplitudes could be due to the previously observed slight dependence of VoltageFluor lifetime sensitivity on cell type.^34^ We attempted a gramicidin-based calibration in cardiomyocytes, but we found that the ionophore was toxic to cells before V_mem_ became fully depolarized (data not shown). A future electrophysiological calibration to relate CRhOMe lifetime to a known voltage in iCMs will enable a more quantitative analysis.

To push the temporal resolution of CRhOMe FLIM further, we treated iCMs with the β adrenoreceptor agonist isoproterenol, which increases beat rate (**Figure 5d**).^47^ Under these conditions, we observed spontaneous APs at approximately twice the frequency (∼1.2 Hz versus ∼0.6 Hz). Even with this more rapid activity, we were able to record the full AP waveform and a consistent baseline τ. As expected with an increase in beat rate, AP duration shortened in isoproterenol treated cells. AP amplitude appeared slightly smaller than in treated cells, and baseline τ remained unchanged (**Figure 5d**). The robustness of the CRhOMe FLIM signal during this treatment suggests that this could be a useful approach for interrogating cardiomyocyte voltage signaling under perturbation or throughout differentiation.

### Outlook/Conclusion

We developed a new VF dye, CRhOMe, which provides red to far-red excitation and emission, improved signal to noise ratios (SNR) compared to our previous best-in-class BeRST indicator, and a monoexponential fluorescence lifetime decay, which substantially simplifies its use in FLIM, requiring an order of magnitude fewer photons for an accurate fit.^35-36^ Used in fluorescence intensity mode, CRhOMe enables imaging of action potential dynamics in single trials in hippocampal neurons with improved SNR compared to BeRST. Used in lifetime mode, CRhOMe reports on physiological plasma membrane hyperpolarization induced by epidermal growth factor-mediated (EGF) signaling in mammalian cells with 15 minutes of continuous illumination, a >5-fold improvement over the first-generation VF-FLIM. CRhOMe is well tolerated in sensitive samples, and coupled with its brightness, fast response time, and mono-exponential fluorescence lifetime decay, this allows tracking of cardiac action potential dynamics in human induced pluripotent stem cell-derived cardiomyocytes (hiPSC-CMs), in both intensity and lifetime mode. Lifetime imaging of AP dynamics eliminates artifacts associated with moving cells and opens the door to future optical determination of resting membrane potential values in excitable cells. In our current implementation, the FLIM hardware, not the fluorophore, limits the maximum acquisition speed; combining CRhOMe with emerging methods for fast FLIM microscopy^52^ would enable even greater acquisition speeds. Overall, CRhOMe is a promising new VF dye for the resolution and quantification of membrane potentials in both excitable and non-excitable cells.

## Acknowledgments

We acknowledge support from the NIH (R01NS098088) and the DOE (DE-SC0023184). A.M.M.G., J.R.L.-D., and S.C.B. were supported, in part, by a training grant from the NIH (T32GM066698). S.K.Y.-W. was supported, in part, by a training grant from the NIH (T32GM008295). G.O. was supported by a Gilliam Fellowship from HHMI. FLIM experiments were performed at the CRL Molecular Imaging Center, RRID:SCR_017852, at UC Berkeley, supported by the NIH (S10OD025063). We thank the C. Chang lab for use of their FLIM microscope. We thank Holly Aaron and Feather Ives for expert technical assistance, advice, and support.

## Figures and Schemes

**Scheme 1.**
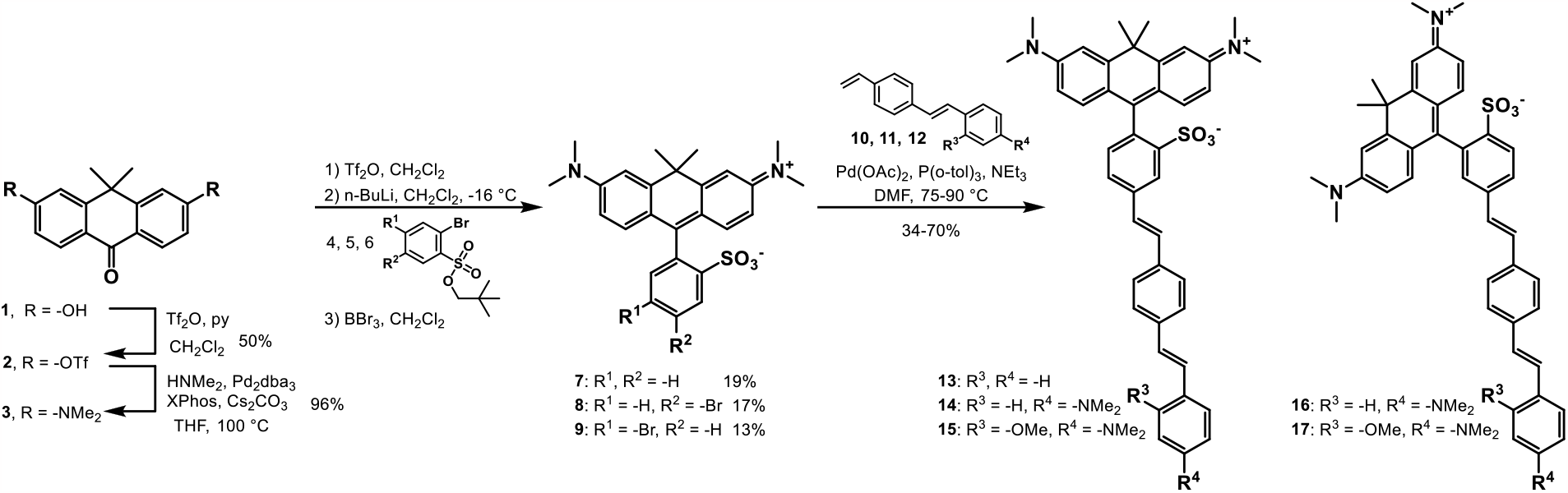
Synthesis of carborhodamine fluorophores (7-9) and voltage dyes (13-17).

