## Supporting Information Document for "A red-emitting carborhodamine for monitoring and measuring membrane potential"

DOI Placeholder

### Chemical Synthesis and Characterization

Chemical reagents and solvents (anhydrous) were purchased from commercial suppliers and used without further purification. Compounds **1**,[1] 2,4-dibromobenzenesulfonyl chloride,[2] sulfonates **4-5**,[3] styrene **10**,[4] styrene **11**,[5] and styrene **12**[4] were prepared according to the literature procedures. All reactions were carried out in flame-dried flasks sealed with septa and conducted under a nitrogen atmosphere. Thin layer chromatography (TLC) (silica gel, F254, 250 μm) was performed on precoated TLC glass plates and were visualized by fluorescence quenching under UV light. Flash column chromatography was performed on Silicycle Silica Flash F60 (230–400 Mesh) using a forced flow of air at 0.5–1.0 bar. NMR spectra were recorded on a Bruker AVB-400 MHz, or at the QB3 Central California 900 MHz NMR Facility. Chemical shifts (δ) are expressed in parts per million (ppm) and are referenced to $CDCl_3$ (7.26 ppm, 77.0 ppm) or $CD_3OD$ (3.31 ppm, 49.0 ppm). Coupling constants are reported as Hertz (Hz). Splitting patterns are indicated as follows: s, singlet; d, doublet; t, triplet; q, quartet; dd, doublet of doublet; m, multiplet.

High-resolution mass spectra (ESI, EI) were measured by the QB3/Chemistry mass spectrometry service at University of California, Berkeley. High performance liquid chromatography (HPLC) and low resolution ESI Mass Spectrometry were performed on an Agilent Infinity 1200 analytical instrument coupled to an Advion CMS-L ESI mass spectrometer. The column used for the analytical HPLC was Phenomenex Luna 5 μm C18(2) (4.6 mm I.D. × 150 mm) with a flow rate of 1.0 mL/min. The mobile phases were MQ-$H_2O$ with 0.05% trifluoroacetic acid (eluent A) and HPLC grade MeCN with 0.05% trifluoroacetic acid (eluent B). Absorbance signals were monitored at 254, 380, 450, 615, and 650 nm.

*Materials*

Dyes were stored either as solids at room temperature or as 1 mM stock solutions in DMSO at -20° C. All stock concentrations were determined via the absorbance of the carborhodamine chromophore using a 2501 Spectrophotometer (Shimadzu).

Epidermal growth factor (Peprotech) was made up as a 1 mg/mL stock solution in water and stored at -80°C. Gramicidin (Sigma-Aldrich) was purchased as a mixture of A, B, C, and D from *Bacillus brevis*. 1 μg/mL stocks of gramicidin were made up in DMSO and stored at -20°C. Isoproterenol was a gift from the Healy lab at UC Berkeley and was stored as a 10 mM stock in DMSO at -20°C.

### Spectroscopic Studies

UV-Vis absorbance and fluorescence spectra were recorded using a 2501 Spectrophotometer (Shimadzu) and a Quantamaster Master 4 L-format scanning spectrofluorometer (Photon Technologies International). The fluorometer is equipped with an LPS-220B 75-W xenon lamp and power supply, A-1010B lamp housing with integrated igniter, switchable 814 photon-counting/analog photomultiplier detection unit, and MD5020 motor driver. Samples were measured in 1-cm path length quartz cuvettes (Starna Cells).

The maximum absorption wavelength ($\lambda_{max}$), maximum emission wavelength ($\lambda_{em}$), extinction coefficient (ε), and quantum yields of fluorescence were taken in MeOH or PBS (0.1 M $Na_2HPO_4$, 0.1 M NaCl, pH 7.4) using stock solutions of TMCRh dyes in DMSO (0.5-1 mM); the reported values for ε is an average (n = 2). Relative quantum yields ($\Phi_{fl}$) were calculated by comparison to standard cresyl violet perchlorate ($\Phi = 0.54$ in MeOH).[6] Briefly, stock solutions of standards were prepared in DMSO (1 mM) and diluted with appropriate solvent (1:1000 dilution). Absorbance and emission ($\lambda_{ex} = 560$ nm) were taken at 5 concentrations. The absorbance value at the $\lambda_{ex}$ was plotted against the integration of the area of fluorescence curve (570-800 nm). The $\lambda_{ex}$ ensured the full fluorescence area of the dyes excited at 560 nm was used for $\Phi_{fl}$ calculations. The slope of the linear best fit of the data was used to calculate the relative $\Phi_{fl}$ by the equation $\Phi_{fl(X)} = \Phi_{fl(R)}(S_R/S_X)(\eta_X/\eta_R)^2$, where $S_R$ and $S_X$ are the slopes of the standard and unknown, respectively, and η is the refractive index of the solution.

### Cell Culture

*Immortalized cell line culture*

HEK293T and A431 cells were obtained from the UC Berkeley Cell Culture Facility. Both cell lines were verified by STR profiling (short tandem repeat) and were tested routinely for mycoplasma. Cell lines were discarded after 25 passages. Cells were maintained in complete DMEM (Gibco, Thermo Fisher Scientific) supplemented with 4.5 g/L glucose, 2 mM GlutaMAX (Gibco), and 10% FBS (Seradigm) in a 37°C humidified incubator at 5% $CO_2$. Cells were passaged into fresh complete media every few days following dissociation with trypsin-EDTA (Gibco, 0.05% for HEK293T, 0.25% for A431). Residual trypsin was removed from A431 cells by centrifugation for 5 minutes at 300xg.

For imaging experiments, cells were plated onto prepared poly-D-lysine (PDL)-coated coverslips. Coverslips (#1.5, either 12 mm or 25 mm diameter, Electron Microscopy Sciences) were acid washed for 2-5 hours in 1 M HCl. Coverslips were then washed three times overnight in 100% ethanol, followed by three times overnight in MilliQ (Millipore) purified water. Coverslips were sterilized by heating for 2-3 hours in a glassware oven to 150°C. Prior to seeding of cells, coverslips were incubated in 1x PDL (Sigma-Aldrich, made as a 0.1 mg/mL solution in phosphate-buffered saline with 10 mM $Na_3BO_4$) for 1-10 hours and then washed twice with sterile water and twice with 1x Dulbecco's Phosphate Buffered Saline (dPBS, Gibco).

For probe loading and gramicidin treatment experiments, HEK293T were seeded onto prepared coverslips in complete DMEM at a density of 42-52 $\times10^3$ cells per $cm^2$ (in a 6 well or 24 well tissue culture plate, Corning) and imaged approximately 24 hours after plating. For electrophysiology experiments, HEK293T were seeded at 26,000 cells/$cm^2$ in low glucose DMEM (Gibco; 1 g/L glucose, 1 mM sodium pyruvate, 2 mM GlutaMAX, 10% FBS) and used 12-24 hours after plating.

A431 cells were serum deprived prior to use.[7] Two days before imaging experiments, cells were trypsinized and suspended in complete media with 10% FBS. The cells were then spun down for 5 minutes at 500xg and resuspended in low serum DMEM (4.5 g/L glucose, 2 mM GlutaMAX, 2% FBS). Cells were then seeded onto PDL-coated glass coverslips at a density of 83,000 cells/$cm^2$ in the low serum DMEM. 3.5-5.5 hours prior to imaging, media was exchanged for serum-free DMEM (4.5 g/L glucose, 2 mM GlutaMAX). Cells were loaded with dye after 4-5.5 hours in the serum-free media.

*Culture of human induced pluripotent stem cell (hiPSC) derived cardiomyocytes*

hiPSCs (WTC11) were cultured on Matrigel (1:100 dilution; Corning)-coated 12 well-plates in StemFlex medium (Gibco). When the cell confluency reached 80–90%, which is referred to as day 0, the medium was switched to RPMI 1640 medium (Life Technologies) containing B27 minus insulin supplement (Life Technologies) and 10 μM CHIR99021 GSK3 inhibitor (Peprotech). At day 1, the medium was changed to RPMI 1640 medium containing B27 minus insulin supplement only. At day 3, medium was replaced with RPMI 1640 medium containing B27 supplement without insulin, and 5 μM IWP4 (Peprotech). On day 5, medium was replaced with RPMI 1640 medium containing B27 minus insulin supplement. On day 7, medium was replaced with RPMI 1640 containing B27 with insulin supplement. After day 7, the medium was changed every two days. Confluent contracting sheets of beating cells appear between days 7 to 15.

Beating sheets were treated with collagenase II for 60-75 minutes. The collagenase solution was carefully transferred to cold DMEM, making sure cardiac sheets were not disturbed. Trypsin (0.25%) was added to dissociated sheets for 4-8 minutes and plated onto 6 well-plates coated with Matrigel (1:100 dilution) in RPMI 1640 medium containing B27 supplement plus ROCK inhibitor Y-27632. 24 hours later, the medium was replaced with fresh RPMI/B27 without ROCK inhibitor. Cardiomyocytes were maintained for 7 days, replacing media every other day, and then were switched to RPMI 1640 medium (-glucose) supplemented with 4 mM sodium lactate (Sigma Aldrich). Cells were maintained in this media for 7 days, replacing every other day, then switched back to RPMI/B27 containing glucose.

To prepare these purified cardiomyocytes for imaging, lactate purified sheets were dissociated with 0.25% trypsin-EDTA (4-8 minutes, depending on density and quality of tissue) and plated onto Matrigel (1:100)-coated Ibidi ® 24 well µ-plates (cat no. 82406**)** in RPMI 1640 medium containing B27 supplement (containing insulin). Medium was changed every 3 days until imaging. For loading hiPSC cardiomyocytes, voltage dyes were diluted 1 in 1000 in RPMI 1640 with B27 supplement minus Phenol Red to the desired final concentration. Cardiomyocytes were incubated in this solution for 20 minutes at 37 °C, then exchanged with dye-free RPMI 1640 with B27 supplement minus Phenol Red. Imaging experiments were performed approximately two weeks after cells were seeded onto Matrigel-coated 24 well plates.

*Neuronal cell culture*

All animal procedures were approved by the UC Berkeley Animal Care and Use Committees and conformed to the NIH Guide for the Care and Use and Laboratory Animals and the Public Health Policy.

Hippocampi were dissected from embryonic day 19 Sprague Dawley rats (Charles River Laboratory) in cold, sterile HBSS (zero $Ca^{2+}$, zero $Mg^{2+}$, phenol red). All dissection products were supplied by Invitrogen, unless otherwise stated. Hippocampal tissue was treated with trypsin (2.5%) for 15 min at 37 °C. The tissue was triturated using fire polished Pasteur pipettes, in minimum essential media (MEM) supplemented with 5% FBS, 2% B-27, 2% 1M dextrose (Fisher Scientific) and 1% GlutaMax. The dissociated cells were plated onto 12 mm diameter coverslips (Fisher Scientific) pre-treated with PDL (as above) at a density of 25-30,000 cells per coverslip in MEM supplemented media (as above). Neurons were maintained at 37 °C in a humidified incubator with 5% $CO_2$. At 1 day in vitro (DIV), half of the MEM supplemented media was removed and replaced with Neurobasal media containing 2% B-27 supplement and 1% GlutaMax. Imaging was performed on mature neurons 13-16 DIV.

Unless stated otherwise, for loading of hippocampal neurons, DMSO stock solutions of TMCRh dyes (1 mM) were diluted directly into HBSS to working concentrations. For neurons, the typical working concentration was 500 nM. Neurons were incubated for 15 min with TMCRh dyes at 37 °C before exchanging dye/HBSS for HBSS without any dye. All epifluorescence imaging was performed in HBSS (with $Ca^{2+}$ and $Mg^{2+}$) at room temperature.

**Fluorescence microscopy**

Imaging was performed on an AxioExaminer Z-1 (Zeiss) equipped with a Spectra-X Light engine LED light (Lumencor), controlled with Slidebook (v6, Intelligent Imaging Innovations). Images were acquired with a W-Plan-Apo 20x/1.0 water objective (20x; Zeiss). Images were focused onto either an OrcaFlash4.0 sCMOS camera (sCMOS; Hamamatsu) or an eVolve 128 EMCCD camera (EMCCD; Photometrix). For TMCRh images, the excitation light was delivered from a LED (4.5 W/cm$^2$; 20 ms exposure time) at 631/28 (bandpass) nm and emission was collected with a quad-band emission filter (430/32, 508/14, 586/30, 708/98 nm, Semrock) after passing through a quad-band dichroic mirror (432/38, 509/22, 586/40, 654 nm LP, Semrock).

For epifluorescence imaging experiments, carborhodamine voltage dyes were loaded at 500 nM in HEK293T HBSS (Gibco) for 20 minutes at 37°C in a humidified incubator with 5% $CO_2$. Coverslips were washed once with HBSS and transferred to fresh HBSS for imaging. All epifluorescence imaging was conducted under ambient atmosphere.

*Whole cell patch clamp electrophysiology*

Pipettes were pulled from borosilicate glass with filament (Sutter Instruments, Novato, CA) with resistances ranging from 4 to 7 MΩ with a P97 pipette puller (Sutter Instruments). Internal solution composition, in mM (pH 7.25, 285 mOsmol/L): 125 potassium gluconate, 10 KCl, 5 NaCl, 1 EGTA, 10 HEPES, 2 ATP sodium salt, 0.3 GTP sodium salt. EGTA (tetraacid form) was prepared as a stock solution in 1 M KOH before addition to the internal solution. Pipettes were positioned with an MP-225 micromanipulator (Sutter Instruments). Voltage clamp protocols for epifluorescence experiments were not corrected for liquid junction potential. For voltage clamp protocols during fluorescence lifetime experiments, a liquid junction potential of -14 mV was determined by the Liquid Junction Potential Calculator in the pClamp software package (Molecular Devices, San Jose, CA), and voltage step protocols used in conjunction with fluorescence lifetime measurements were corrected for this offset.

Electrophysiology recordings were made with an Axopatch 200B amplifier and digitized with a Digidata 1440A (Molecular Devices). For epifluorescence voltage sensitivity measurements, cells were held at -60 mV and potentials from +100 to -100 mV were applied in descending order with 20 mV steps. Signals were sampled at 50 kHz and filtered with a 5 kHz low-pass Bessel filter. Correction for pipette capacitance was performed in the cell attached configuration.

For voltage sensitivity measurements on the fluorescence lifetime system, cells were held at -60 mV and potentials from +80 to -80 mV were applied. Voltage steps were applied with a variable order of -80, -40, 0, and +40 mV steps, followed by a +80 mV step as the final voltage applied, with each voltage step holding for 15 seconds. Signals were sampled at 50 kHz and filtered with a 5 kHz low-pass Bessel filter. Correction for pipette capacitance was performed in the cell attached configuration. Recordings were only included if they maintained a 30:1 ratio of membrane resistance ($R_m$) to access resistance ($R_a$) and an $R_a$ value below 30 MΩ throughout the recording.

*Neuronal imaging*

Extracellular field stimulation was delivered by a SD9 Grass Stimulator connected to a recording chamber containing two platinum electrodes (Warner), with triggering provided through the same Digidata 1440A digitizer and pCLAMP 9 software (Molecular Devices) that ran the electrophysiology. Action potentials were triggered by 1 ms 60 V field potentials delivered at 5 Hz. To prevent recurrent activity, the HBSS bath solution was supplemented with synaptic blockers: 10 μM 2,3-Dioxo-6-nitro-1,2,3,4- tetrahydrobenzo[f]quinoxaline-7-sulfonamide (NBQX; Santa Cruz Biotechnology) and 25 μM DL-2-Amino-5-phosphonopentanoic acid (APV; Sigma-Aldrich). For both evoked action potentials and spontaneous activity, images were binned 4x4 to allow sampling rates of 0.5 kHz and 2500 frames (5 s) were acquired for each recording. Images were acquired with a LED (1.85 W/cm$^2$; 2 ms exposure time) at 631/28 (bandpass) nm and emission was collected with a quad-band emission filter (430/32, 508/14, 586/30, 708/98 nm) after passing through a quad-band dichroic mirror (432/38, 509/22, 586/40, 654 nm LP).

*Sample preparation and treatment for fluorescence lifetime imaging*

For all experiments following concentration optimization (**Fig S7**) in HEK293T or A431, CRhOMe and TMCRhZero were used at 300 nM, directly from a 1 mM (3333x) DMSO stock (final DMSO concentration in loading solution, 0.03). 1x loading solutions were prepared in imaging buffer (IB, made in-house; composition in mM: 139.5 NaCl, 10 HEPES, 5.6 D-glucose, 5.33 KCl, 1.26 $CaCl_2$, 0.49 $MgCl_2$, 0.44 $KH_2PO_4$, 0.41 $MgSO_4$, 0.34 $NaH_2PO_4$; 290 mOsm/L; pH 7.25). Osmolarity was determined with a µOsmette (Precision Instruments). Cells were incubated at 37°C for 20-25 minutes in loading solution, washed once in IB, and transferred to fresh IB in an Attofluor cell imaging chamber (Invitrogen).

iCMs were loaded in RPMI with B27 supplement without phenol red containing 500 nM of CRhOMe or TMCRh Zero (0.05% final concentration of DMSO). Media was exchanged for fresh RPMI-B27 without phenol red before imaging.

Gramicidin solutions were prepared at either 0.5 µg/mL or 1 µg/mL in IB immediately before use. Immediately after dye loading, HEK293T and A431 were incubated for 5 minutes at room temperature in IB containing either gramicidin or DMSO vehicle. Coverslips were then transferred to fresh IB with gramicidin or vehicle and imaged.

For EGF treatment experiments in A431 cells, 2x EGF (1 µg/mL) solutions were prepared in IB on the day of the experiment. Serum-starved A431 cells were loaded with 300 nM dye in IB after they had spent 4-5.5 hours in the serum-free media. Approximately twenty seconds into the FLIM recording, 500 µL of IB or IB containing 2x EGF was gently pipetted into the imaging chamber (which contained 500 µL IB). IB control recordings were made prior to EGF treatment recordings from a different field of view on the same sample.

Most imaging experiments except for those involving serum starved A431 cells were conducted under ambient atmosphere and at room temperature (18-22°C). To improve cell viability, some of the cardiomyocyte experiments were conducted with the XL incubation chamber on the LSM 880 set to 25°C. To minimize temperature fluctuations during extended recordings with serum-starved A431 cells, all work with these cells was conducted with the system incubation chamber set to 25°C and solutions pre-warmed to approximately 25°C. Samples were generally kept on the microscope for under 20 minutes; longer term EGF studies required samples to be kept on the microscope for up to 45 minutes.

*Time Correlated Single Photon Counting (TCSPC) FLIM Studies*

Instrument 1

Concentration curve data, A431 data, and cardiomyocyte data were acquired on an inverted LSM 880 confocal microscope (Carl Zeiss AG, Oberkochen, Germany) equipped with a FLIM upgrade kit (PicoQuant GmbH, Berlin, Germany). The LSM 880 was controlled with Zen Black software (Zeiss); the TCSPC unit was controlled with SymPhoTime 64 software (PicoQuant). Pulsed excitation at 40 MHz was supplied with a 640 nm diode laser (PicoQuant LDH-P-C-640B with a ZET635/20x cleanup filter, AHF/Chroma; average output wavelength 637 nm) controlled by a PDL-800-D laser controller (PicoQuant). Laser intensity from the PDL-800-D was kept at 24% to optimize instrument response function (IRF) shape; excitation power at the sample was controlled with a micrometer in the laser combining unit (PicoQuant). Power at the sample was adjusted to maximize count rate without exceeding the pile-up limit (a count rate of 4e6 photons/s at any individual pixel, 10% of the laser repetition rate). Because the dye was localized to membranes, large regions of the field of view were relatively dark, so average count rates varied from 1e5-2e6 counts per second depending on the sample and field of view. Average power at the sample ranged from 1-5 µW.

Excitation light was coupled to the microscope with a polarization-maintaining single mode fiber and passed through a line pass dichroic (MBS 405/488/560/640) before reaching the sample. Emitted photons were collected with a 63x oil/1.4 NA Apochromat objective (Zeiss) immersed in Immersol 518F (Zeiss). The confocal pinhole was set to 300 (4.5 AU, optical slice thickness of approximately 3.9 µm), increasing the size of the optical section but improving photon count rates for TCSPC. A 690/70 nm bandpass emission filter (AHF) was used. Single photons were detected with a PMA-Hybrid 40 detector (PicoQuant) and processed with a TH260 Pico Dual TCSPC Unit (T3 TCSPC mode). The detector parameters used were the following: constant fraction discriminator (CFD) -100 mV, zero cross -10 mV, offset -47000 ps. Sync parameters used were the following: CFD -150 mV, zero cross -10 mV, offset 0 ps, sync divider 8. Data were acquired with 1000 TCSPC time channels (0.025 ns/channel).

Instrument 2

Electrophysiological lifetime calibration data were collected via TCSPC on an inverted LSM 980 confocal microscope (Carl Zeiss AG, Oberkochen, Germany) equipped with a SPC-150NX single photon counting card (Becker and Hickl). The LSM 980 was controlled with Zen Black software (Zeiss), and the TCSPC unit was controlled with SPCM software (Becker-Hickl). Pulsed excitation at 50 MHz was supplied with a 562 or 640 nm diode laser (BDS-SM-640-FBC-101, Becker-Hickl) and controlled through a LHB-104 Laser Hub (Becker-Hickl). Power was adjusted with a neutral density filter wheel to maximize count rate without exceeding pile-up limit ($5 \times 10^6$ photons/s at an individual pixel, 10% of the laser repetition rate). Average power at the sample ranged from 1-3 µW.

Excitation light was coupled to the microscope with a polarization-maintaining fiber and passed through a 405/485/560/640 beam splitter (Zeiss). Emitted photons were collected with a 40x oil/1.3 NA Apochromat objective (Zeiss), immersed in Immersol 518F (Zeiss). The confocal pinhole was set to 100 µm (1.68 AU, 0.9 µm optical section). The emitted photons were passed through a 676/37 emission filter (Semrock) and were detected with an HPM-100-40 hybrid detector (Becker-Hickl). The detector parameters used were the following: constant fraction discriminator (CFD) -50.98 mV, zero cross 5.29 mV. Sync parameters used were the following: CFD -50.98 mV, zero cross 5.29 mV, offset 5.10 %, sync divider 1, dither range 1/16. Data were acquired with 256 TCSPC time channels.

General Acquisition Procedures

The IRF was measured each day from a sample of 1 mg/mL erythrosin B through the same imaging substrate (#1.5 coverglass or ibidi 24 well plate). 1 µM TMCRh (small molecule dye head) in water was recorded daily as an additional standard to verify correct instrument function.

Monodirectional scanning was used for all applications except iCM spontaneous activity, where bidirectional scanning was used. Bidirectional scan correction is not possible in SymPhoTime, so the images of cardiomyocyte field of view (FOV) were acquired in Zen (same acquisition settings as FLIM, with the addition of 16x line averaging). Spatial and temporal resolution of the data acquisition for specific applications is enumerated in Table S1.

*Fluorescence lifetime data analysis*

Time resolved fluorescence decays I(t) were modeled as a monoexponential decay (equation 1) in custom Matlab code via a weighted least-squares approach. A represents the amplitude of the photon count signal and τ represents the fluorescence lifetime. TMCRhZero was well-modeled by a single exponential decay; CRhOMe contained slight structure in the residuals but two component decays resulted in overfitting. BeRST1 was not well-modeled by a single exponential decay, and for that reason not pursued for further characterization; lifetimes reported for BeRST1 are the amplitude-weighted average of a two-component decay fit, resulting from the same custom Matlab code.

$$I(t) = Ae^{-t/\tau} \qquad \text{Equation 1}$$

Minimization of the reduced chi squared ($\chi^2$, equation 2) was used to determine the final parameter values, where n is the number of free parameters (number of fit time channels – number of free parameters in model), st is the first fit time channel, fi is the final fit time channel, z is the value of the fit model, and x is the value of the experimentally measured decay. Poisson weighting of the time channels was used.

$$\chi^2 = \frac{1}{n}\sum_{i=st}^{fi}\frac{(z_i - x_i)^2}{|z|} \qquad \text{Equation 2}$$

The built-in minimization function fmincon with the interior-point algorithm was used, with the following settings: StepTolerance 1e-4, OptimalityTolerance 1e-4, ConstraintTolerance 1e-6. All fits converged in fewer than the maximum number of function evaluations (1e5). The fluorescence lifetime τ was constrained between 0 and 10 ns.

The experimentally measured instrument response function (IRF, erythrosin B fluorescence decay) was cropped to time channels 40-60 out of 1000 channels (25 ns) for data from the LSM 880 microscope and cropped to time channels 29-48 out of 256 channels (20 ns) for data from the LSM 980 microscope, and reconvolved with the decay model before the reduced chi squared was determined. The time resolved signal was binned by a factor of 4 in time (to 250 channels representing 25 ns) for the data from the LSM 880 before fitting was performed to reduce fit noise with low photon counts, while the data from the LSM 980 was not binned in time. Time channels 20 through 980 (st through fi) of the unbinned signal from the LSM 880 data were used in the fit where photon counts were sufficient. Otherwise, the final time bin used was the first instance of zero photon counts in a time channel. The shift between the IRF and the measured time resolved decays was fixed to 0 for the data from the LSM 880; for the data from the LSM 980 microscope, the shift was fixed to -1, as this parameter can vary between TCSPC systems. The offset (time-independent background) was also fixed to 0, as the dark counts on both systems were generally negligible (20-100 counts per second during experiments). A threshold of 5000 total photons was used in pixelwise analysis. Pixels with photon counts below this threshold were excluded from analysis and appear black in images. Spatial binning was performed before fitting for pixel-by-pixel analysis (**Table S1**). For global analysis of cardiomyocyte data, all photons in the field of view were used to fit one decay per time point.

### Image Analysis

*Image analysis (epifluorescence microscopy)*

For image intensity measurements, regions of interest were drawn around cells or neuronal cell bodies and the mean fluorescence was calculated in ImageJ (FIJI, NIH). Background fluorescence was subtracted by measuring the fluorescence where no cells grew. Analysis of voltage sensitivity in HEK cells was performed using ImageJ (FIJI). Briefly, a region of interest (ROI) was selected automatically based on fluorescence intensity and applied as a mask to all image frames. Fluorescence intensity values were calculated at known baseline and voltage step epochs. For analysis of voltage responses in neurons, regions of interest encompassing cell bodies (all of approximately the same size) were drawn in ImageJ and the mean fluorescence intensity for each frame extracted. ΔF/F values were calculated by first subtracting a mean background value from all raw fluorescence frames, to give a background subtracted trace (bkgsub). A baseline fluorescence value (Fbase) is calculated from the median of all the frames, and subtracted from each timepoint of the bkgsub trace to yield a ΔF trace. The ΔF was then divided by Fbase to give ΔF/F traces. No temporal averaging has been applied to any voltage traces.

*Analysis of cardiomyocyte spontaneous action potential recordings*

For action potential detection, lifetime recordings were transformed into a series of differences between successive frames ("difference trace"). This difference trace was filtered with a 6$^{th}$ order low pass Butterworth filter with the signal processing toolkit in SciPy (scipy.signal). A threshold that differentiated between action potentials and baseline was selected manually. The peak lifetime was defined as the value of the lifetime at the index identified by the above process. For each recording, the peak is reported as the mean of peaks of all action potentials in the recording. No filtering was used on lifetime traces shown or on lifetime data used for analysis; filtering was used exclusively in peak identification.

The baseline was defined from a series of 15 frames (750 ms) before the peak of each action potential (AP), beginning 20 frames before the peak and ending 5 frames before the peak. While this omits some areas of the baseline, it allows automatic processing at a variety of beat rates. The baseline reported for each recording is the average lifetime in all of these 15 frame windows except the first, which was often cut off by the start of the recording. For example, if a recording contained 6 APs, the baseline τ would represent the average of 75 frames of lifetime data (15 frames per AP from 5 APs).

The peak height was calculated as the average baseline subtracted from the average peak for each trace. For recordings containing gramicidin or made with TMCRhZero, no APs were visible, so the lifetime reported is the average across all frames in the 10 second recording.

For analysis of the photon count data (**Figure S13**), photon count data were taken from the same indices as defined by the lifetime trace (see above). The relative fluorescence intensity ($F/F_0$) was calculated by dividing each value in the photon count trace by the average photon count baseline.

*FLIM electrophysiology calibration curves*

Lifetime values were determined for membrane potentials using the lifetime-voltage standard curves determined with whole-cell voltage-clamp electrophysiology (**Fig 3**). The lifetime reported is the mean lifetime across all pixels in an ROI. ROIs were defined as the entire plasma membrane for the individual measured cell.

The relationship described by these curves between lifetime and membrane potential was determined by linear regression, resulting in a sensitivity (slope (m), ps/mV) and a 0 mV lifetime (y-intercept (b), ps) for each cell. The average sensitivity and 0 mV lifetime were determined from each of the cells. These means of these values were calculated to determine an "average calibration," which can be used to translate between measured lifetime and membrane voltage (equation 3). For quantifying and discussing changes in voltage, only the sensitivity (slope) is necessary (equation 4).

$$\tau = m * V_{mem} + b \qquad \text{Equation 3}$$

$$\Delta V_{mem} = \frac{\Delta\tau}{m} \qquad \text{Equation 4}$$

Differences in the measured lifetime at a defined reference point (for us, the 0 mV lifetime) across measured cells provides an estimate of the voltage-independent noise for the lifetime-voltage calibration of CRhOMe, and thus the resolution we can expect when using this calibration. We report this resolution as the root-mean-square deviation (RMSD) between the optically determined voltage using the developed calibration ($V_{FLIM}$) and the voltage of the cell as set by voltage clamp ($V_{ephys}$). The RMSD of a set of n measurements (equation 5) can be determined from the variance (equation 6) and the bias (equation 7) of the estimator ($V_{FLIM}$) as compared to the assumed "true" value, in this case determined by electrophysiology ($V_{ephys}$).

$$RMSD = \sqrt{\sigma^2 + Bias^2} \qquad \text{Equation 5}$$

$$\sigma^2 = \frac{1}{n}\sum_{i=1}^{n}(V_{FLIM,i} - V_{ephys,i})^2 \qquad \text{Equation 6}$$

$$Bias = \frac{1}{n}\sum_{i=1}^{n} V_{FLIM,i} - \frac{1}{n}\sum_{i=1}^{n} V_{ephys,i} \qquad \text{Equation 7}$$

The variations in lifetime are larger between cells than within a cell, and thus the voltage resolution of CRhOMe lifetime in HEK 293T cells can be estimated with two values: the intra-cell resolution and the inter-cell resolution. The "intra-cell" resolution is a measure of the voltage resolution expected for tracking and quantifying changes within an individual cell, determined by the RMSD between an individual cell's lifetime-based estimate of $V_{mem}$ and the electrophysiologically determined $V_{mem}$ value. For CRhOMe, this intra-cell resolution is 6.36 ± 0.76 mV (RMSD, ± SEM). The "inter-cell" resolution is the single-trial resolution of estimating a cell's membrane potential based on the average calibration from all cells (i.e. with no prior information about that particular cell). For CRhOMe in HEK 293T cells, this inter-cell resolution is 21 mV.

*Statistical analysis*

Data are reported as either mean ± standard deviation (SD) or mean ± standard error of the mean (SEM) throughout the text (as indicated). All aggregated data were acquired across at least two independent experimental days.

For analysis of differences between groups, all datasets were tested for normality (Shapiro-Wilk test) and homoscedasticity (Levene's test on the median) before ANOVA was applied. For data that were both normal and homoscedastic, Fisher's one-way ANOVA with Tukey-Kramer post hoc tests was used ($p>0.05$ on both the Shapiro-Wilk test and Levene's test). For data that were non-normally distributed and homoscedastic, the Kruskal-Wallis rank sum test with Dunn's test for pairwise post hoc tests was used. For data that were heteroscedastic ($p<0.05$ on Levene's test), Welch's one-way ANOVA with Games-Howell post hoc tests was used, even if the data were significantly non-normal by Shapiro-Wilk. All statistical analysis was performed in Python using the SciPy, Scikit-Posthocs and Pingouin[8] statistical packages.

**Detailed Synthetic Procedures**

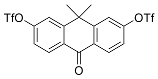

*9,9-dimethyl-10-oxo-9,10-dihydroanthracene-2,7-diyl bis(trifluoromethanesulfonate) (2):*

To a solution of 1[1] (2.45 g, 9.643 mmol) in $CH_2Cl_2$ (96 mL) cooled to 0 °C was added pyridine (6.10 g, 77.14 mmol, 8 eq), then trifluoromethanesulfonic anhydride (10.8 g, 38.57 mmol, 4 eq). The reaction was warmed to room temperature and stirred for 2h. It was then diluted with water and extracted with $CH_2Cl_2$ (3x). The organic extracts were combined and dried with anhydrous magnesium sulfate, filtered, and concentrated *in vacuo*. The crude residue was purified by silica gel column chromatography (0-30% EtOAc/hexanes, linear gradient) to afford 2 (2.50 g, 50%) as a white solid.

$^1$H NMR (400 MHz, $CDCl_3$) δ 8.49 (d, $J$ = 8.8 Hz, 2H), 7.63 (d, $J$ = 2.4 Hz, 2H), 7.42 (dd, $J$ = 8.8, 2.4 Hz, 2H), 1.82 (s, 6H).

HRMS (EI) calcd for $C_{18}H_{12}F_6O_7S_2$ $[M\cdot]^+$ 517.9929, found 517.9922.

The NMR and HRMS agreed with reported values.[9]

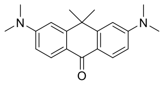

*3,6-bis(dimethylamino)-10,10-dimethylanthracen-9(10H)-one (3):*

A bomb flask was charged with ditriflate **2** (400 mg, 0.772 mmol), $Pd_2dba_3$ (71 mg, 0.077 mmol, 0.1 eq), XPhos (110 mg, 0.232 mmol, 0.3 eq), and $Cs_2CO_3$ (704 mg, 2.16 mmol, 2.8 eq) and sealed with a rubber septum. The flask was evacuated and backfilled with nitrogen (3x). Dimethylamine (2 M in THF, 7.72 mL, 15.4 mmol, 20 eq) was added, the septum replaced with a Teflon screw top, and the sealed reaction stirred at 100 °C for 4 h. It was then cooled to room temperature, diluted with $CH_2Cl_2$, and filtered. The reaction mixture was deposited onto Celite and concentrated to dryness. Silica gel chromatography purification (0-30% EtOAc/hexanes w/ 20% $CH_2Cl_2$ & 1% $NEt_3$) afforded 229 mg (96%) of **3** as a yellow solid.

$^1$H NMR (400 MHz, $CDCl_3$) δ 8.31 (d, *J* = 8.6 Hz, 2H), 6.84 – 6.77 (m, 4H), 3.12 (s, 12H), 1.76 (s, 6H);

$^{13}$C NMR (101 MHz, $CDCl_3$) δ 181.2, 153.1, 152.3, 129.1, 119.9, 110.8, 107.8, 40.2, 38.2, 33.8;

HRMS (ESI) calcd for $C_{20}H_{25}N_2O$ $[M+H]^+$ 309.1961, found 309.1959.

The NMR and HRMS agreed with reported values.[10]

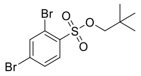

*neopentyl 2,4-dibromobenzenesulfonate (6):*

A solution of 2,4-dibromobenzenesulfonyl chloride[2] (5.46 g, 16.33 mmol) and neopentyl alcohol (3.60 g, 40.8 mmol, 2.5 eq) in $CH_2Cl_2$ (65 mL) was cooled to 0 °C, and 1,4-Diazabicyclo[2.2.2]octane (4.58 g, 40.8 mmol, 2.5 eq) was added. The reaction was stirred at room temperature for 30 min. The reaction mixture was then filtered, washed several times with $CH_2Cl_2$, and concentrated *in vacuo*. The crude residue was purified by flash chromatography on silica gel (0-15% EtOAc/hexanes, linear gradient, dry load on silica) to afford **5** (4.32 g, 69%) as a white solid.

$^1$H NMR (400 MHz, $CDCl_3$) δ 8.00 (d, *J* = 4.8 Hz, 1H), 7.99 (d, *J* = 1.7 Hz, 1H), 7.67 (dd, *J* = 8.5, 1.9 Hz, 1H), 3.77 (s, 2H), 1.00 (s, 9H);

$^{13}$C NMR (101 MHz, $CDCl_3$) δ 138.0, 135.1, 133.0, 130.9, 128.8, 121.7, 80.7, 31.8, 26.1;

HRMS (EI) calcd for $C_{11}H_{14}Br_2O_3S$ $[M\cdot]^+$ 383.9030, found 383.9034.

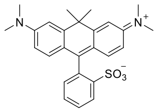

*2-(6-(dimethylamino)-3-(dimethyliminio)-10,10-dimethyl-3,10-dihydroanthracen-9-yl)benzenesulfonate (7):*

In flask A, ketone **3** (100 mg, 0.324 mmol) was dissolved in $CH_2Cl_2$ (1.0 mL) and a drop of $Et_3N$ was added. Trifluoromethanesulfonic anhydride (100 mg, 0.357 mmol, 1.1 eq) was added and the reaction stirred for 30 min. In flask B, a solution of neopentyl 2-bromobenzenesulfonate **4**[3] (298 mg, 0.97 mmol, 3.0 eq) dissolved in $CH_2Cl_2$ (2.4 mL) under nitrogen was cooled to -16 °C using a salt-ice bath. *n*-Butyllithium (1.6 M in hexanes, 0.61 mL, 0.973 mmol, 3.0 eq) was added slowly, and the reaction was stirred at -16 °C for 15 min. Then, the solution from flask A was added slowly to flask B at -16 °C, warmed to room temperature, and stirred for 1-2 h. The reaction was quenched with MeOH and purified by preparative TLC (10% MeOH/$CH_2Cl_2$). The blue bands were collected, the solid dissolved in $CH_2Cl_2$ (1 mL) and subjected to 1.0 M $BBr_3$ (1 mL). The reaction mixture was stirred at room temperature overnight and purified by PTLC (10% MeOH/$CH_2Cl_2$) to afford **7** (27 mg, 19%) as a blue solid.

$^1$H NMR (400 MHz, 10:1 $CDCl_3$/$CD_3OD$ v/v) δ 8.23 (d, *J* = 7.9 Hz, 1H), 7.57 (t, *J* = 7.6 Hz, 1H), 7.47 (t, *J* = 7.4 Hz, 1H), 7.19 (d, *J* = 9.3 Hz, 2H), 7.05 (d, *J* = 7.5 Hz, 1H), 6.93 (d, *J* = 2.5 Hz, 2H), 6.61 (dd, *J* = 9.3, 2.5 Hz, 2H), 3.24 (s, 12H), 1.77 (s, 3H), 1.68 (s, 3H);

$^{13}$C NMR (101 MHz, Chloroform-*d*) δ 168.0, 157.0, 156.2, 145.2, 139.1, 132.6, 129.3, 129.2, 128.9, 128.6, 121.7, 112.2, 109.7, 41.8, 40.6, 35.6, 32.0;

HRMS (ESI) calcd for $C_{26}H_{28}N_2NaO_3S$ $[M+Na]^+$ 471.1713, found 471.1707.

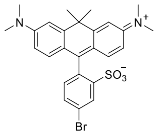

*5-bromo-2-(6-(dimethylamino)-3-(dimethyliminio)-10,10-dimethyl-3,10-dihydroanthracen-9-yl)benzenesulfonate (8):*

In flask A, ketone **3** (101 mg, 0.327 mmol) was dissolved in $CH_2Cl_2$ (1.0 mL) under nitrogen and a drop of $Et_3N$ was added. Trifluoromethanesulfonic anhydride (102 mg, 0.360 mmol, 1.1 eq) was added and the reaction stirred for 30 min. In flask B, a solution of neopentyl 2,5-dibromobenzenesulfonate **5**[3] (379 mg, 0.982 mmol, 3.0 eq) dissolved in $CH_2Cl_2$ (2.5 mL) under nitrogen was cooled to -16 °C using a salt-ice bath. *n*-Butyllithium (1.6 M in hexanes, 0.61 mL, 0.98 mmol, 3.0 eq) was added slowly, and the reaction was stirred at -16 °C for 15 min. Then, the solution from flask A was added slowly to flask B at -16 °C, warmed to room temperature, and stirred for 1-2 h. The reaction was quenched with MeOH and purified by preparative TLC (10% MeOH/$CH_2Cl_2$). The blue bands were collected and subjected to 1.0 M $BBr_3$ (2 mL). The reaction mixture was stirred at room temperature overnight and purified by PTLC (10% MeOH/$CH_2Cl_2$) to afford **8** (29 mg, 17%) as a blue solid.

$^1$H NMR (400 MHz, $CDCl_3$) δ 8.53 (d, *J* = 1.9 Hz, 1H), 7.59 (dd, *J* = 8.1, 2.0 Hz, 1H), 7.28 (d, *J* = 9.3 Hz, 2H), 7.02 – 6.92 (m, 3H), 6.62 (dd, *J* = 9.3, 2.2 Hz, 2H), 3.26 (s, 12H), 1.83 (s, 3H), 1.68 (s, 3H);

$^{13}$C NMR (101 MHz, $CDCl_3$) δ 167.4, 157.0, 156.1, 148.7, 139.0, 132.2, 131.4, 131.3, 130.5, 123.2, 122.0, 112.4, 110.0, 42.1, 40.9, 36.1, 31.4;

HRMS (ESI) calcd for $C_{26}H_{27}BrN_2NaO_3S$ $[M+Na]^+$ 549.0818, found 549.0816.

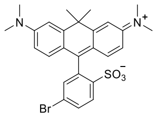

*4-bromo-2-(6-(dimethylamino)-3-(dimethyliminio)-10,10-dimethyl-3,10-dihydroanthracen-9-yl)benzenesulfonate (9):*

In flask A, ketone **3** (93 mg, 0.29 mmol) was dissolved in $CH_2Cl_2$ (1.0 mL) and a drop of $Et_3N$ was added. Trifluoromethanesulfonic anhydride (92 mg, 0.32 mmol, 1.1 eq) was added and the reaction stirred for 30 min. In flask B, a solution of neopentyl 2,4-dibromobenzenesulfonate **6** (375 mg, 0.973 mmol, 3.4 eq) dissolved in $CH_2Cl_2$ (2.1 mL) under nitrogen was cooled to -16 °C using a salt-ice bath. *n*-Butyllithium (1.6 M in hexanes, 0.61 mL, 0.97 mmol, 3.4 eq) was added slowly, and the reaction was stirred at -16 °C for 15 min. Then, the solution from flask A was added slowly to flask B at -16 °C, warmed to room temperature, and stirred for 1-2 h. The reaction was quenched with MeOH and purified by preparative TLC (10% MeOH/$CH_2Cl_2$). The blue bands were collected and subjected to 1.0 M $BBr_3$ (2 mL). The reaction mixture was stirred at room temperature overnight and purified by PTLC (10% MeOH/$CH_2Cl_2$) to afford **9** (20 mg, 13%) as a blue solid.

$^1$H NMR (400 MHz, $CD_3OD$) δ 8.07 (d, *J* = 8.5 Hz, 1H), 7.85 (dd, *J* = 8.5, 2.1 Hz, 1H), 7.42 (d, *J* = 2.0 Hz, 1H), 7.20 (d, *J* = 2.6 Hz, 2H), 7.12 (s, 1H), 7.10 (s, 1H), 6.81 (dd, *J* = 9.4, 2.5 Hz, 2H), 3.34 (s, 12H), 1.86 (s, 3H), 1.78 (s, 3H);

$^{13}$C NMR (101 MHz, $CD_3OD$) δ 163.4, 157.1, 156.5, 143.7, 137.9, 135.5, 132.3, 132.0, 129.7, 123.4, 121.0, 112.2, 110.3, 41.8, 39.5, 34.3, 31.3;

HRMS (ESI) calcd for $C_{26}H_{28}BrN_2O_3S$ $[M+H]^+$ 527.0999, found 527.1000.

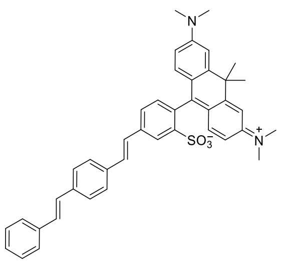

*2-(6-(dimethylamino)-3-(dimethyliminio)-10,10-dimethyl-3,10-dihydroanthracen-9-yl)-5-((E)-4-((E)-styryl)styryl)benzenesulfonate (13):*

A vial was charged with 8 (8.0 mg, 15.2 µmol, 1.0 eq), styrene 10[4] (3.5 mg, 16.7 µmol, 1.1 eq), $Pd(OAc)_2$ (1.4 mg, 6.07 µmol, 0.4 eq), and P(*o*-tol)$_3$ (3.7 mg, 12.1 µmol, 0.8 eq). The vial was sealed and evacuated/backfilled with nitrogen (3x). Anhydrous DMF (220 µL) was added and the vial was evacuated/backfilled again with nitrogen (3x). Anhydrous $Et_3N$ (110 µL) was added and the reaction was stirred at 90 °C for 18 h. It was then cooled, diluted with $CH_2Cl_2$, and filtered through a cotton plugged glass pipette. After filtration, the solvent was removed under reduced pressure. The crude residue was purified by preparative TLC (10% MeOH/$CH_2Cl_2$) to afford 13 (7 mg, 70%) as a dark green solid.

$^1$H NMR (900 MHz, 10:1 $CDCl_3$/$CD_3OD$ v/v) δ 8.38 (d, *J* = 1.8 Hz, 1H), 7.57 (dd, *J* = 7.8, 1.8 Hz, 1H), 7.52 – 7.47 (m, 5H), 7.31 (t, *J* = 7.6 Hz, 2H), 7.28 – 7.24 (m, 4H), 7.22 – 7.18 (m, 2H), 7.12 – 7.06 (m, 2H), 7.02 (d, *J* = 7.7 Hz, 1H), 6.90 (d, *J* = 2.6 Hz, 2H), 6.60 (dd, *J* = 9.4, 2.5 Hz, 2H), 3.22 (s, 12H), 1.74 (s, 6H);

$^{13}$C NMR (226 MHz, 10:1 $CDCl_3$/$CD_3OD$ v/v) δ 168.2, 157.1, 156.4, 145.9, 139.3, 138.9, 137.4, 137.3, 136.4, 131.6, 130.5, 129.9, 128.9, 128.8, 128.3, 127.8, 127.3, 127.0, 127.0, 126.7, 126.7, 126.6, 121.9, 112.4, 109.8, 42.0, 40.7, 35.7, 32.2;

HRMS (ESI) calcd for $C_{42}H_{40}N_2O_3S$ $[M+Na]^+$ 675.2652, found 675.2648.

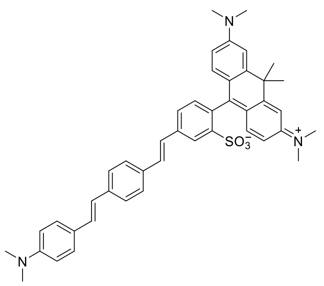

*2-(6-(dimethylamino)-3-(dimethyliminio)-10,10-dimethyl-3,10-dihydroanthracen-9-yl)-5-((E)-4-((E)-4-(dimethylamino)styryl)styryl)benzenesulfonate (14):*

A vial was charged with 8 (11 mg, 20.8 μmol, 1.0 eq), styrene 11[5] (5 mg, 20.8 μmol, 1.0 eq), $Pd(OAc)_2$ (2.3 mg, 10.4 μmol, 0.5 eq), and P(*o*-tol)$_3$ (6.4 mg, 20.8 μmol, 1.0 eq). The vial was sealed and evacuated/backfilled with nitrogen (3x). Anhydrous DMF (230 μL) was added and the vial was evacuated/backfilled again with nitrogen (3x). Anhydrous $Et_3N$ (110 μL) was added and the reaction was stirred at 75 °C for 1 h. It was then cooled, diluted with $CH_2Cl_2$, and filtered through a cotton plugged glass pipette. After filtration, the solvent was removed under reduced pressure. The crude residue was purified by preparative TLC (10% MeOH/$CH_2Cl_2$) to afford 14 (5.2 mg, 36%) as a dark green solid.

$^1$H NMR (900 MHz, 10:1 $CDCl_3$/$CD_3OD$ v/v) δ 8.36 (d, *J* = 1.8 Hz, 1H), 7.56 (dd, *J* = 7.9, 1.8 Hz, 1H), 7.48 (d, *J* = 7.9 Hz, 2H), 7.44 (d, *J* = 8.1 Hz, 2H), 7.38 (d, *J* = 8.5 Hz, 2H), 7.26 – 7.24 (m, 3H), 7.16 (d, *J* = 16.4 Hz, 1H), 7.04 (d, *J* = 16.2 Hz, 1H), 7.01 (d, *J* = 7.7 Hz, 1H), 6.90 – 6.87 (m, 3H), 6.69 (d, *J* = 8.1 Hz, 2H), 6.61 (dd, *J* = 9.4, 2.5 Hz, 2H), 3.22 (s, 12H), 2.93 (s, 6H), 1.74 (s, 3H), 1.67 (s, 3H);

$^{13}$C NMR (226 MHz, 10:1 $CDCl_3$/$CD_3OD$ v/v) δ 168.3, 157.1, 156.4, 150.4, 145.8, 139.3, 139.1, 138.3, 135.4, 131.5, 130.7, 129.9, 129.4, 127.8, 127.2, 126.6, 126.5, 126.5, 126.4, 124.1, 122.0, 112.8, 112.4, 109.8, 42.0, 40.7, 40.6, 35.7, 32.2;

HRMS (ESI) calcd for $C_{44}H_{46}N_3O_3S$ $[M+H]^+$ 696.3254, found 696.3247.

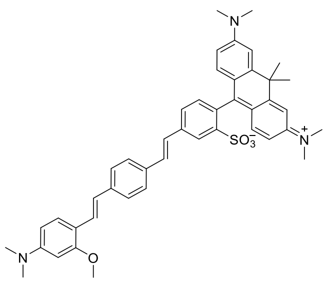

*5-((E)-4-((E)-4-(dimethylamino)-2-methoxystyryl)styryl)-2-(6-(dimethylamino)-3-(dimethyliminio)-10,10-dimethyl-3,10-dihydroanthracen-9-yl)benzenesulfonate (15):*

A vial was charged with 8 (20 mg, 37.9 μmol, 1.0 eq), styrene 12[4] (11 mg, 37.9 μmol, 1.0 eq), $Pd(OAc)_2$ (4.3 mg, 19.0 μmol, 0.5 eq), and P(*o*-tol)$_3$ (11 mg, 37.9 μmol, 1.0 eq). The vial was sealed and evacuated/backfilled with nitrogen (3x). Anhydrous DMF (410 μL) was added and the vial was evacuated/backfilled again with nitrogen (3x). Anhydrous $Et_3N$ (210 μL) was added and the reaction was stirred at 75 °C for 1 h. It was then cooled, diluted with $CH_2Cl_2$, and filtered through a cotton plugged glass pipette. After filtration, the solvent was removed under reduced pressure. The crude residue was purified by preparative TLC (10% MeOH/$CH_2Cl_2$) to afford 15 (9.2 mg, 34%) as a dark green solid.

$^1$H NMR (900 MHz, 10:1 $CDCl_3$/$CD_3OD$ v/v) δ 8.36 (d, *J* = 1.8 Hz, 1H), 7.56 (dd, *J* = 7.9, 1.8 Hz, 1H), 7.48 – 7.44 (m, 4H), 7.43 (d, *J* = 8.4 Hz, 1H), 7.39 (d, *J* = 16.4 Hz, 1H), 7.26 – 7.23 (m, 3H), 7.15 (d, *J* = 16.3 Hz, 1H), 7.00 (d, *J* = 7.7 Hz, 1H), 6.93 – 6.89 (m, 3H), 6.60 (dd, *J* = 9.4, 2.5 Hz, 2H), 6.32 (d, *J* = 8.3 Hz, 1H), 6.20 (s, 1H), 3.85 (s, 3H), 3.22 (s, 12H), 2.95 (s, 6H), 1.74 (s, 3H), 1.66 (s, 3H);

$^{13}$C NMR (226 MHz, 10:1 $CDCl_3$/$CD_3OD$ v/v) δ 168.2, 158.1, 156.9, 156.2, 151.5, 145.6, 139.1, 138.9, 138.8, 134.9, 131.2, 130.6, 129.6, 127.2, 126.9, 126.3, 126.3, 125.9, 124.2, 123.8, 121.8, 112.2, 109.6, 105.2, 95.7, 55.4, 41.8, 40.5, 40.4, 35.5, 32.0;

HRMS (ESI) calcd for $C_{45}H_{48}N_3O_4S$ $[M+H]^+$ 726.3360, found 726.3347.

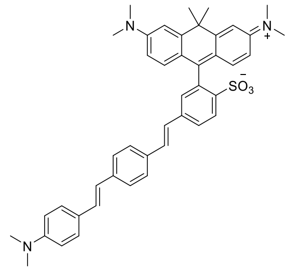

*2-(6-(dimethylamino)-3-(dimethyliminio)-10,10-dimethyl-3,10-dihydroanthracen-9-yl)-4-((E)-4-((E)-4-(dimethylamino)styryl)styryl)benzenesulfonate (16):*

A vial was charged with **9** (5.5 mg, 10.4 μmol, 1.0 eq), styrene 11[5] (2.8 mg, 11.2 μmol, 1.1 eq), $Pd(OAc)_2$ (0.94 mg, 4.19 μmol, 0.4 eq), and P(*o*-tol)$_3$ (2.5 mg, 8.21 μmol, 0.8 eq). The vial was sealed and evacuated/backfilled with nitrogen (3x). Anhydrous DMF (94 μL) was added and the vial was evacuated/backfilled again with nitrogen (3x). Anhydrous $Et_3N$ (47 μL) was added and the reaction was stirred at 75 °C for 3 h. It was then cooled, diluted with $CH_2Cl_2$, and filtered through a cotton plugged glass pipette. After filtration, the solvent was removed under reduced pressure. The crude residue was purified by preparative TLC (10% MeOH/$CH_2Cl_2$) and then triturated with MeOH to afford **16** (3.7 mg, 51%) as a dark green solid.

$^1$H NMR (900 MHz, 10:1 $CDCl_3$/$CD_3OD$ v/v) δ 8.17 (d, *J* = 8.3 Hz, 1H), 7.68 (dd, *J* = 8.3, 1.8 Hz, 1H), 7.40 (q, *J* = 8.3, 7.6 Hz, 6H), 7.26 (m, 3H), 7.15 (d, *J* = 1.8 Hz, 1H), 7.09 (d, *J* = 16.2 Hz, 1H), 7.06 – 7.00 (m, 3H), 6.91 (d, *J* = 2.5 Hz, 2H), 6.62 (dd, *J* = 9.5, 2.5 Hz, 3H), 3.23 (s, 12H), 2.98 (s, 6H), 1.76 (s, 3H), 1.67 (s, 3H);

$^{13}$C NMR (226 MHz, Chloroform-*d*) δ 167.9, 157.1, 156.4, 144.2, 139.3, 133.3, 130.7, 129.2, 127.9, 127.9, 127.2, 127.2, 127.1, 126.7, 121.8, 112.5, 109.9, 42.0, 40.7, 35.7, 32.2;

HRMS (ESI) calcd for $C_{44}H_{46}N_3O_3S$ $[M+H]^+$ 696.3254, found 696.3251.

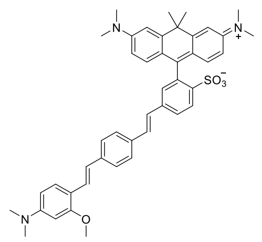

*4-((E)-4-((E)-4-(dimethylamino)-2-methoxystyryl)styryl)-2-(6-(dimethylamino)-3-(dimethyliminio)-10,10-dimethyl-3,10-dihydroanthracen-9-yl)benzenesulfonate (17):*

A vial was charged with 9 (4.4 mg, 8.34 µmol, 1.0 eq), styrene 12[4] (2.6 mg, 9.18 µmol, 1.1 eq), $Pd(OAc)_2$ (0.94 mg, 4.17 µmol, 0.5 eq), and P(*o*-tol)$_3$ (2.5 mg, 8.34 µmol, 1.0 eq). The vial was sealed and evacuated/backfilled with nitrogen (3x). Anhydrous DMF (91 µL) was added and the vial was evacuated/backfilled again with nitrogen (3x). Anhydrous $Et_3N$ (45 µL) was added and the reaction was stirred at 75 °C for 3 h. It was then cooled, diluted with $CH_2Cl_2$, and filtered through a cotton plugged glass pipette. After filtration, the solvent was removed under reduced pressure. The crude residue was purified by preparative TLC (10% MeOH/$CH_2Cl_2$) and then triturated with MeOH to afford 17 (3 mg, 50%) as a dark green solid.

$^1$H NMR (900 MHz, 10:1 $CDCl_3$/$CD_3OD$ v/v) δ 8.17 (dd, *J* = 8.3, 3.5 Hz, 1H), 7.69 (dd, *J* = 8.3, 1.7 Hz, 1H), 7.56 (d, *J* = 27.6 Hz, 1H), 7.47 – 7.39 (m, 4H), 7.35 (d, *J* = 16.4 Hz, 1H), 7.25 (m, 3H), 7.16 (d, *J* = 1.8 Hz, 1H), 7.10 (d, *J* = 16.3 Hz, 2H), 7.08 – 7.03 (m, 2H), 6.92 (d, *J* = 2.6 Hz, 2H), 6.62 (dt, *J* = 9.4, 2.5 Hz, 2H), 3.89 (s, 3H), 3.23 (s, 12H), 3.10 (s, 6H), 1.77 (s, 3H), 1.67 (s, 3H);

$^{13}$C NMR (226 MHz, 10:1 $CDCl_3$/$CD_3OD$ v/v) δ 167.9, 158.0, 158.0, 157.2, 156.4, 156.4, 144.3, 139.3, 139.3, 138.2 133.3, 130.7, 130.7, 129.3, 127.6, 127.3, 127.2, 127.1, 126.7, 121.9, 112.5, 109.9, 42.0, 40.75, 35.7, 32.2;

HRMS (ESI) calcd for $C_{45}H_{48}N_3O_4S$ $[M+H]^+$ 726.3360, found 726.3348.

### Supporting Tables

**Table S1.** Spatiotemporal resolution of fluorescence lifetime imaging data.

| Application | Raw Frame Acquisition Time (s) | Binned Frame Acquisition Time (s) | Number of Time Points (Total Time Series Duration) | Image Size (width x height, μm$^2$) | Pixel Size of Lifetime Image (binned, μm$^2$) |
|---|---|---|---|---|---|
| Single Image | 0.94 | 75-90 | 1 (75-90 s) | 135 x 135 | 2.1 x 2.1 |
| Electrophysiology | 0.24 | 15 | 5 (15 s) | 53 x 53 | 1.7 x 1.7 |
| EGF Treatment | 0.63 | 5 | 36 (180 s), 177 (900 s) | 135 x 33.7 | 6.3 x 6.3 * |
| iCM Spontaneous Activity | 0.025 | 0.050 | 200 (10 s) | 135 x 18 | N/A (global analysis) |

"Single image" data includes concentration curve loading tests and gramicidin treatment in cell lines (**Figure S7 and S9**). EGF time series are shown in **Figure 4 and Figure S11**. iCM spontaneous activity data are shown in **Figure 5 and Figures S12-S13**. Raw frame acquisition time reflects the frame rate at which data were captured; successive frames were combined for later analysis to obtain good photon statistics. The number of time points is the number of analyzed, binned frames.

*For EGF treatment, pixel size after standard binning is the same as single images (2.1 x 2.1 μm$^2$). To improve photon statistics for pixelwise fitting, moving average binning was performed during analysis, as is common for Becker & Hickl FLIM products (e.g. SPCImage). Therefore, the lifetimes at each pixel were determined by photons from a 6.3 x 6.3 μm$^2$ area centered on the displayed pixel. The photon count image is displayed at the native resolution (2.1 μm per side of pixel).

### Supporting Figures

**Figure S1.** Characterization of voltage sensitivities of other carborhodamine voltage indicators.

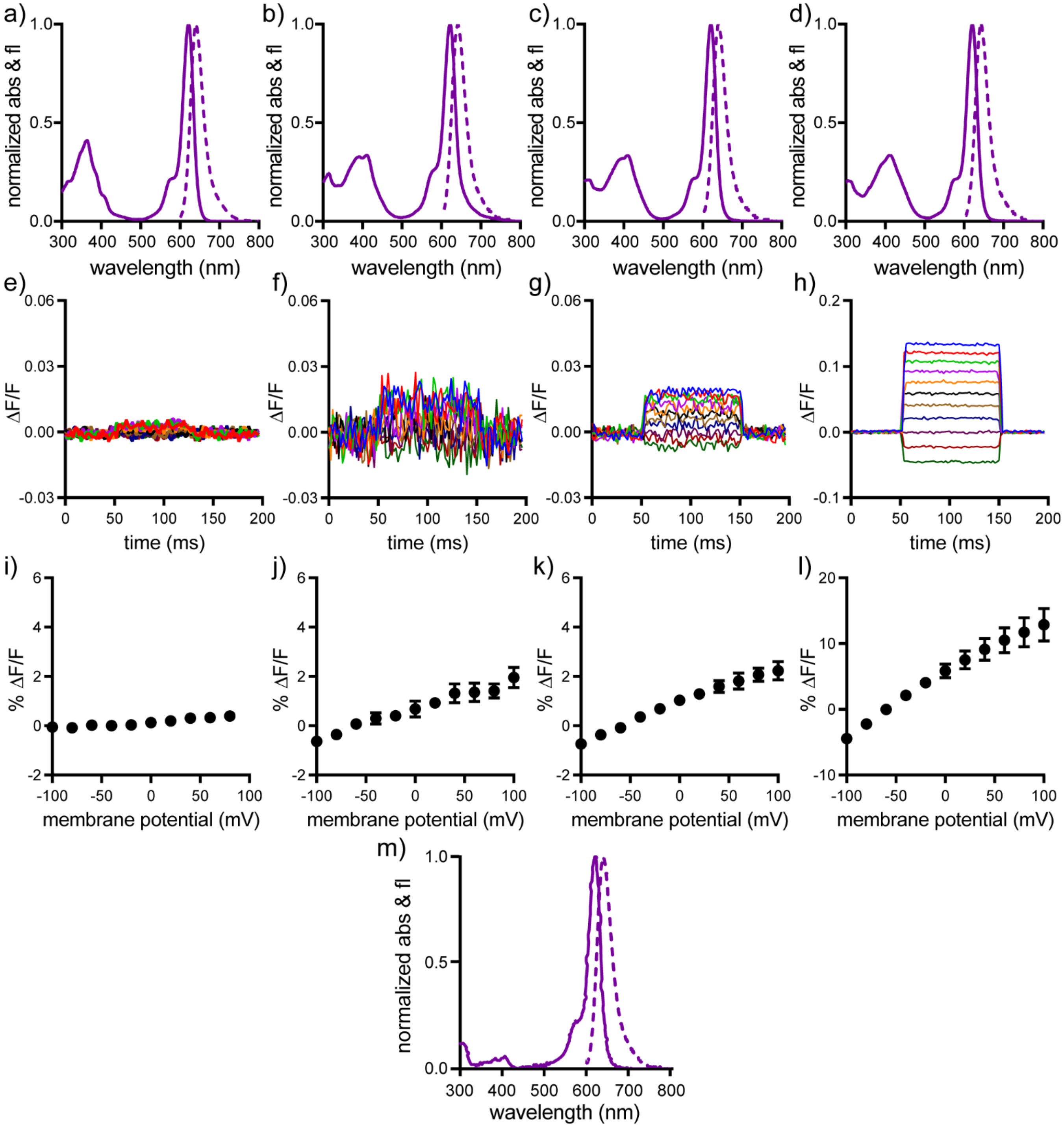

Normalized absorbance and emission of a) **13**, b) **14**, c) **16**, d) **17** and m) TMCRh-sulfonate **7** in PBS, pH 7.4, 0.1% SDS. Voltage sensitivity of TMCRh dyes. e,i) **13**; f,j) **14**; g,k) **16**; and h,l) **17**. e-h: Plots of fractional change in fluorescence (ΔF/F) vs time for HEK cells held at -60 mV and stepped to 100 ms hyper- and depolarizing steps in 20 mV increments (±100 mV) under whole-cell voltage-clamp conditions. i-l: Plots of % ΔF/F vs. final membrane potential (mV) for n = 4-7 cells for each TMCRh dye. Error bars are ± S.D.

**Figure S2.** Cell loading and brightness comparison of TMCRh voltage indicators.

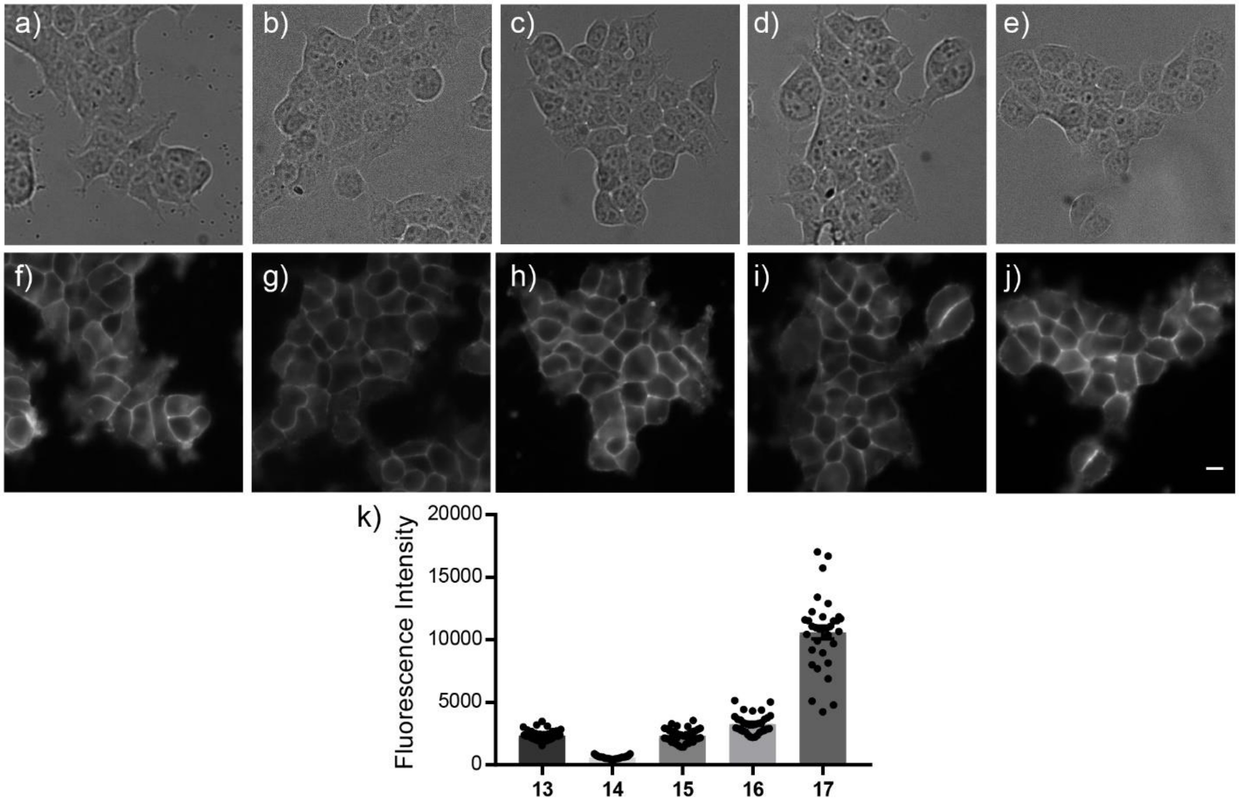

**a-e**) Differential interference contrast (DIC) images of human embryonic kidney (HEK) cells loaded with TMCRh dyes (500 nM, 15 min) then washed with fresh HBSS. **f-j**) Wide-field fluorescence images of cells stained with TMCRh dyes with brightness enhanced for panels **f-i** to enable visual inspection of cellular localization. Labeling of HEK cells with respective dyes is as follows: a,f) **13** b,g) **14** c,h) **15** d,i) **16** e,j) **17**. **k**) Plot of relative brightness of TMCRh dyes loaded in HEK cells (n = 4 coverslips). Each individual point represents the average brightness of HEK cells in a single image. Error bars are ± S.E.M.

**Figure S3.** Field stimulation of neurons stained with CRhOMe (**15**).

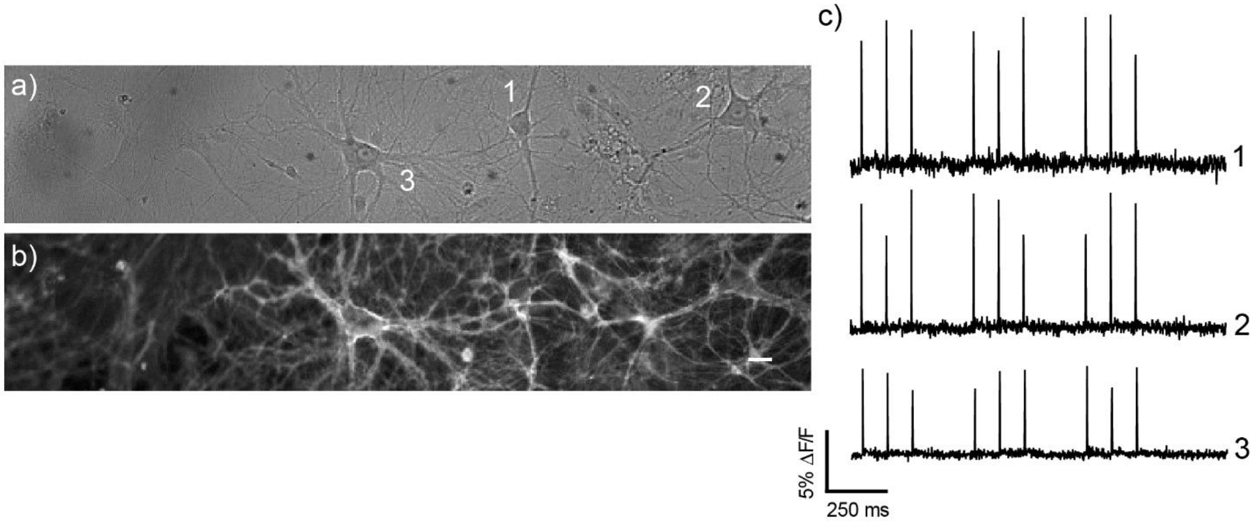

Rat hippocampal neurons (DIC, a) were imaged with 500 nM CRhOMe (fluorescence, b)**.** c) Electrode-evoked activity (1.85 $W/cm^2$) from the neurons in a-b) are shown as $\Delta F/F$ traces. Scale bar is 20 μm.

**Figure S4.** Comparison of CRhOMe (**15**) and BeRST 1 in neurons.

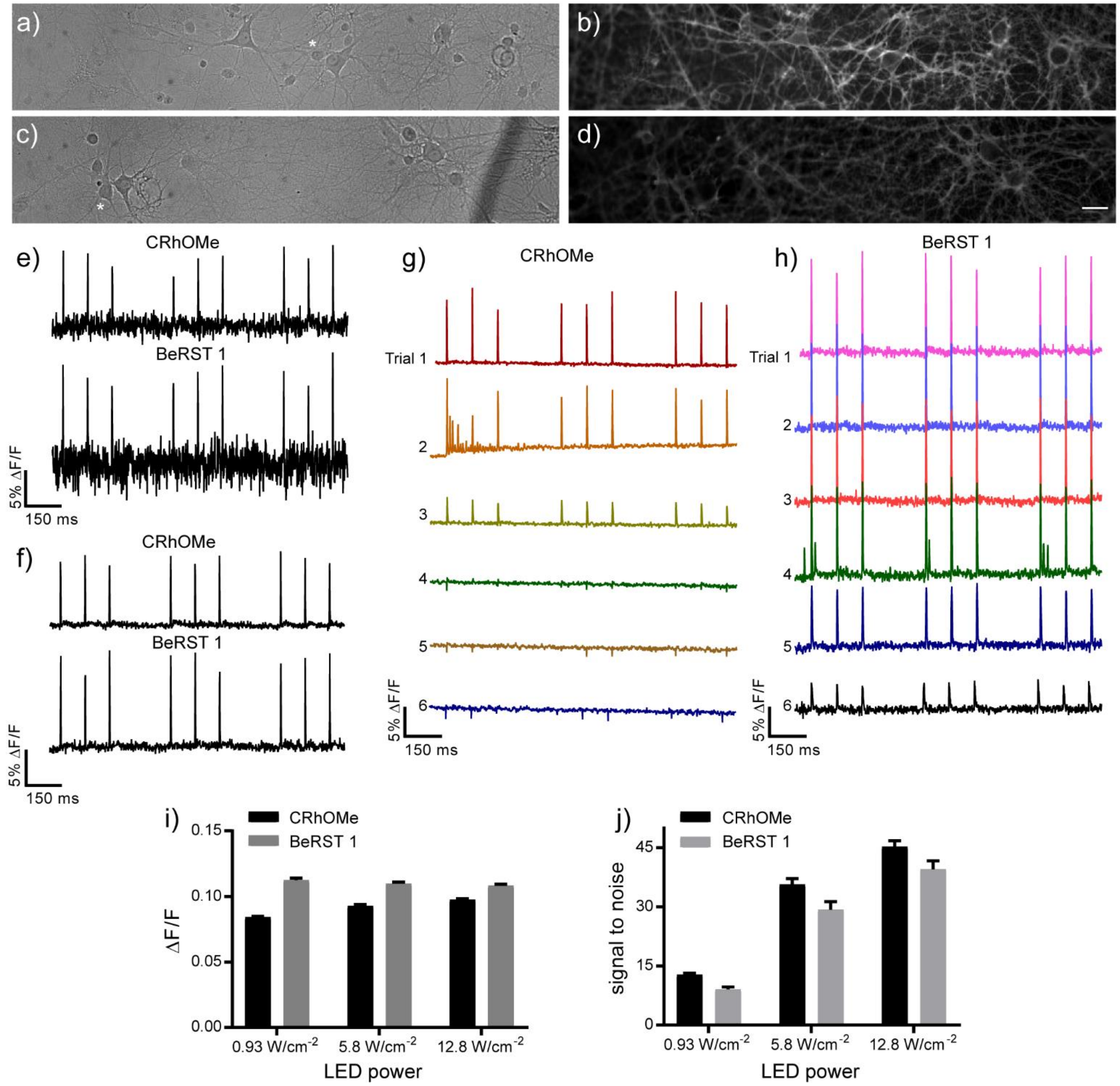

a-d) Rat hippocampal neurons (DIC, a) imaged with 500 nM CRhOMe (fluorescence, b). Neurons (DIC, c) imaged with 500 nM BeRST 1 (fluorescence, d) and normalized to CRhOMe fluorescence**.** Scale bar is 20 μm. e) Electrode-evoked activity from neurons loaded with either CRhOMe or BeRST 1 at 500 nM for 15 min and imaged using 0.93 $W/cm^2$ light intensity is shown as ΔF/F traces. f) Electrode-evoked activity recordings using 5.78 $W/cm^2$ light power (these data are shown in **Figure 2e** in the main text). g) ΔF/F traces acquired using 12.8 $W/cm^2$ light intensity of the neuron highlighted in panel a stained with CRhOMe. Top to bottom: Repeated applications of light cause the neuron to stop responding to field stimulation. h) ΔF/F traces acquired using 12.8 $W/cm^2$ light intensity of the neuron highlighted in panel c stained with BeRST 1. Top to bottom: Repeated applications of light cause the neuron to stop respond to field stimulation with a lower ΔF/F. For panels g and h, Trials 1-3 represent a 5 second recording. For Trial 4, 5, and 6, 30 seconds of light illumination was applied immediately before recording the 5 second movie. i-j) Comparison of the average ΔF/F or SNR per spike for CRhOMe (*n* = 21-50 cells) and BeRST 1 (*n* = 21-24 cells) using different light intensities with $\lambda_{ex}$ centered at 633 nm. Summaries of these values are in **Figure 2f** in the main text.

**Figure S5.** Lifetime decay of CRhOMe, BeRST, and TMCRh fluorophore

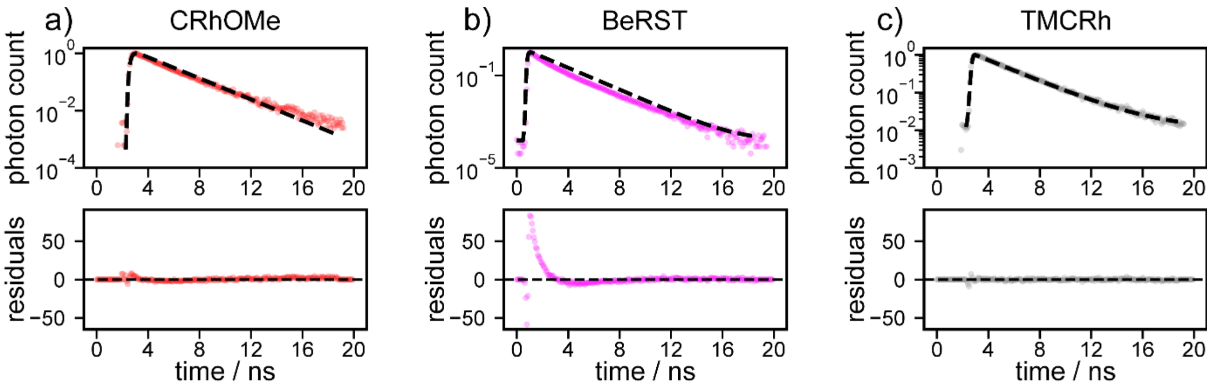

Plots of fluorescence lifetime decays for **a)** CRhOMe (red), **b)** BeRST 1 (magenta), and **c)** TMCRh fluorophore (**7**) (grey). Dashed black line is the line of best fit for a monoexponential decay. Upper panels are normalized photon counts per second vs. time. Lower panels are weighted residuals vs. time. Panel (**a**) is repeated from Figure 3a in the main text, for comparison purposes.

**Figure S6.** Distributions of mean lifetimes and resting potentials across FLIM systems

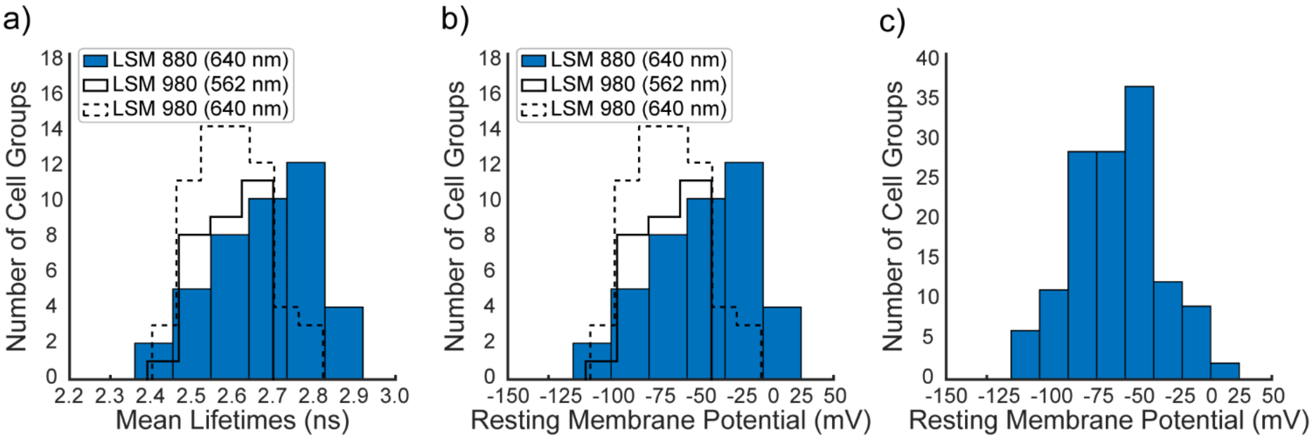

Histograms representing the mean lifetimes of CRhOMe and resting membrane potentials of HEK293T cells across both FLIM systems used. a) Mean lifetimes (ns) plotted against the frequency (in number of cell groups) for both the LSM 880 system at 640 nm (blue) and LSM 980 system at 562 nm (solid line) or 640 nm (dashed line). b) The resting membrane potentials were determined by applying the calibration determined (Figure 3 in main text) to the lifetimes in a. c) Combined resting membrane potentials from both systems.

**Figure S7.** Fluorescence lifetime vs. concentration for CRhOMe and BeRST in different cell lines.

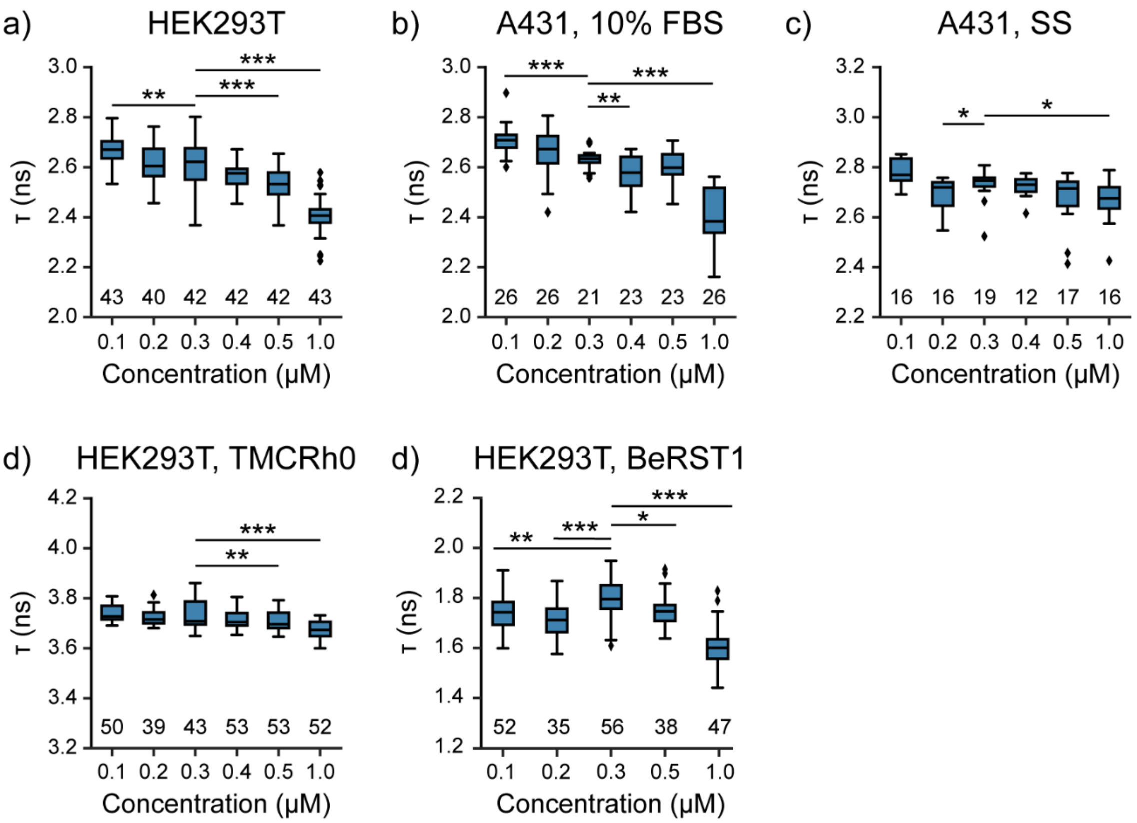

Plots of fluorescence lifetime versus CRhOMe concentration in cells. a) CRhOMe in HEK293T cells. b) CRhOMe in A431 cells. c) CRhOMe in serum starved A431 cells. d) TMCRh0 (aka TMCRhZero) in HEK293T cells. e) BeRST1 dye in HEK 293T cells. Significance levels are reported as compared to 300 nM dye, and are the result of Kruskal-Wallis tests, and Dunn post-hoc analysis. * = $p < 0.05$, ** = $p < 0.01$, *** = $p < 0.001$. The y axis range for all plots is 1 ns.

**Figure S8.** Comparison of voltage and lifetime response of TMCRhZero dye in HEK293T cells.

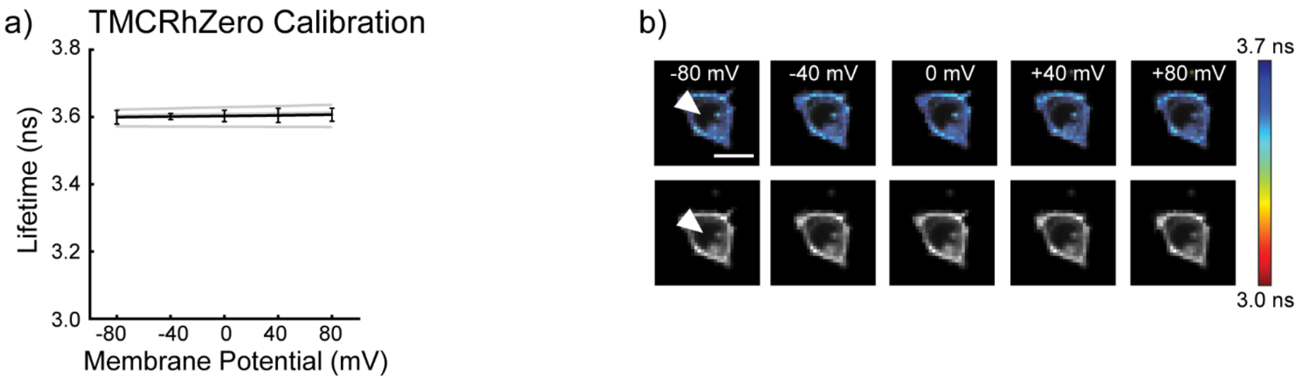

a) Plot of lifetime vs. membrane potential for TMCRhZero in HEK cells. Gray lines indicate individual cell responses (n = 3), and the black line indicates the overall average response, with a sensitivity (slope) of $4.5 \times 10^{-5}$ ps/mV, and a lifetime at 0 mV of 3.6 ns. b) Fluorescence lifetime and intensity responses in HEK293T cells when voltage clamped at the indicated voltages. Lifetime heatmap scaled from 3.0 to 3.7 ns. Scalebar 20 µm.

**Figure S9.** Lifetime of CRhOMe in cells at rest and depolarized with gramicidin.

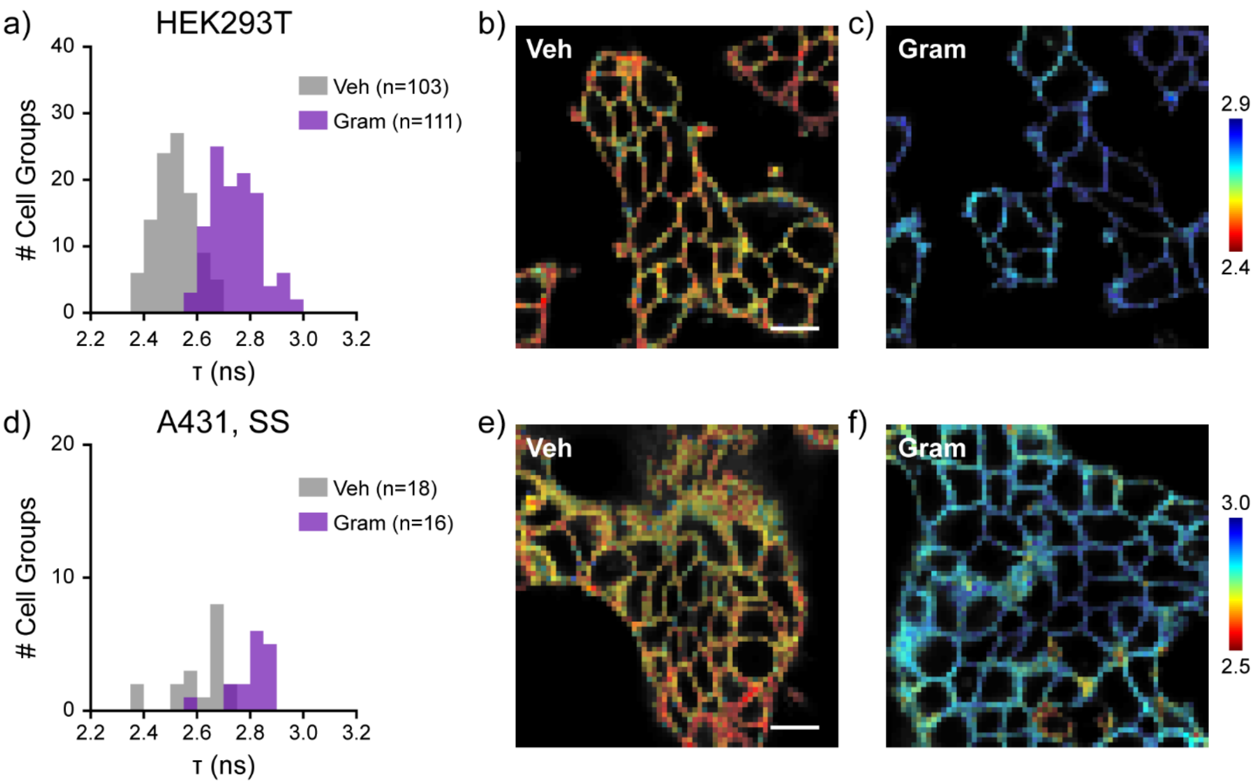

a) Histogram of lifetimes observed for 300 nM CRhOMe in groups of HEK293T at rest (vehicle, Veh, 0.05-0.1% DMSO) and treated with 500 ng/mL gramicidin (Gram). b) Lifetime-photon count overlay image of HEK293T at rest and c) treated with 500 ng/mL gramicidin. d) Histogram of CRhOMe lifetimes observed in groups of serum starved (SS) A431 cells at rest and depolarized with gramicidin. e-f). Lifetime images of SS A431 at rest and gramicidin-treated, as in (b)-(c). Bin sizes were determined by the Freedman-Diaconis rule. Scale bars represent 20 µm. Photon counts of lifetime-photon count overlay images are not scaled to each other.

**Figure S10.** Gramicidin effects on CRhOMe and TMCRhZero lifetimes.

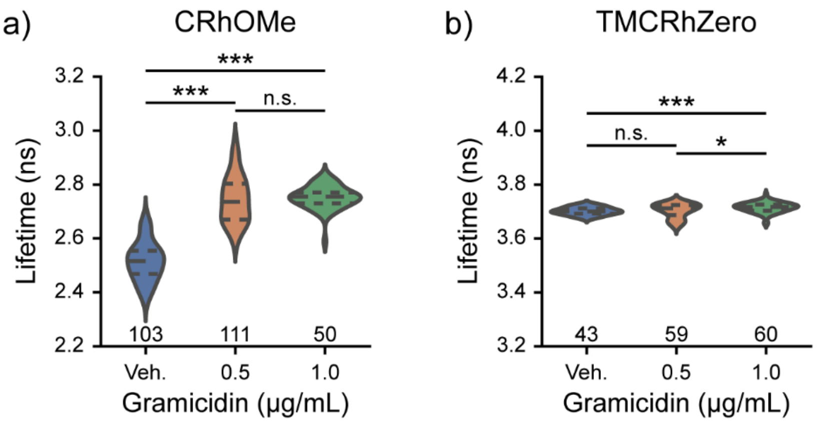

a) Violin plots of the fluorescence lifetime of CRhOMe in HEK293T incubated with varying concentrations of gramicidin or DMSO vehicle. Dashed lines within plots indicate the first quartile, median, and third quartile of the data. Significant differences between groups were observed (Welch's one-way ANOVA, $F(2, 166.3) = 352.86$, $p<0.001$). Data for vehicle and 500 ng/mL are reproduced from **Figure 7**. b) Fluorescence lifetime of TMCRhZero in HEK293T plotted as in (a). Significant differences between groups were observed (Welch's one-way ANOVA, $F(2,104.5) = 8.70$, $p<0.001$). Asterisks indicate the results of Games-Howell post hoc tests (n.s. $p>0.05$, * $p<0.05$, ** $p<0.01$, *** $p<0.001$). Sample sizes (number of cell groups) are indicated on the plot for each category. Veh. = 0.05-0.1% DMSO.

**Figure S11.** Individual recordings of the A431 $V_{mem}$ response to EGF treatment.

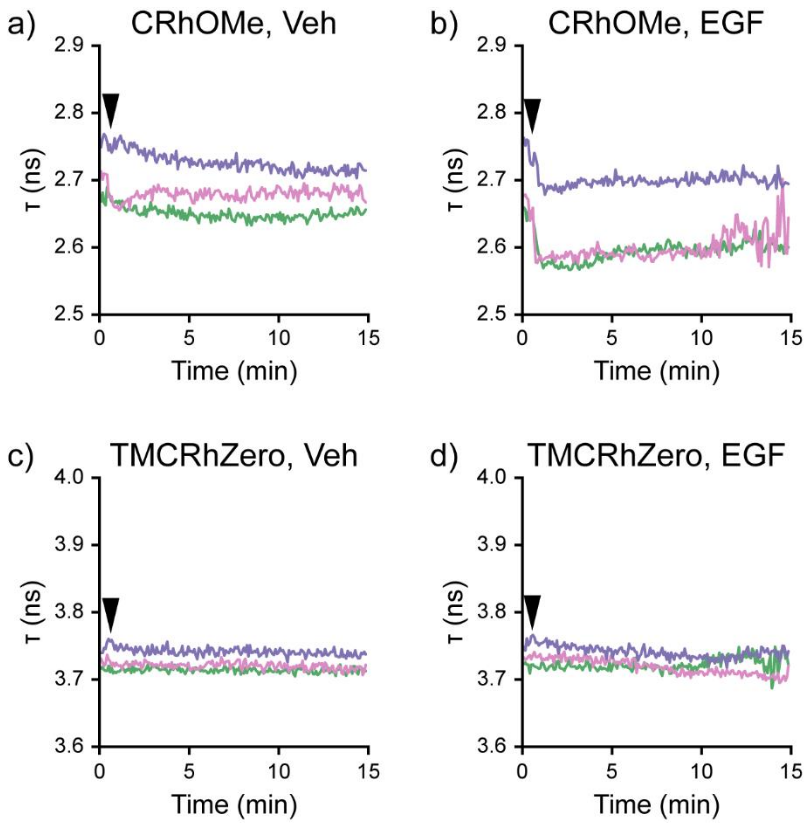

Representative individual trials for CRhOMe stained A431 treated with a) vehicle (Veh) or b) 500 ng/mL EGF. Representative individual trials for TMCRhZero stained A431 treated with c) vehicle (Veh) or d) 500 ng/mL EGF. Vehicle or EGF was added at the black arrow. Recordings are quantified as the average lifetime per frame for all contiguous cells in the field of view. Colors represent distinct recordings; there is no relationship for colors between panels.

**Figure S12.** CRhOMe staining in iCMs is membrane-localized.

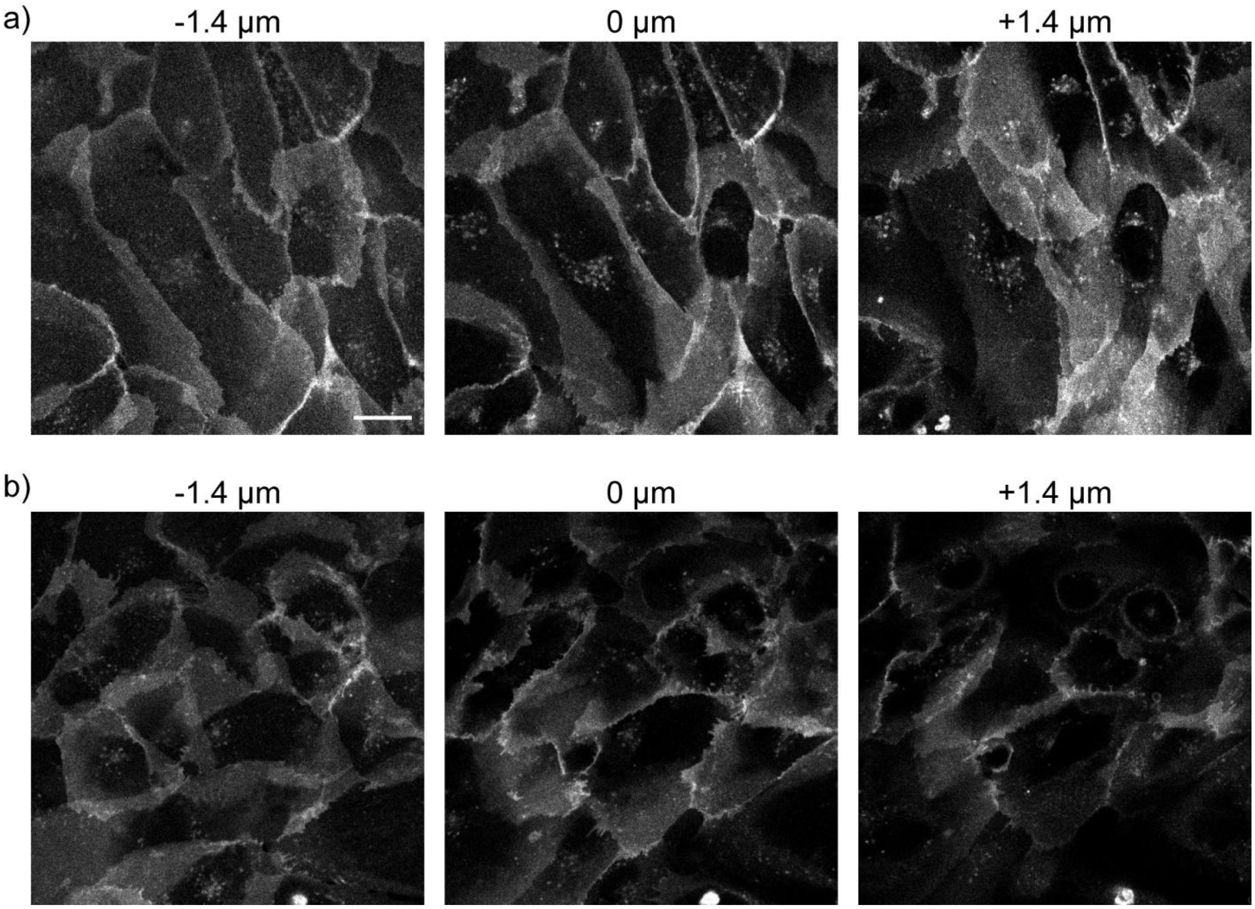

Confocal z stack images of two fields of view (a) and (b) of iCMs stained with 500 nM CRhOMe. Data were taken with the confocal pinhole set to the same settings as for FLIM (4.5 AU, 3.8 μm section). Scale bar is 20 μm.

**Figure S13.** Comparison between lifetime, photon count, and relative photon count in hiPSC-CMs.

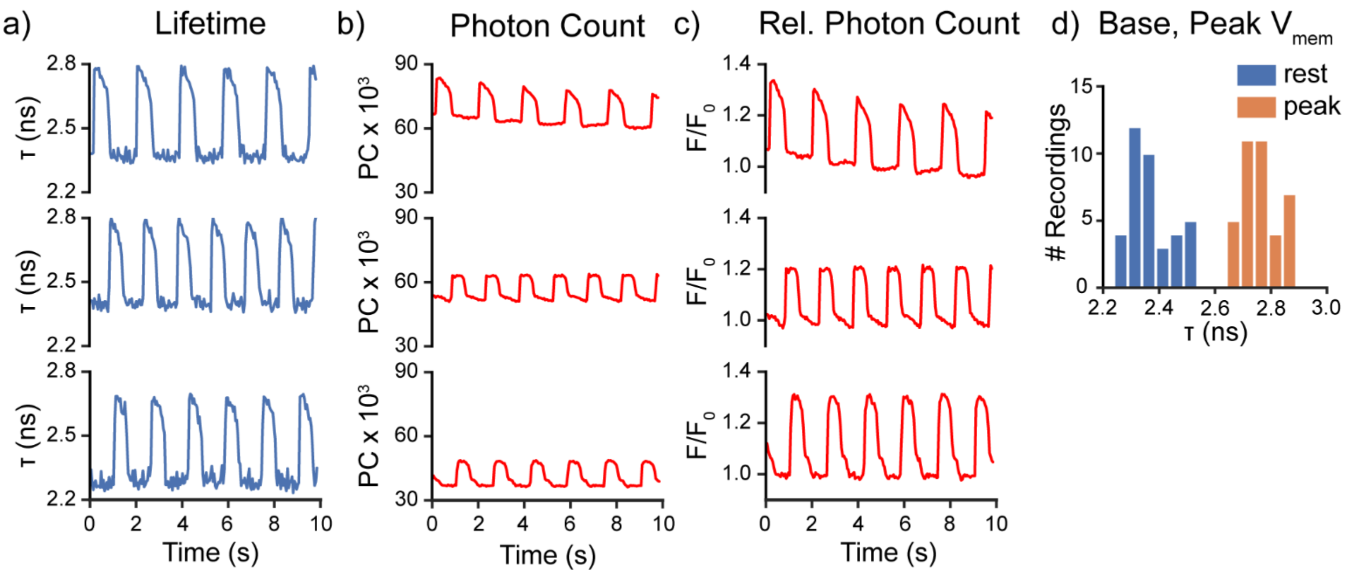

Data are from **Figure 5** in the main text. Panels (a) and (c) are reproduced here for comparison purposes with the raw photon count rendering in panel b.

a) Fluorescence lifetime recordings from 3 representative fields of view. Data are not filtered or processed in any way after fitting. b) Photon count recordings from the same fields of view as in (a). c) Relative photon count (photon count divided by the average photon count baseline) for the same fields of view as in (a). d) Histograms representing the distributions of lifetimes observed at the base of an action potential (rest) and at the peak.

**Figure S14.** The fluorescence lifetime of TMCRhZero does not change in beating hiPSC-CMs

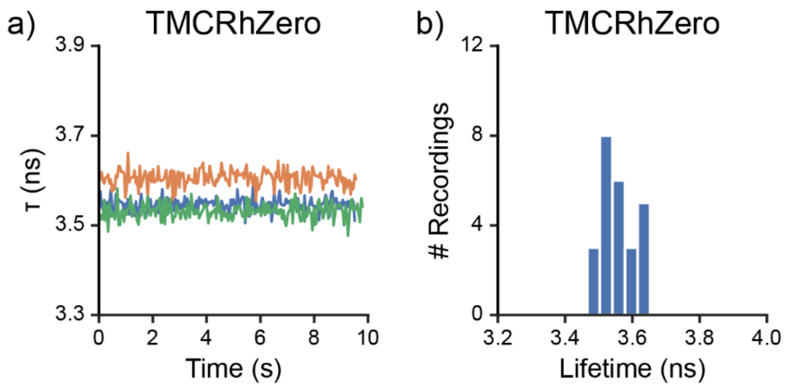

a) Fluorescence lifetime recordings of TMCRhZero in hiPSC-CMs from 3 representative fields of view. Data are not filtered or processed in any way after fitting. b) Histograms representing the distributions of lifetimes from imaging hiPSC-CMs with TMCRhZero

**Figure S15.** Effects of CRhOMe concentration on recorded iCM APs.

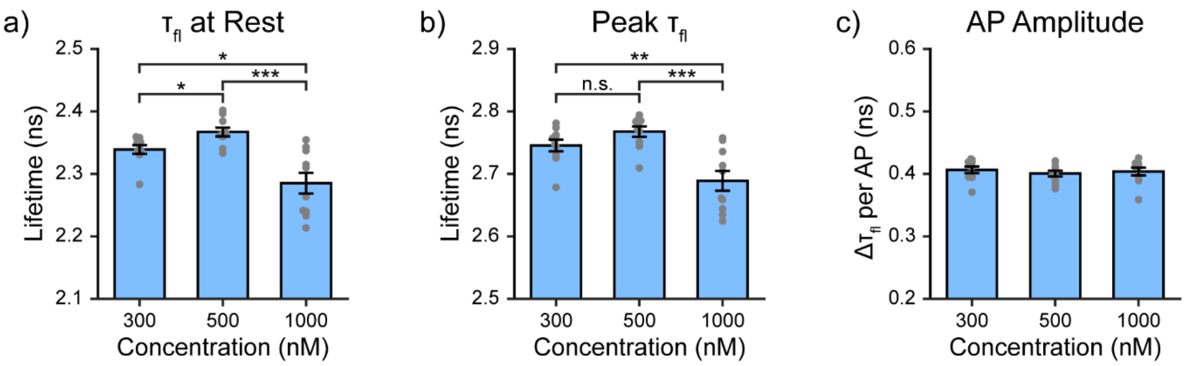

a) Resting lifetime of CRhOMe iCMs at different probe concentrations. Significant differences between concentrations were observed (Welch's ANOVA, $F(2,16.80) = 11.35$, $p=0.0008$). b) Lifetime at the peak of iCM action potential (AP) at different CRhOMe concentrations. Significant differences between conditions were observed (Welch's ANOVA, $F(2, 17.13)=9.48$, $p=0.0017$). c) Action potential height (difference between resting and peak lifetime) at 300, 500, and 1000 nM CRhOMe. No significant differences between concentrations were observed (Fisher's ANOVA, $F(2,27)=0.291$, $p=0.75$). Data are shown as mean ± SEM of 10 recordings from two wells for each concentration. Asterisks reflect the significance level of Games-Howell post hoc tests (n.s. $p>0.05$, *$p<0.05$, **$p<0.01$, *** $p<0.001$). The 500 nM data shown here is a subset of the data shown in **Figure 5** in the main text and is reproduced here for comparison purposes.

### Supporting Spectra

**Spectrum S1.** $^{1}$H NMR of compound 2

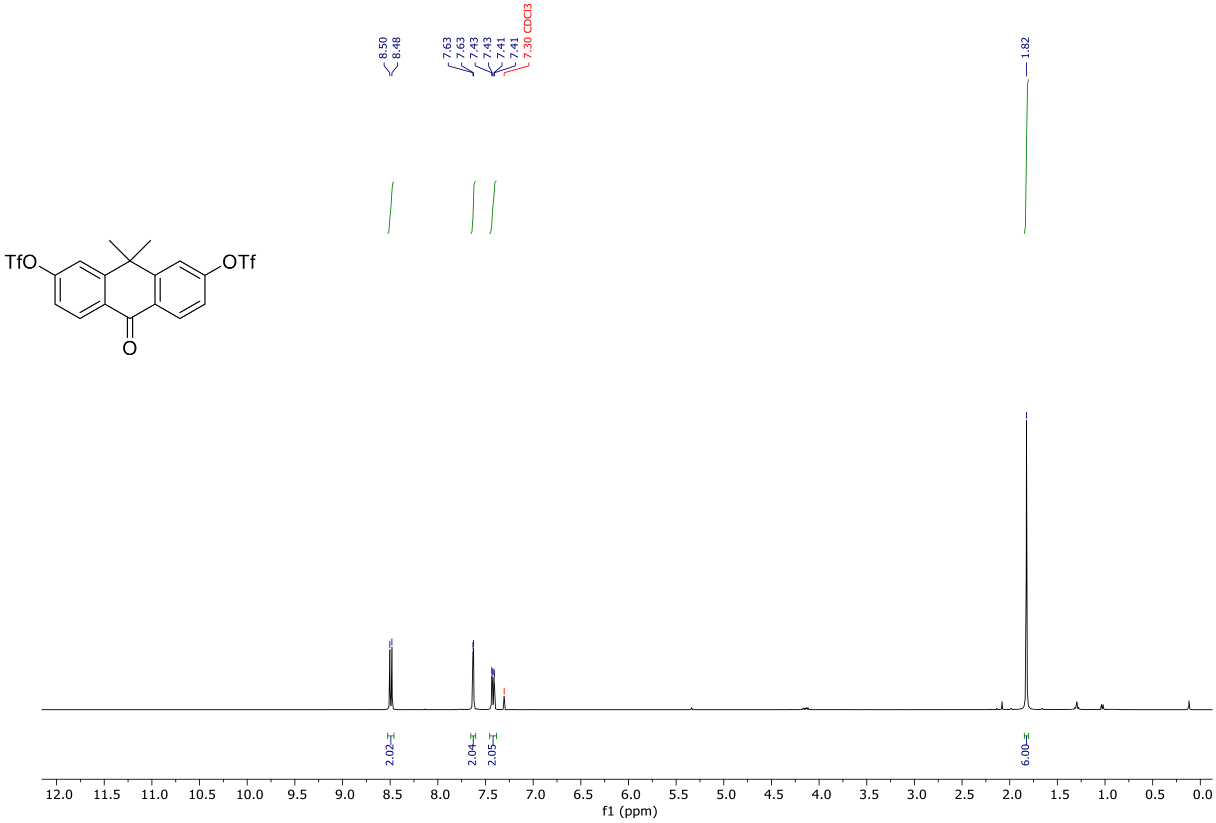

**Spectrum S2.** $^{1}$H NMR of compound 3

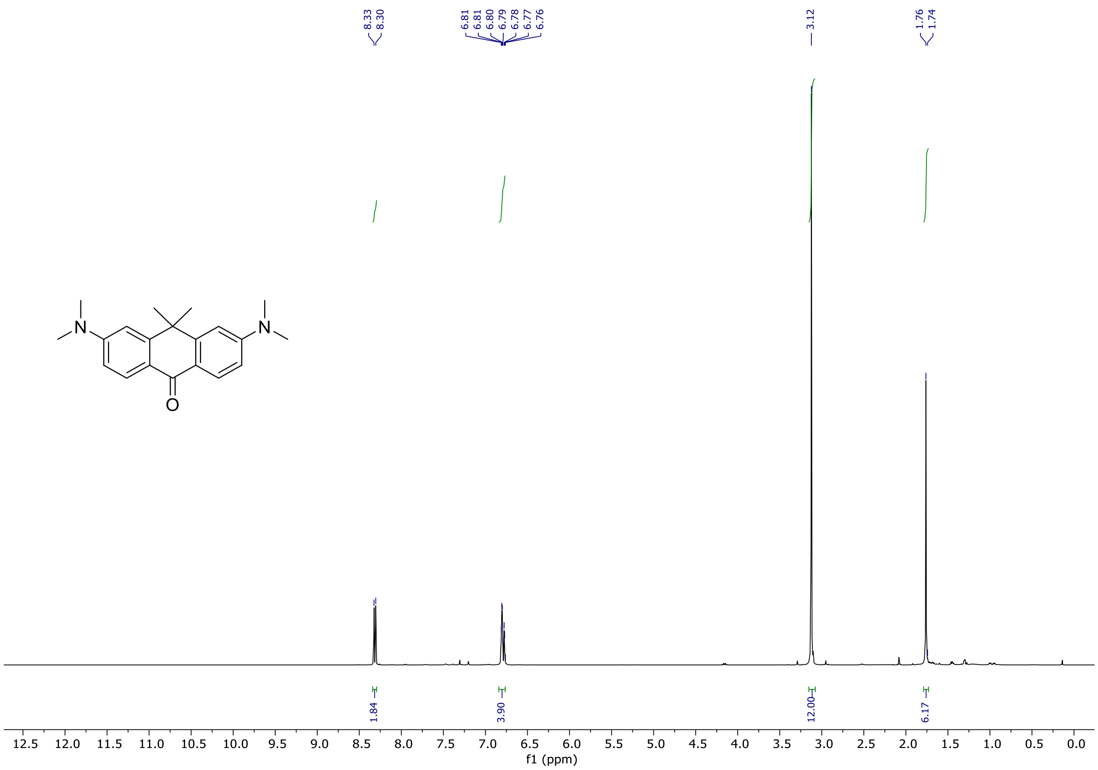

**Spectrum S3.** $^{13}$C NMR of compound 3

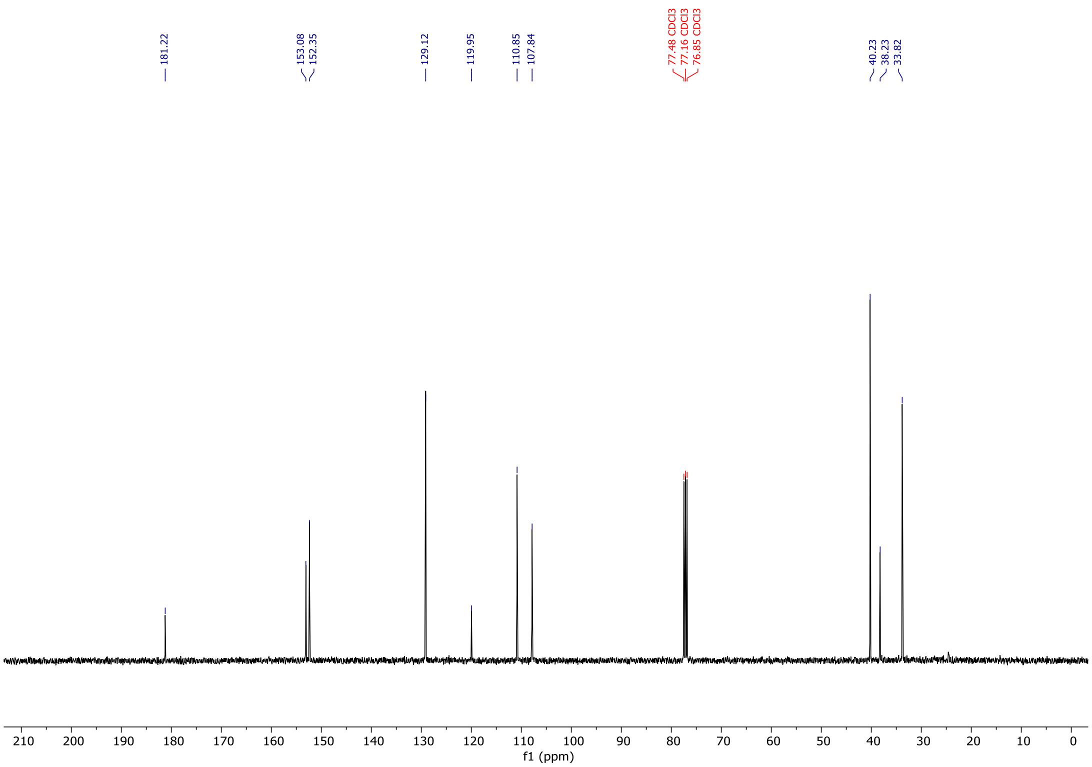

**Spectrum S4.** $^{1}$H NMR of compound 6

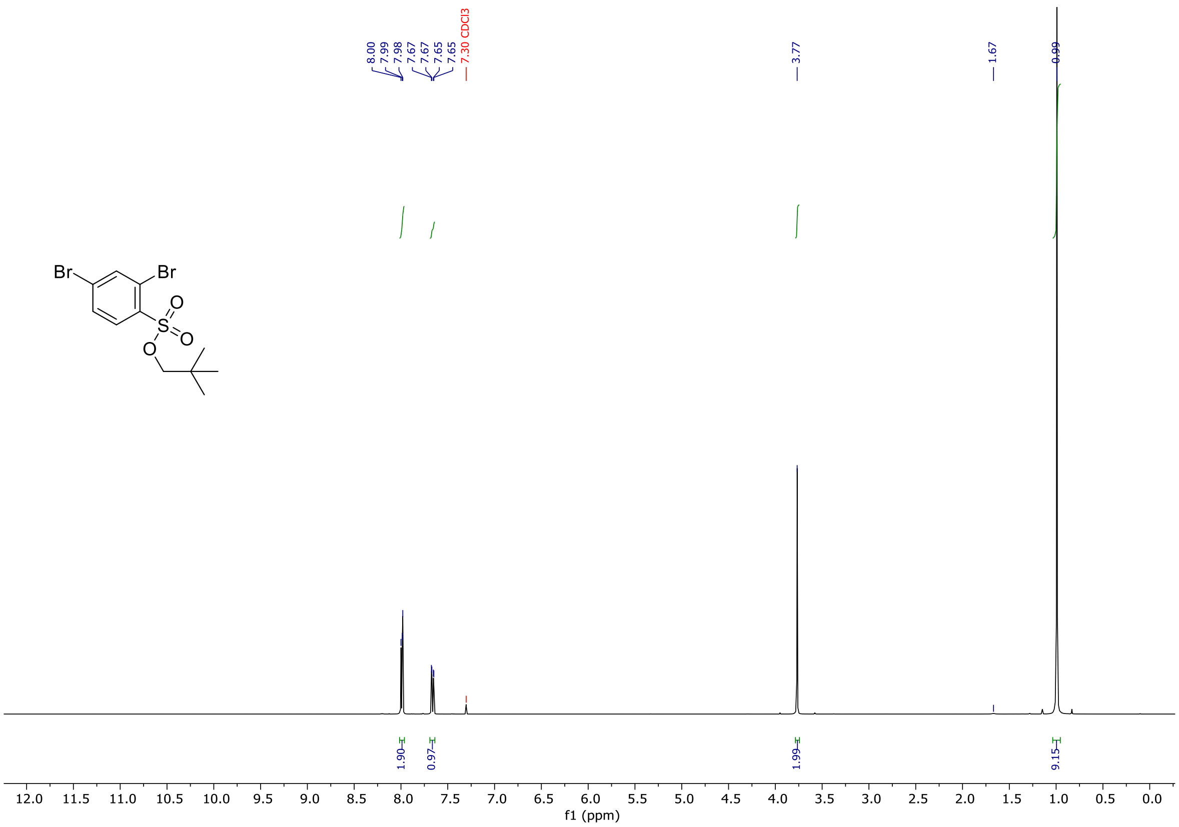

**Spectrum S5.** $^{13}$C NMR of compound 6

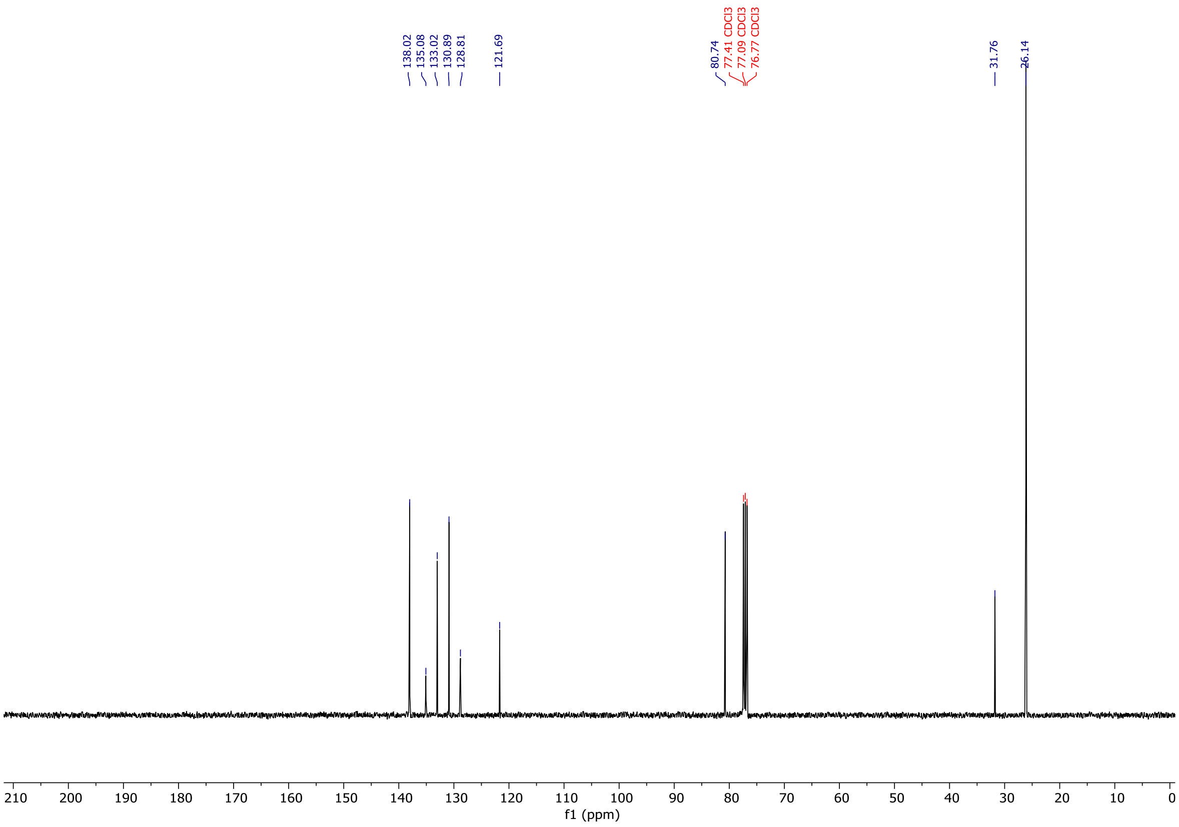

**Spectrum S6.** $^{1}$H NMR of compound 7

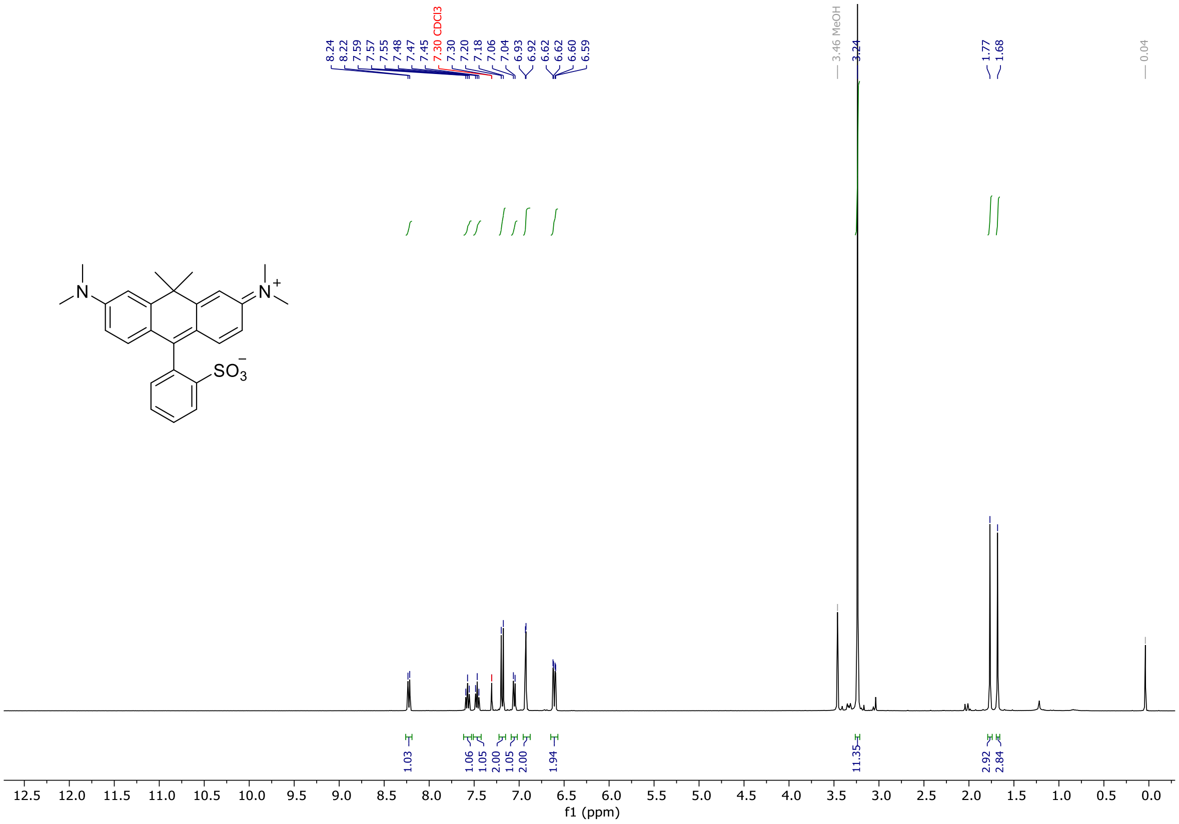

**Spectrum S7.** $^{13}$C NMR of compound 7

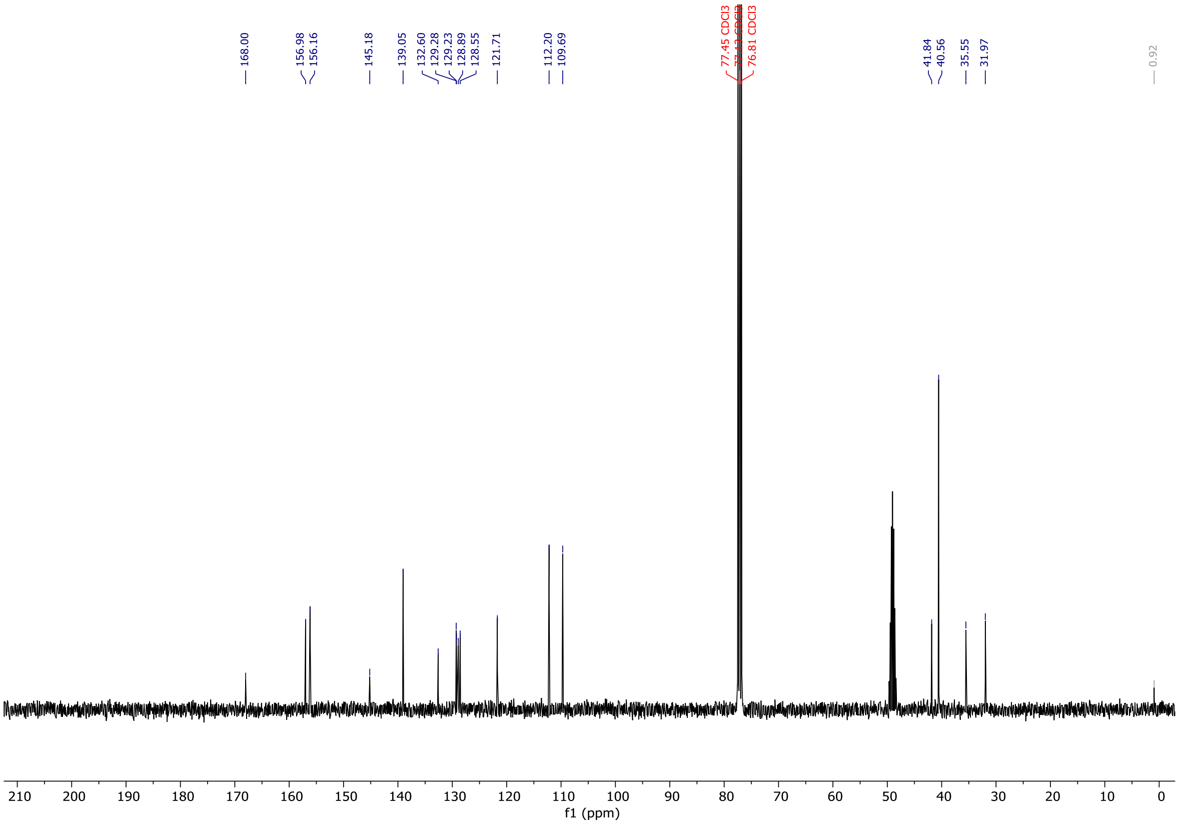

**Spectrum S8.** LCMS of Compound 7

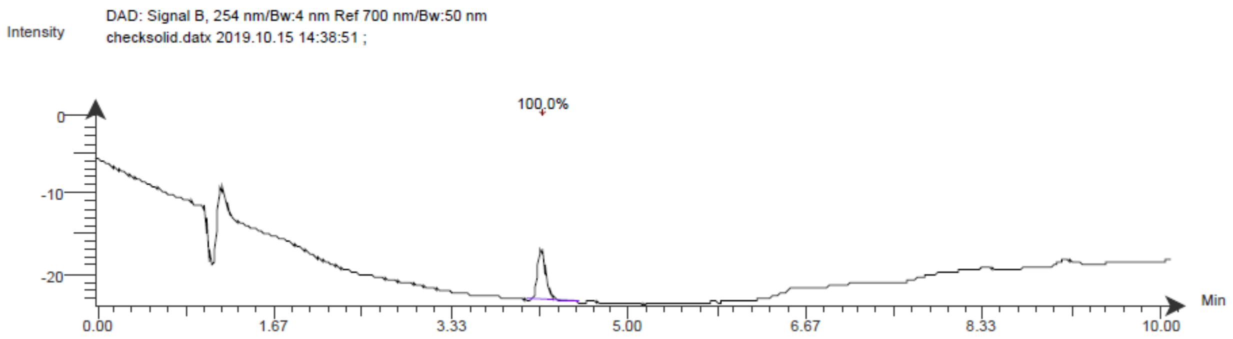

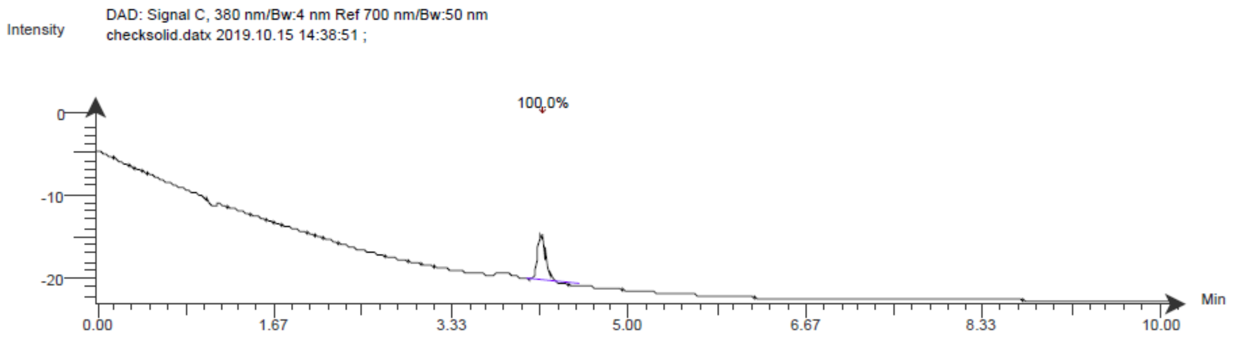

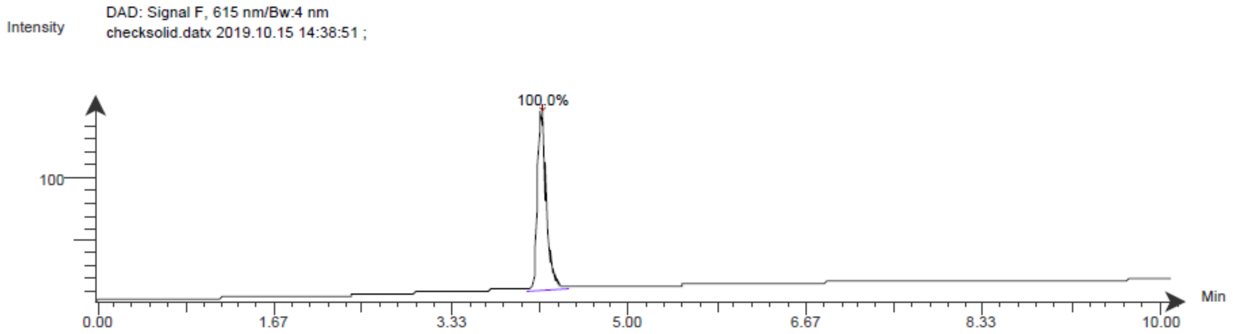

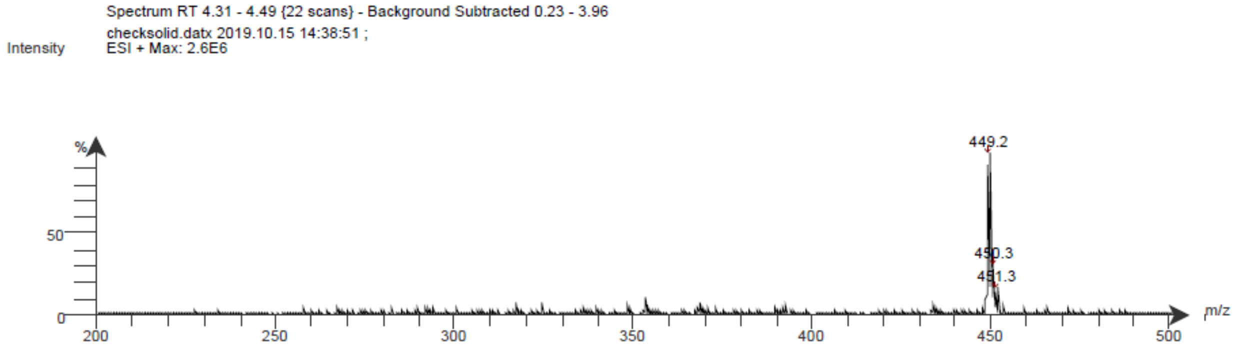

**Spectrum S9.** $^{1}$H NMR of compound 8

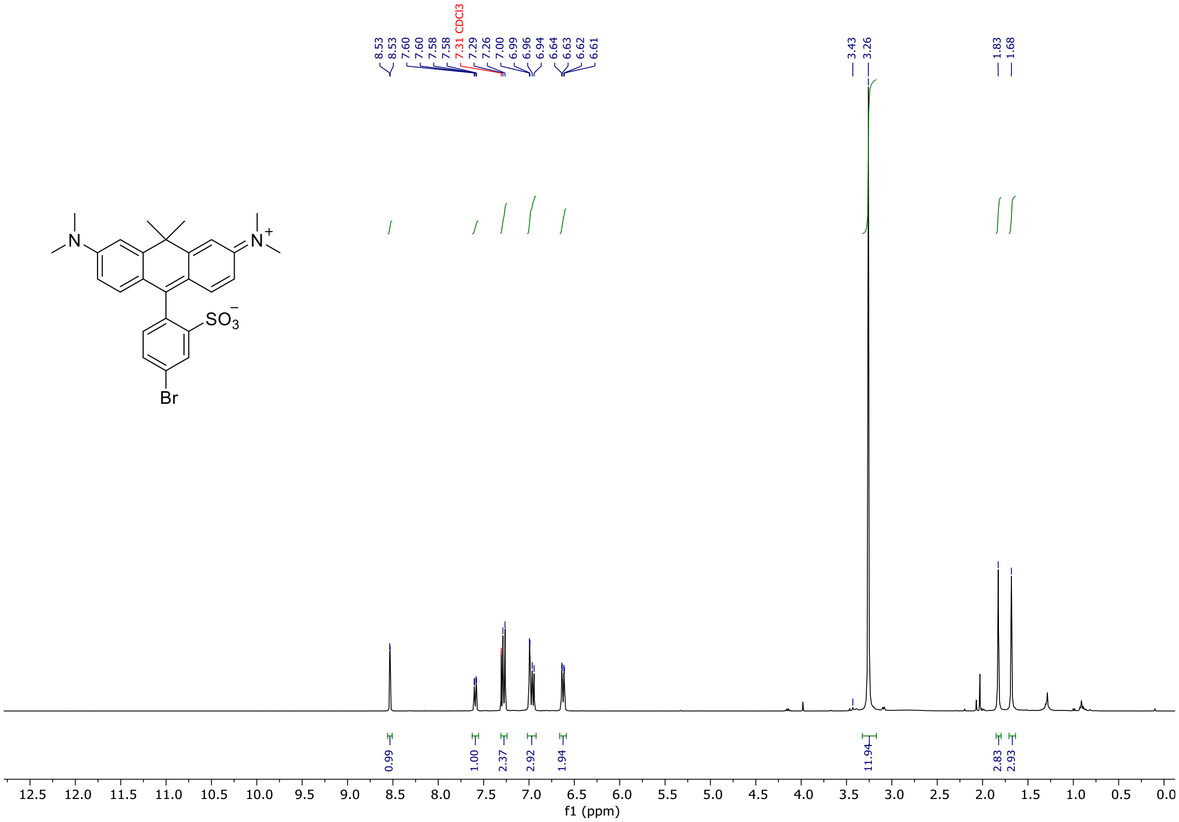

**Spectrum S10.** $^{13}$C NMR of compound 8

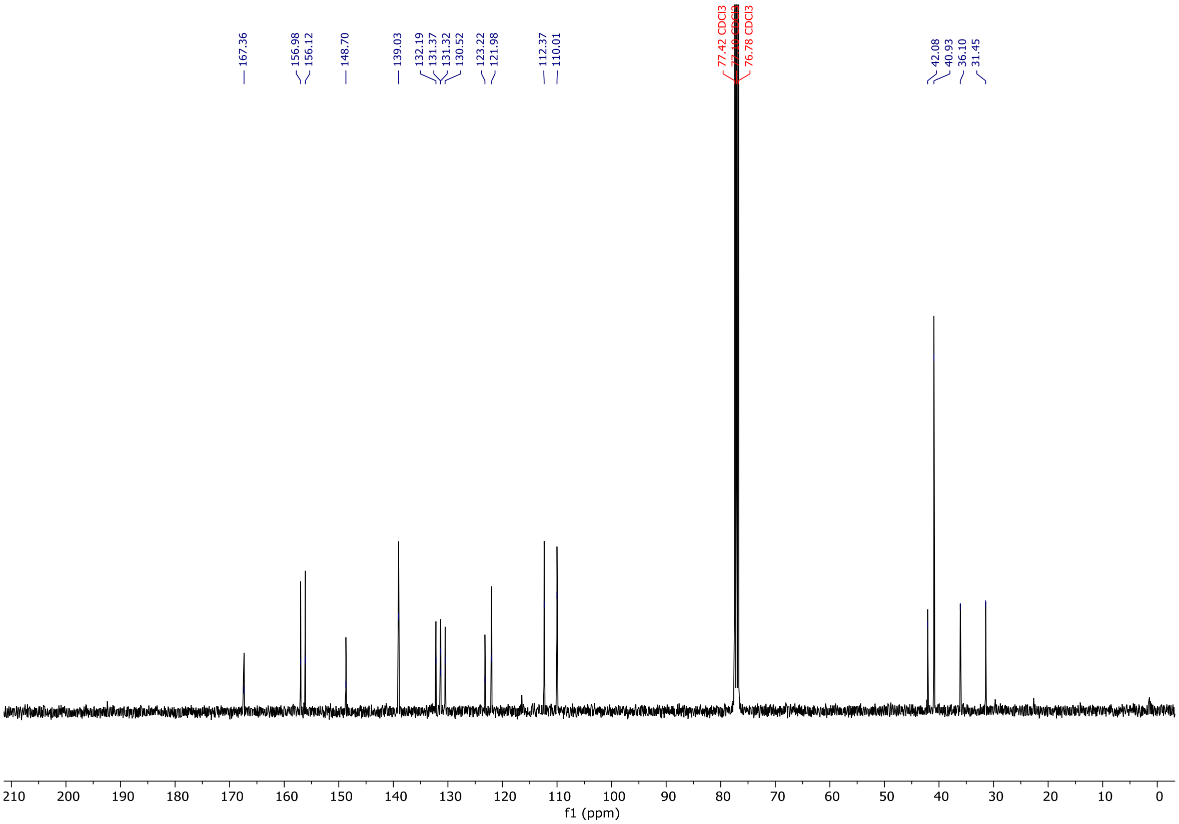

**Spectrum S11.** $^{1}$H NMR of compound 9

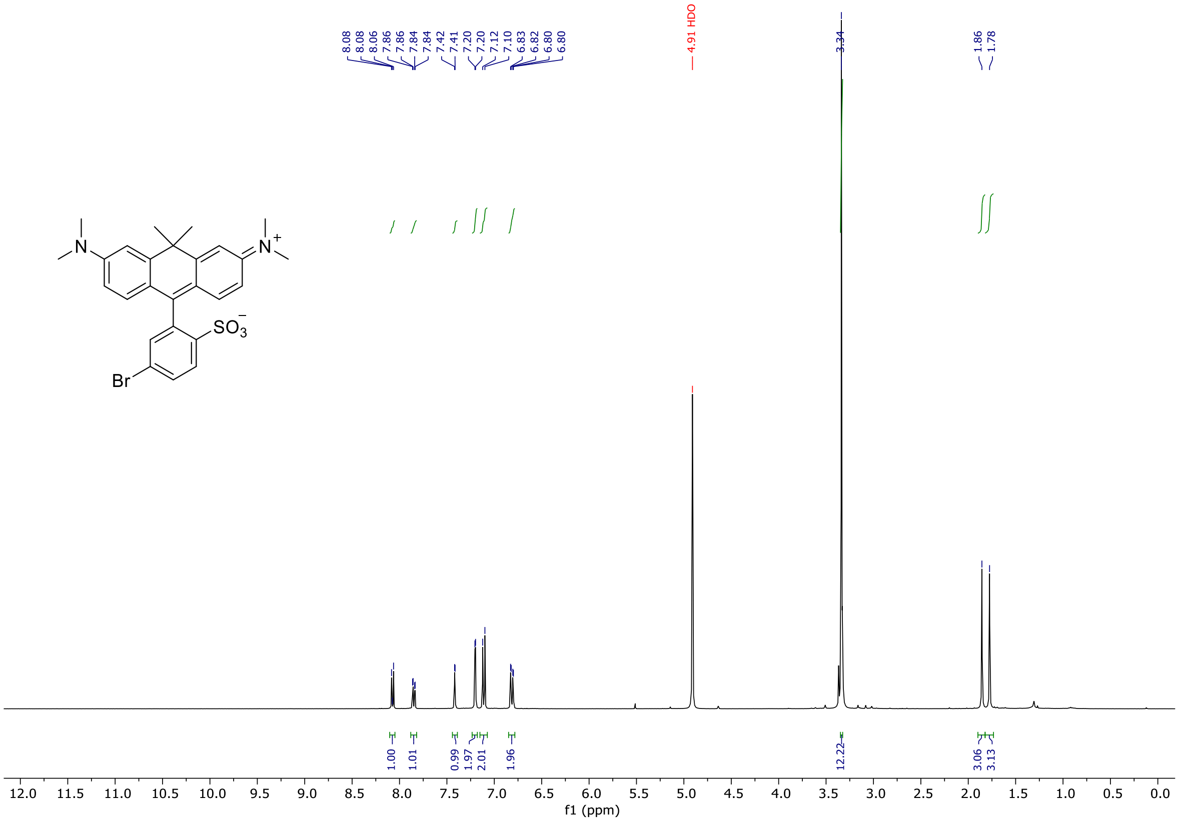

**Spectrum S12.** $^{13}$C NMR of compound 9

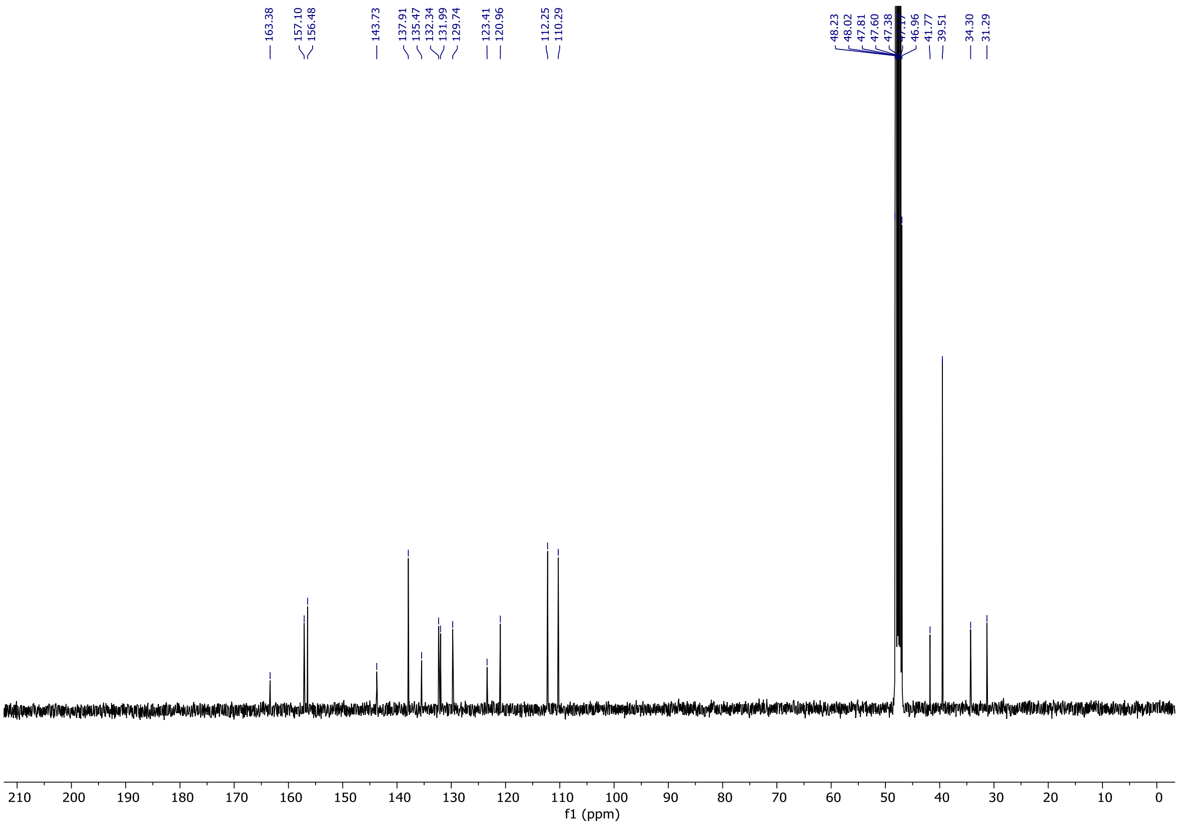

**Spectrum S13.** $^{1}$H NMR of compound 13

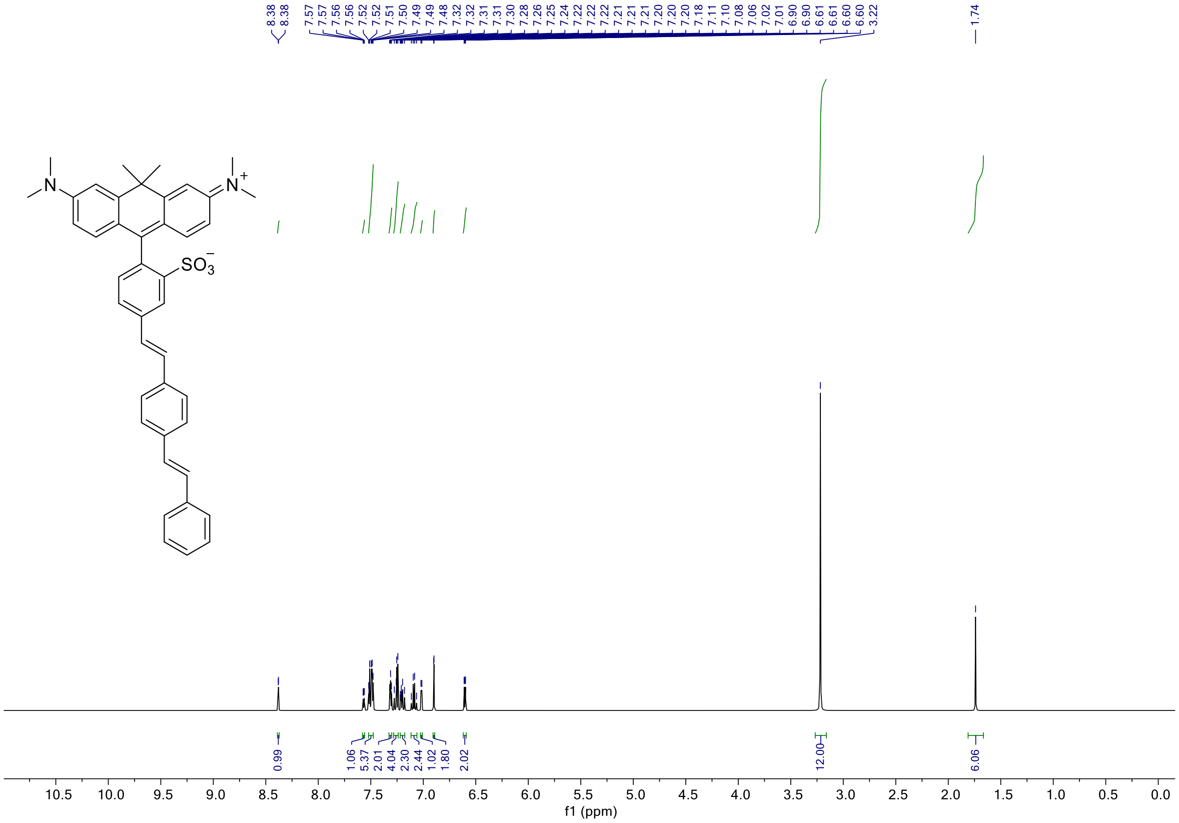

**Spectrum S14.** $^{13}$C NMR of compound 13

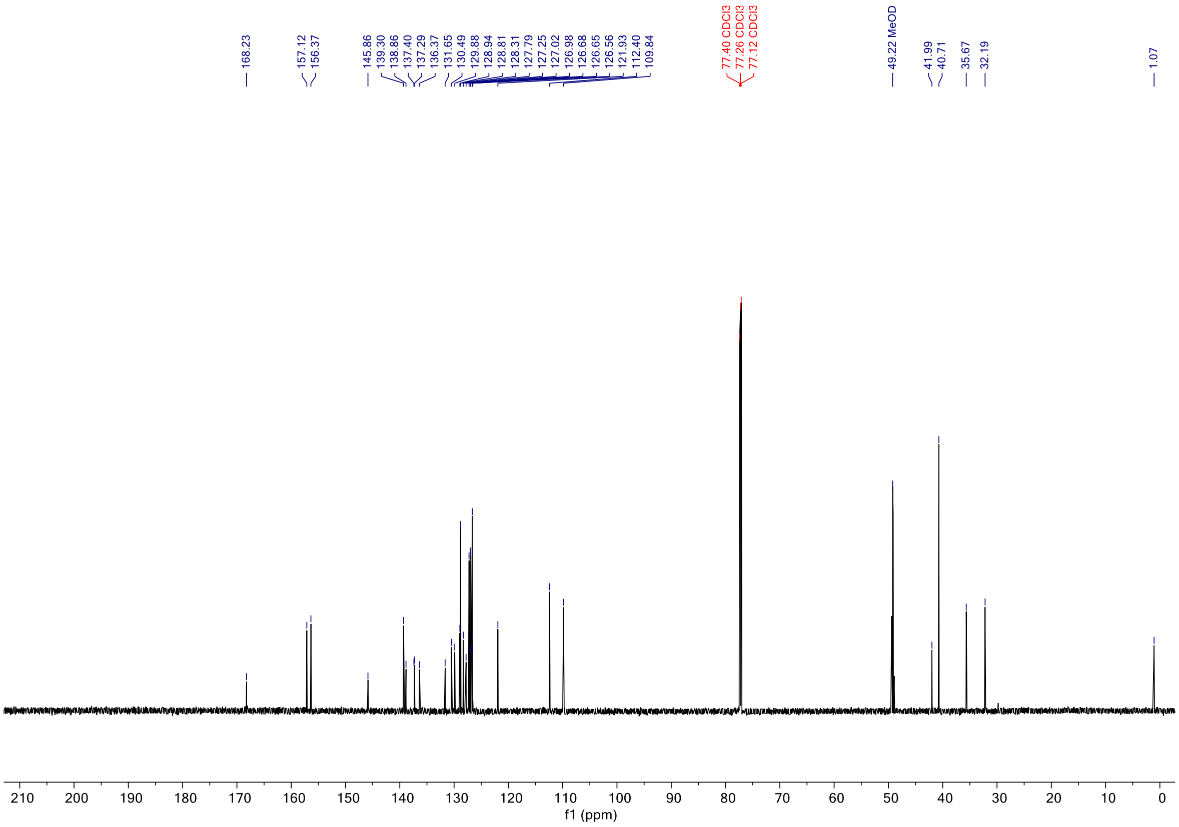

**Spectrum S15.** LCMS of compound 13

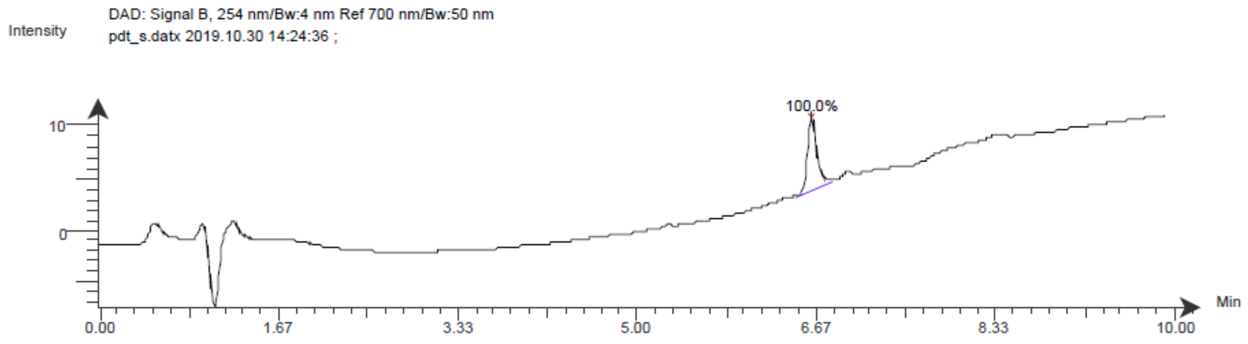

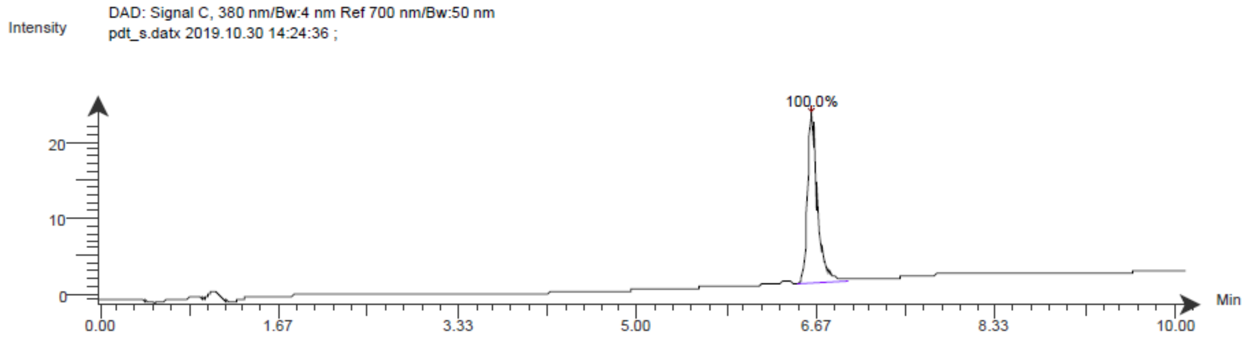

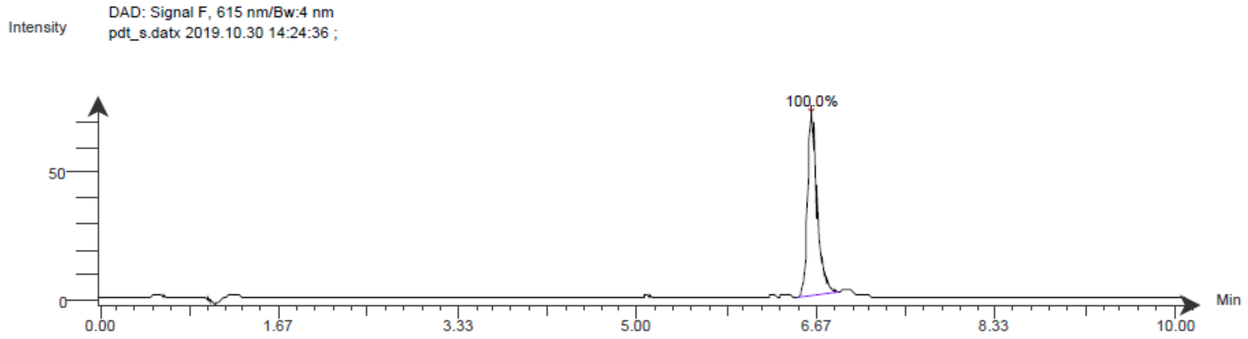

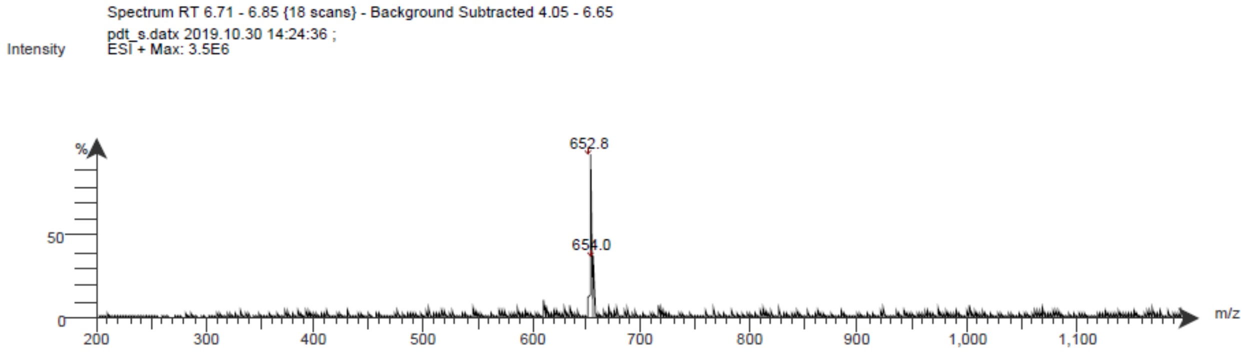

**Spectrum S16.** $^{1}H$ NMR of compound 14

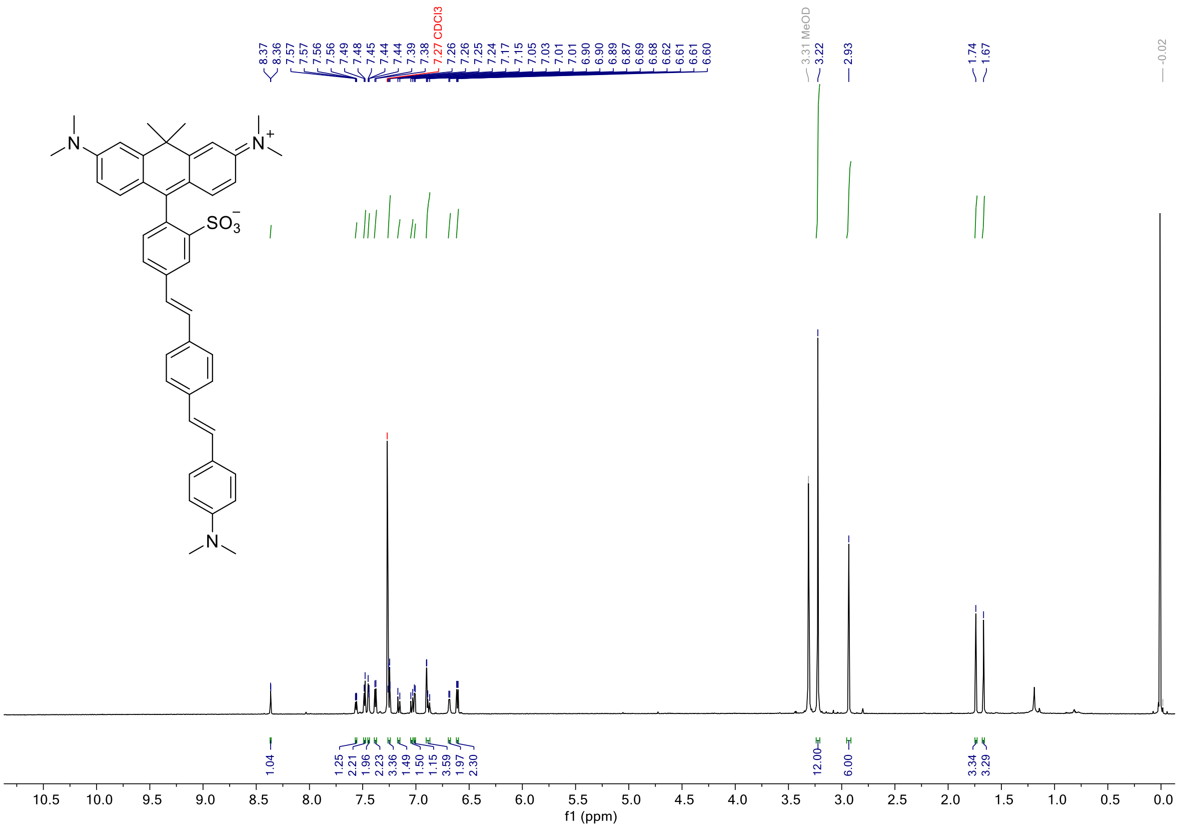

**Spectrum S17.** $^{13}C$ NMR of compound 14

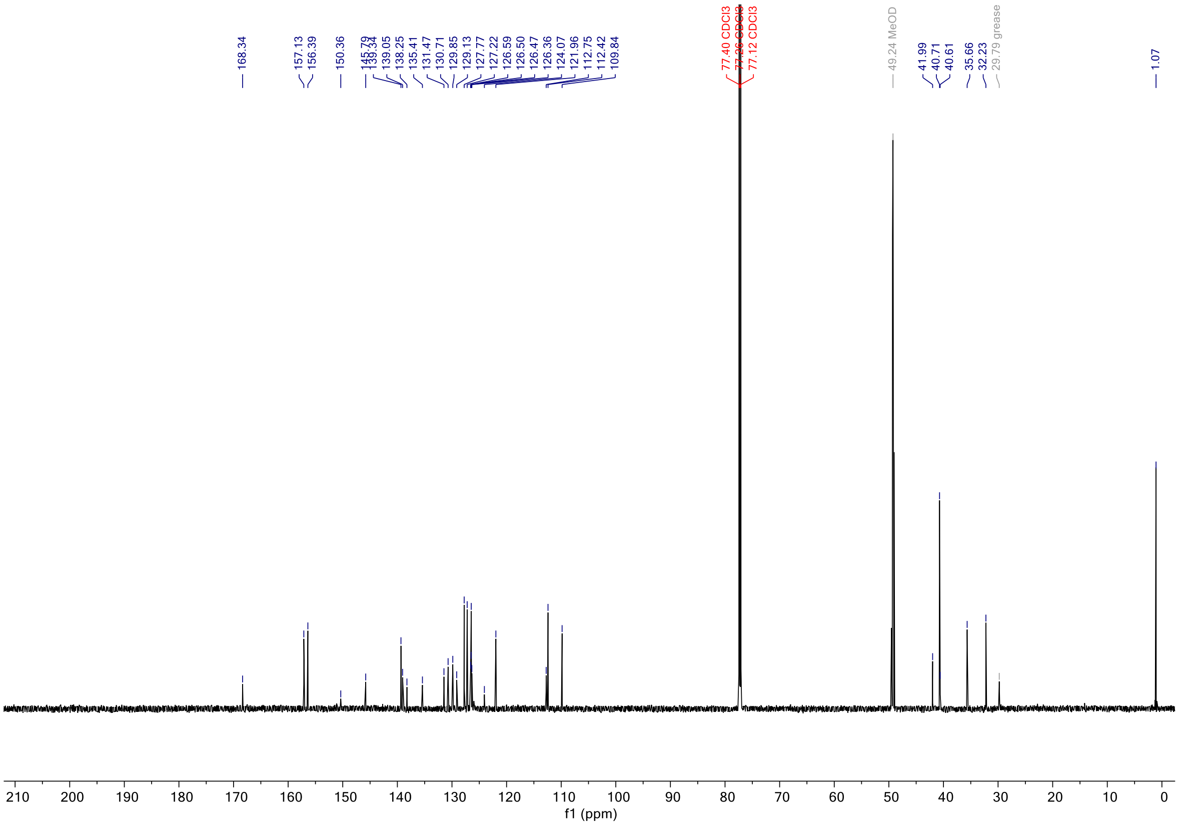

**Spectrum S18.** LCMS of compound 14

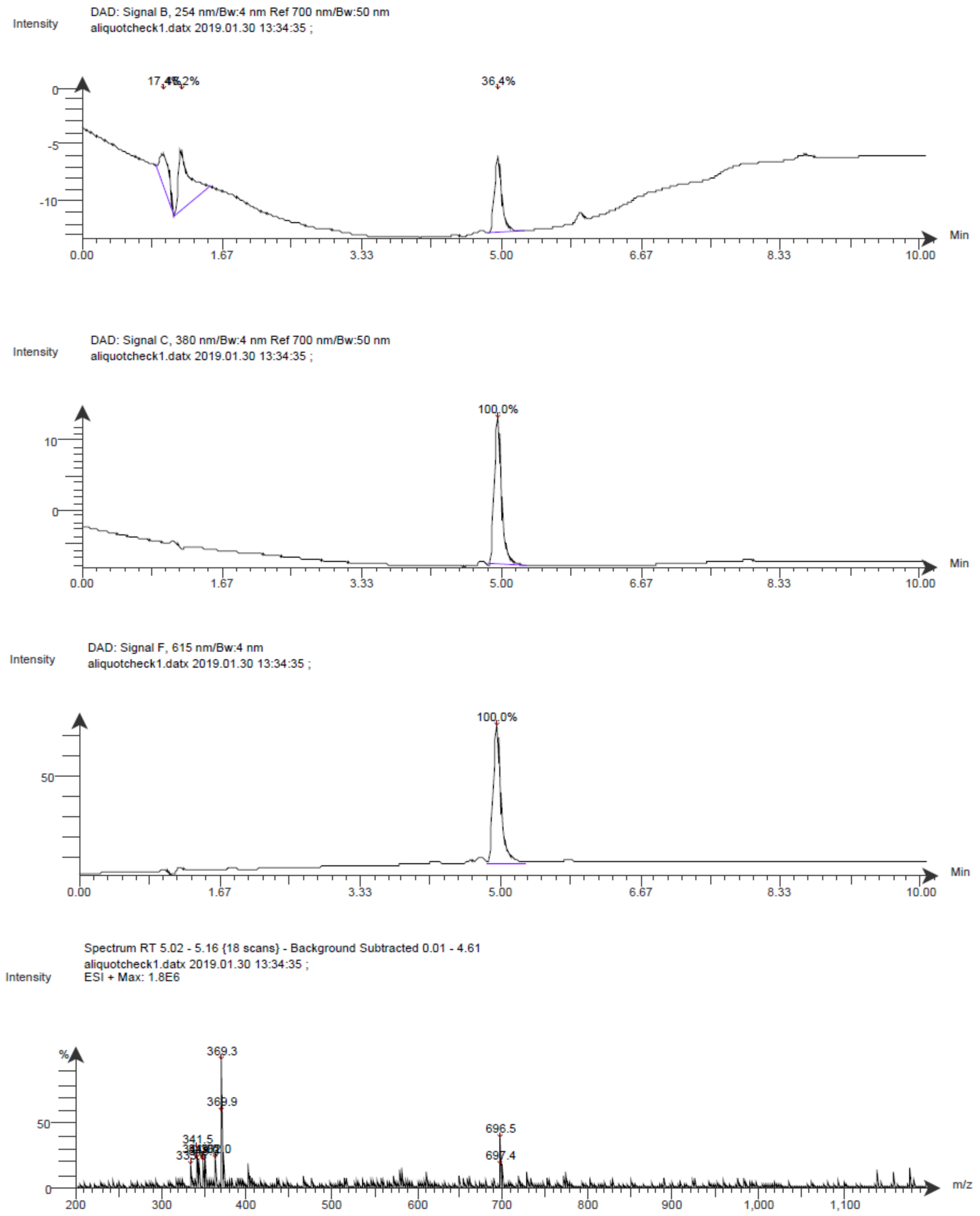

**Spectrum S19.** $^{1}$H NMR of compound 15

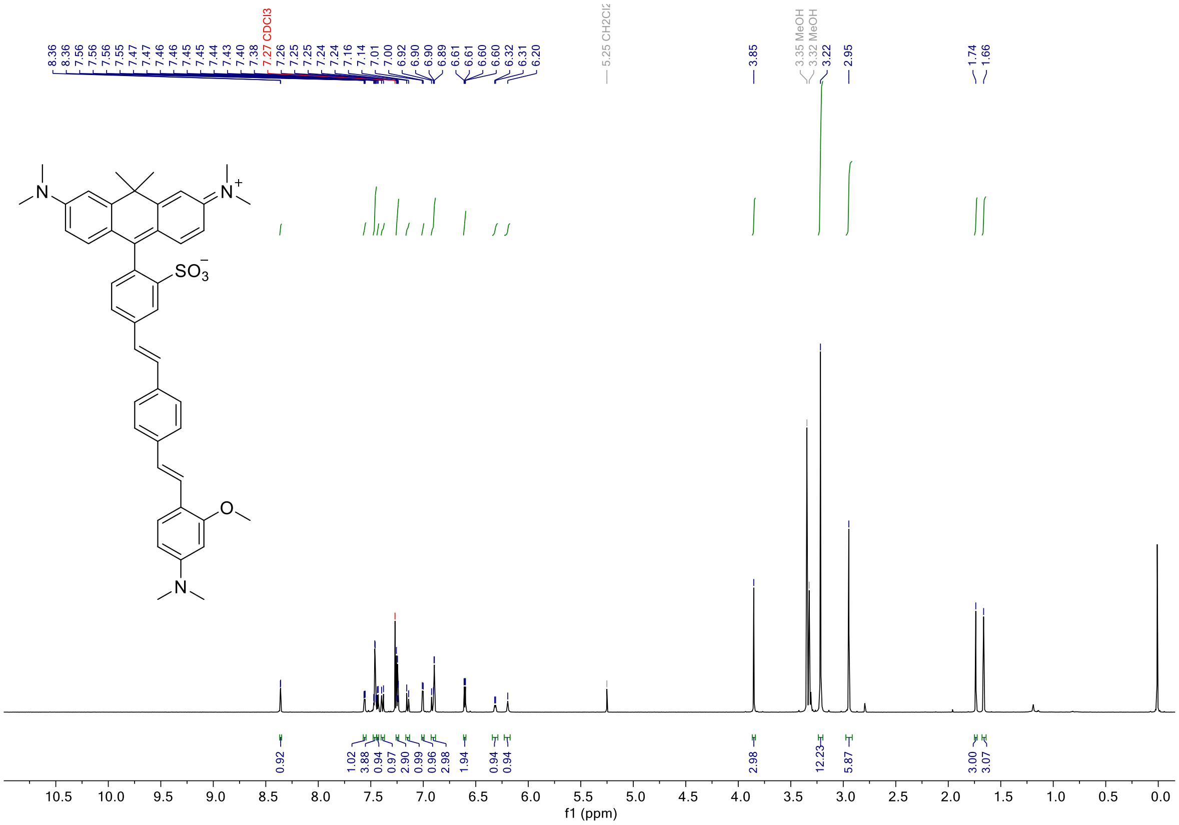

**Spectrum S20.** $^{13}$C NMR of compound 15

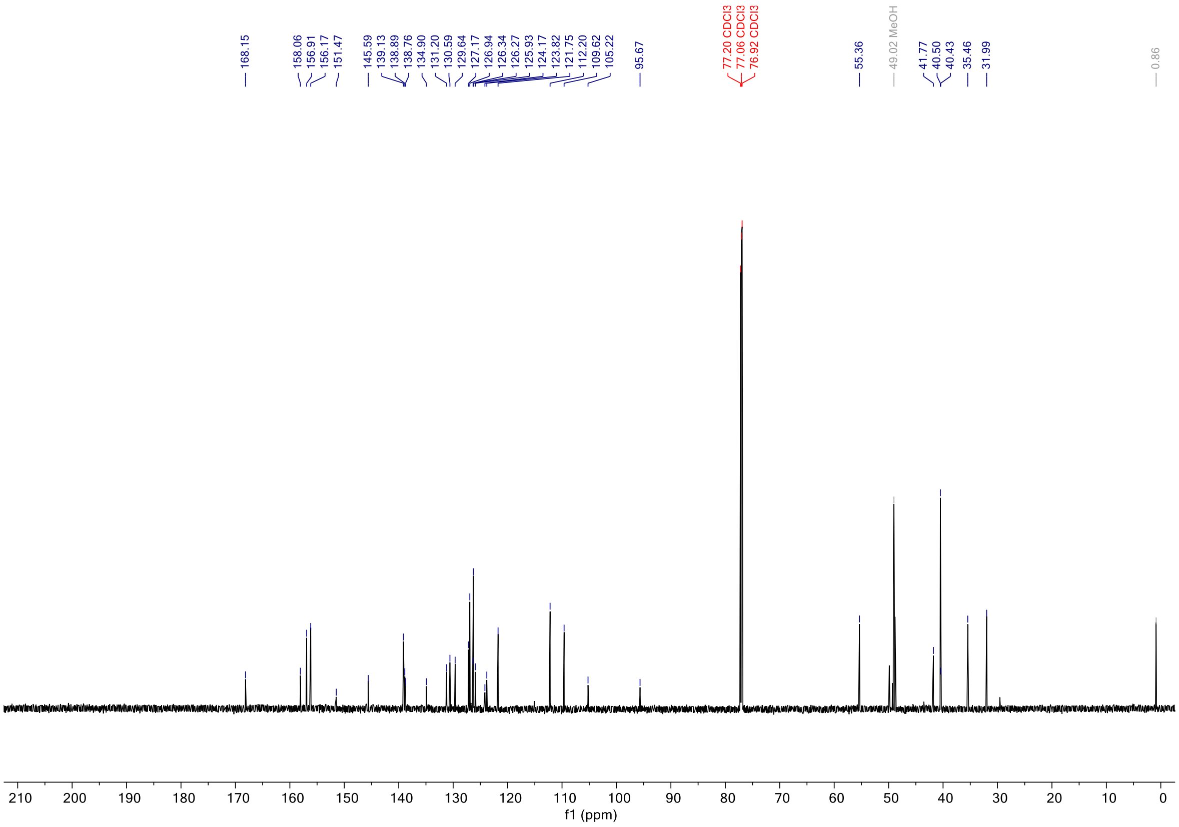

**Spectrum S21.** LCMS of compound 15

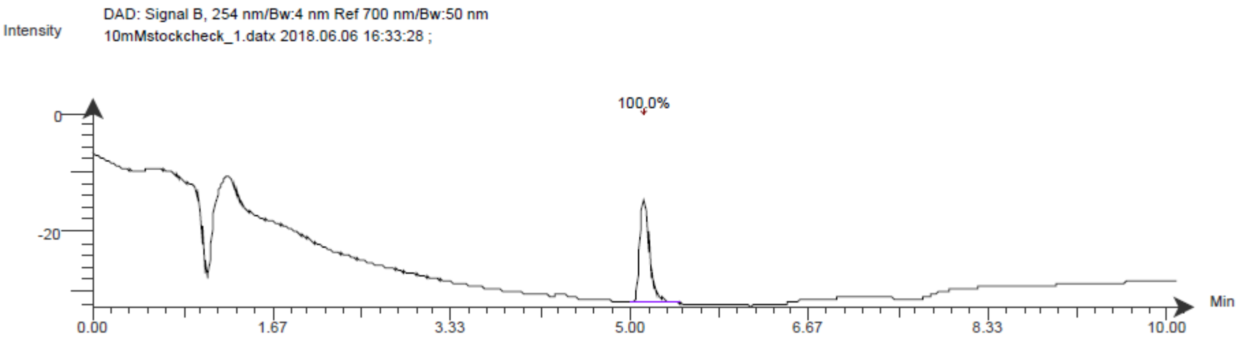

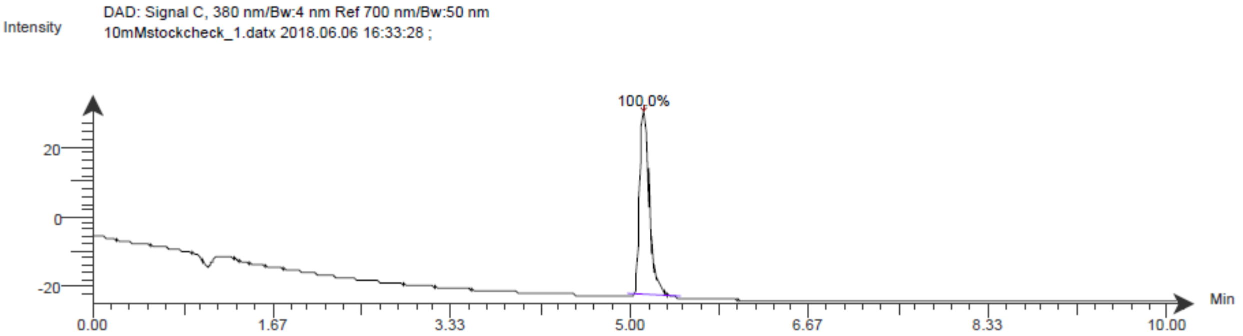

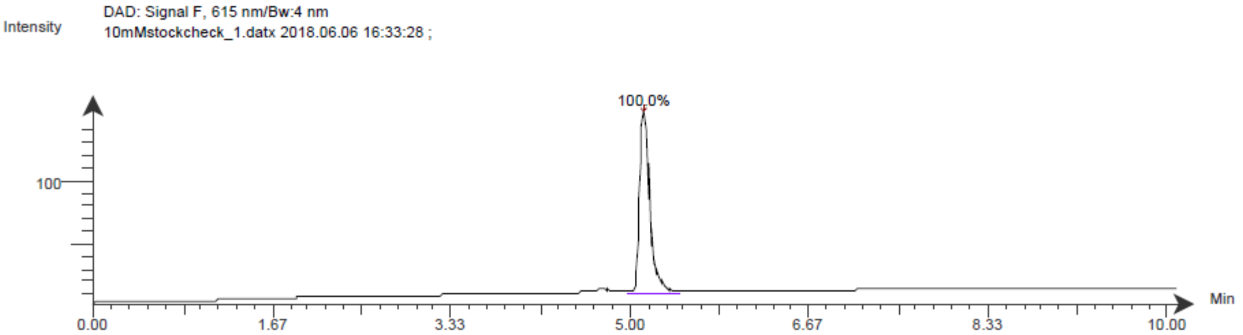

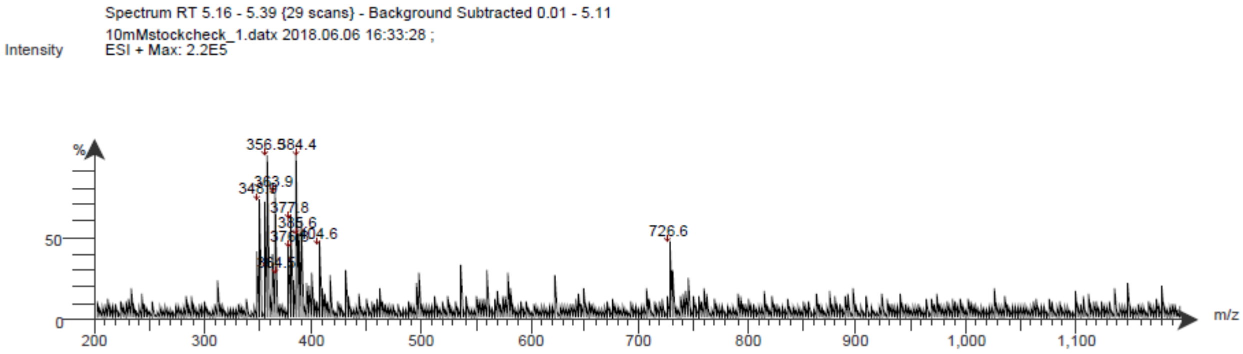

**Spectrum S22.** $^{1}$H NMR of compound 16

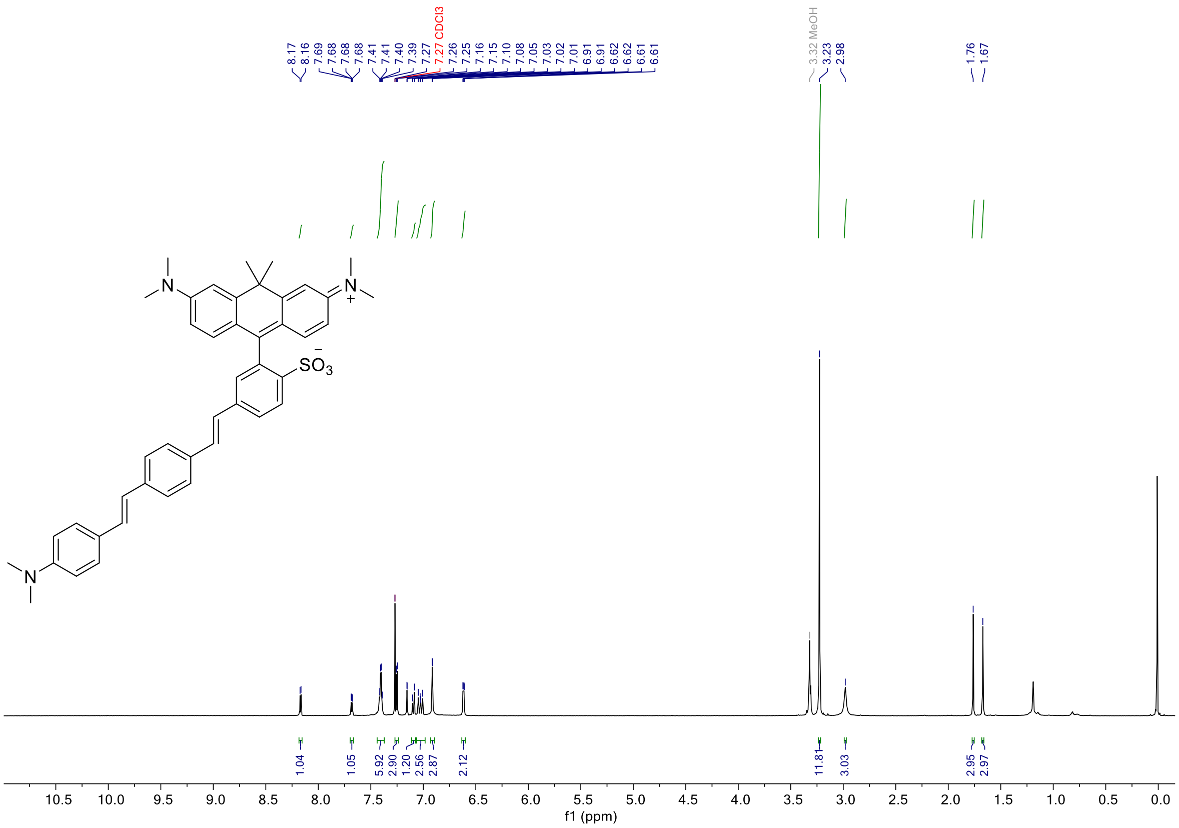

**Spectrum S23.** $^{13}$C NMR of compound 16

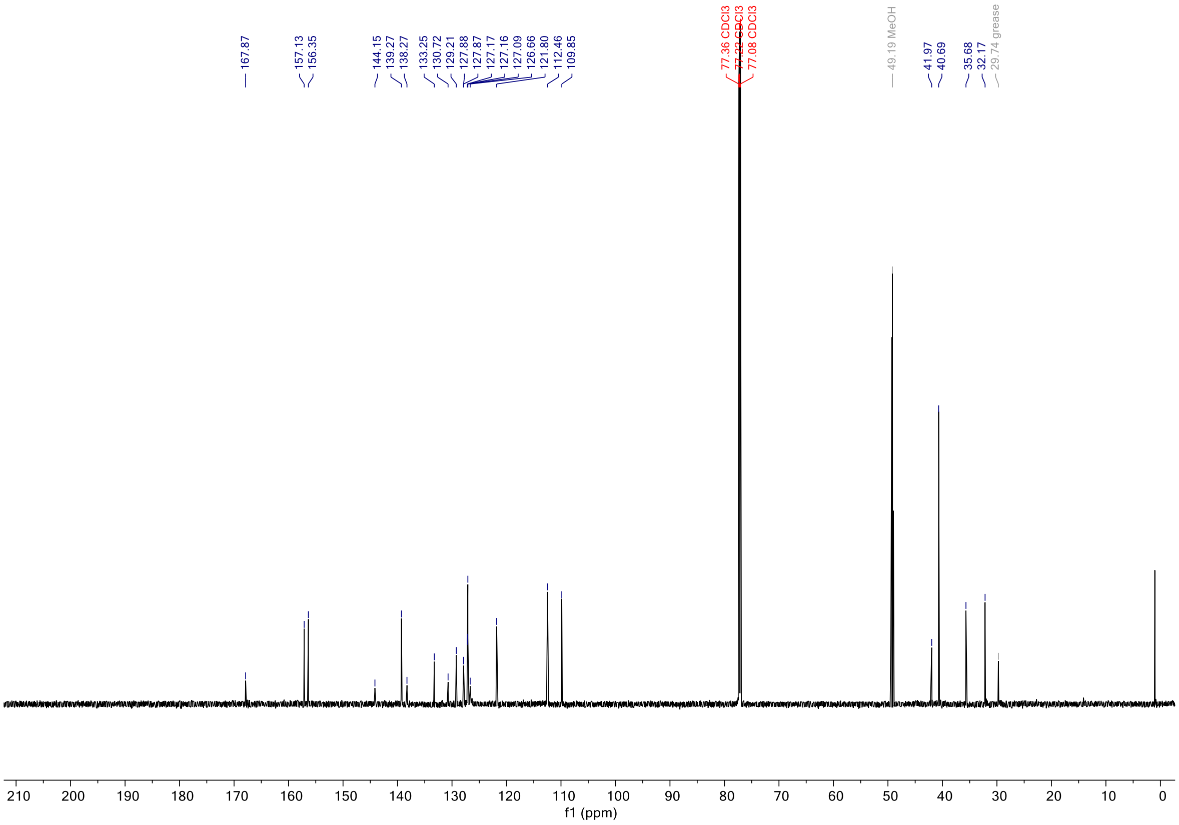

**Spectrum S24.** LCMS of compound 16

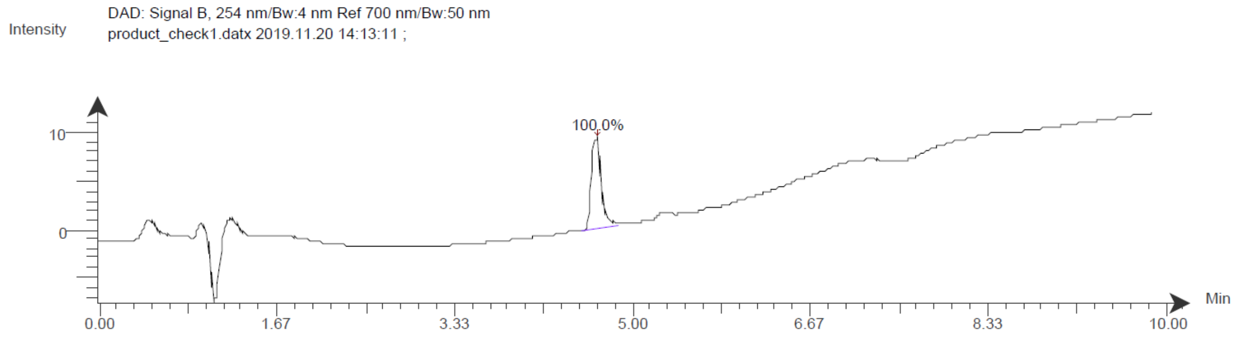

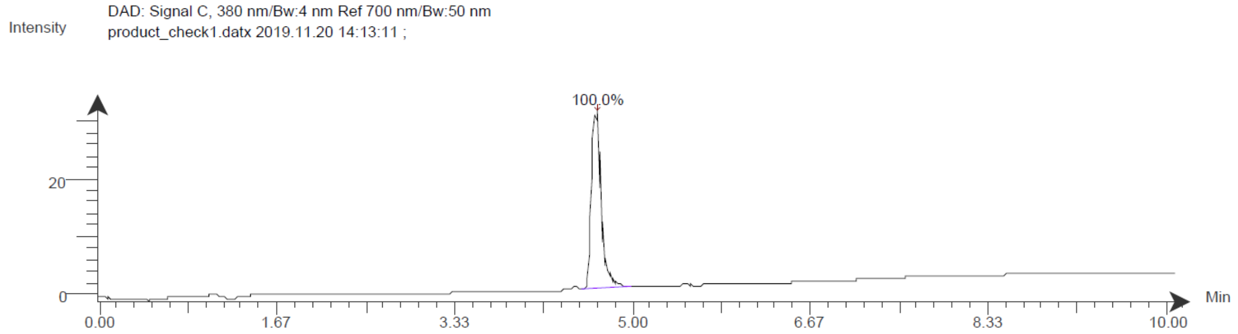

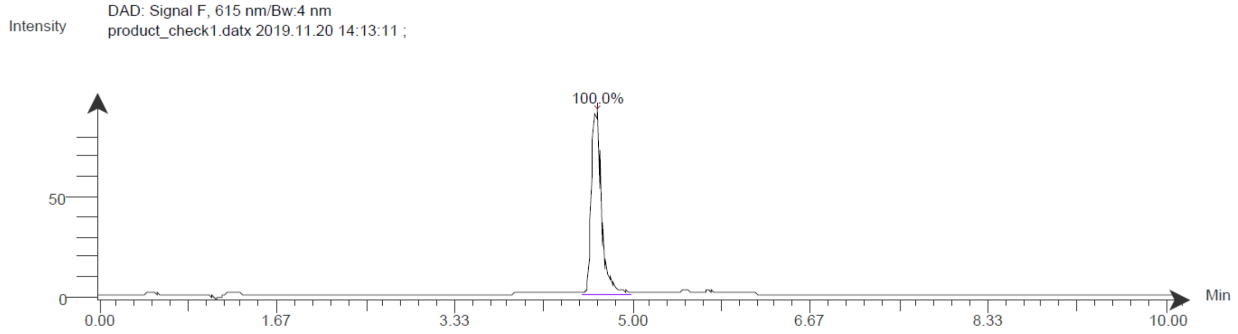

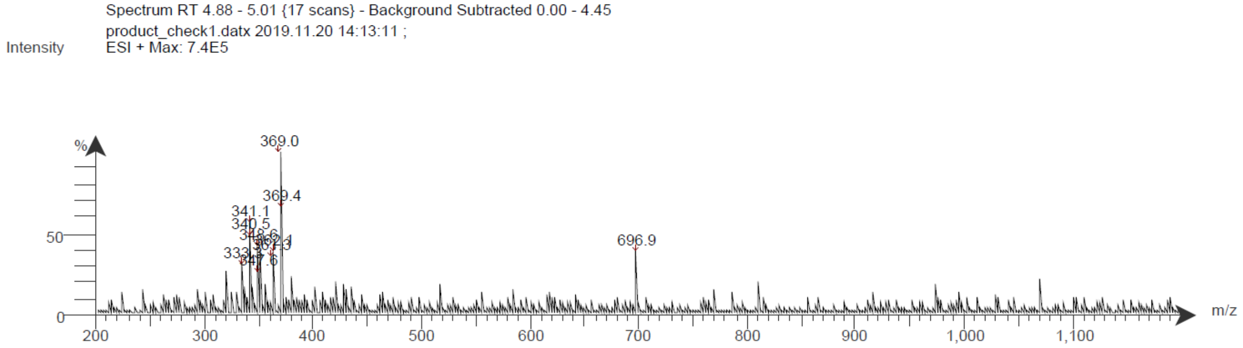

**Spectrum S25.** $^{1}$H NMR of compound 17

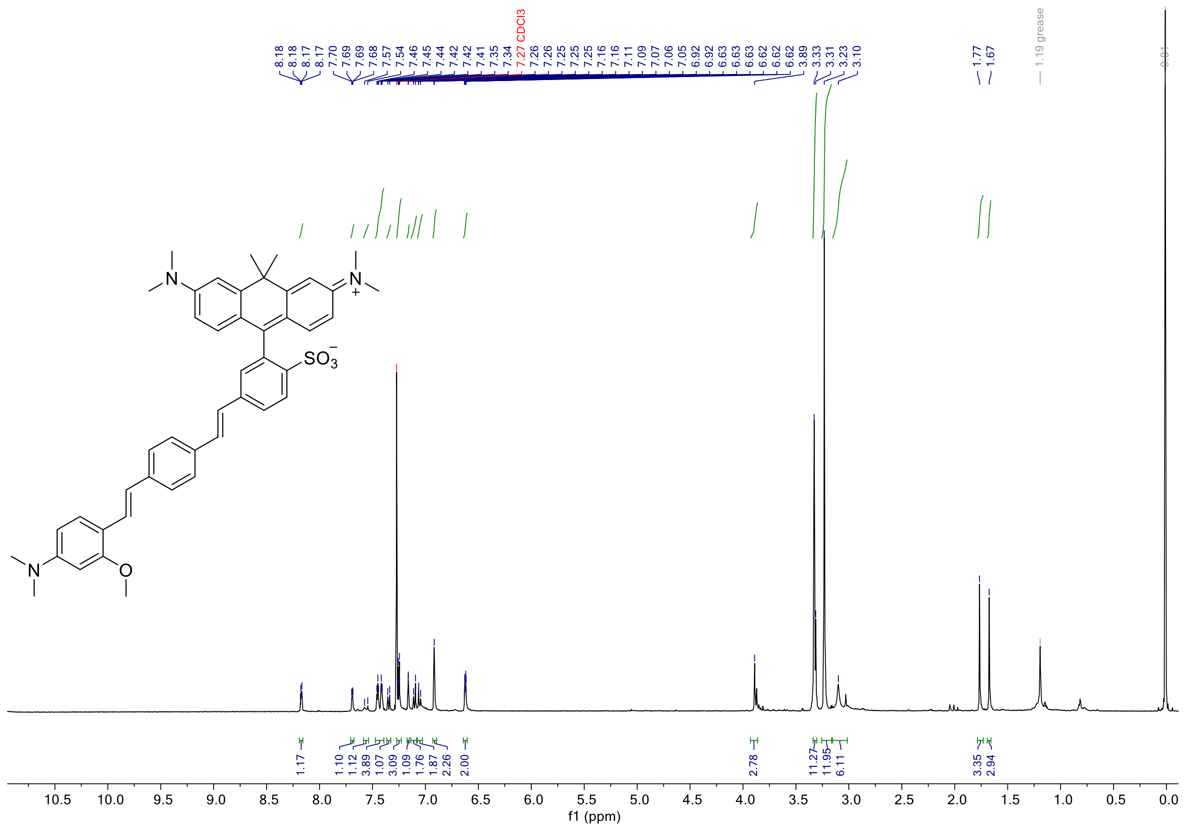

**Spectrum S26.** $^{13}$C NMR of compound 17

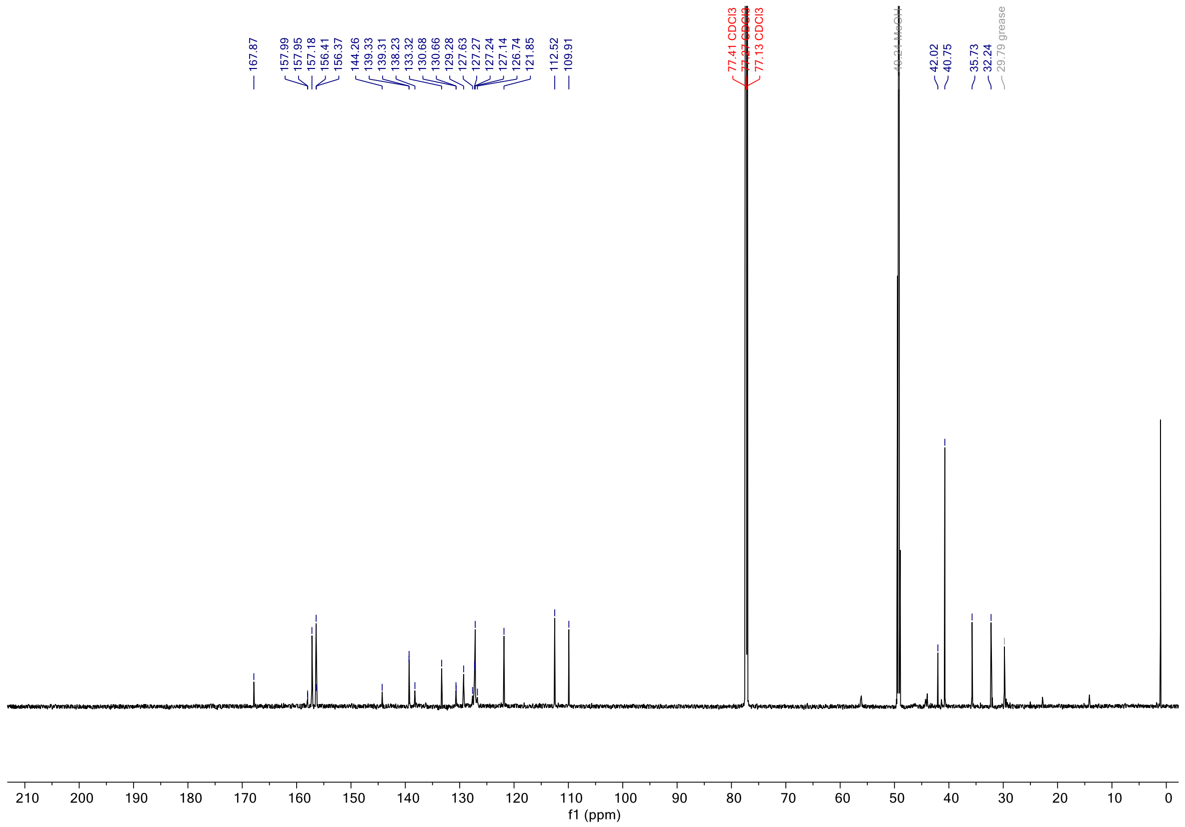

**Spectrum S27.** LCMS of compound 17

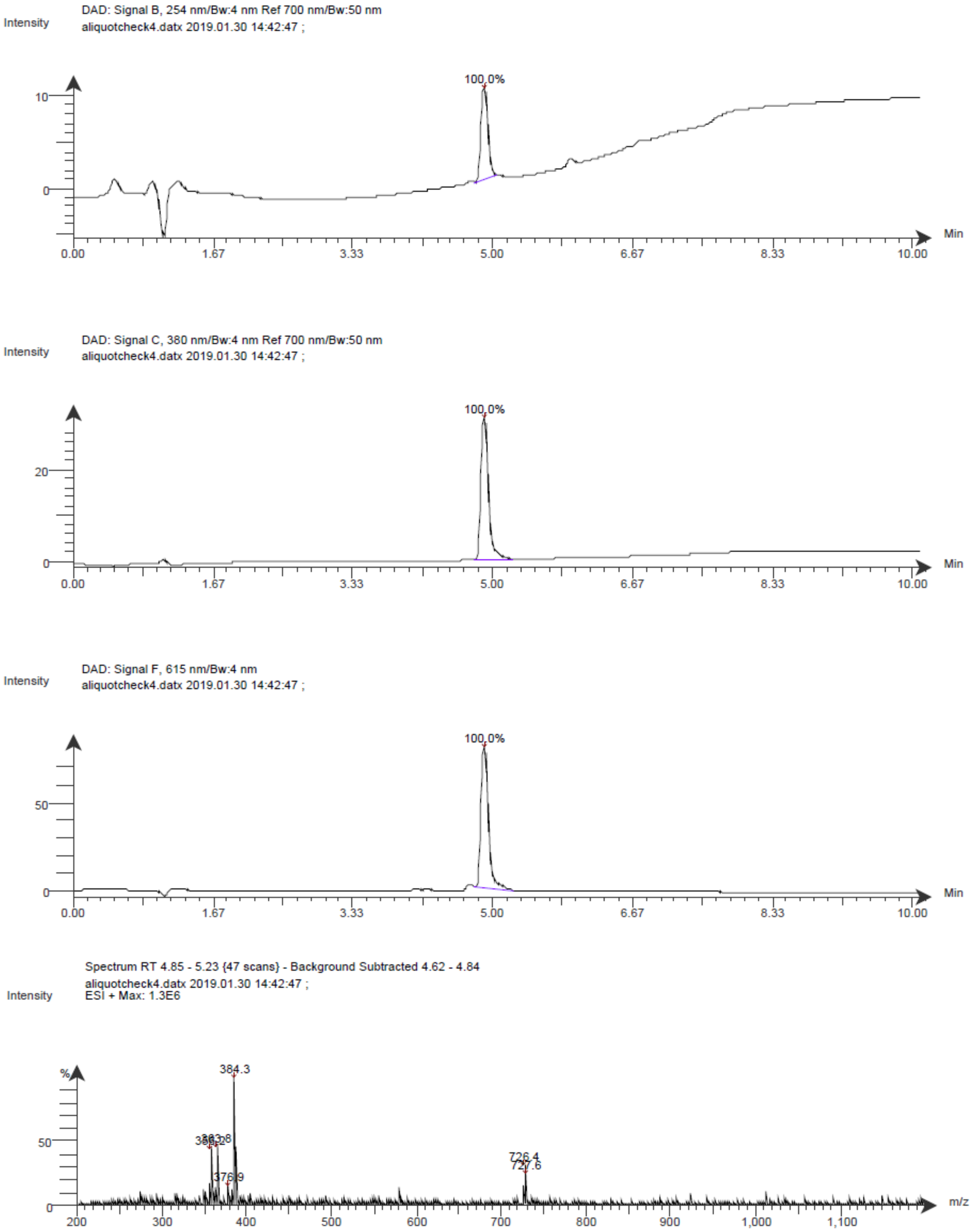
